# Qualitative and Quantitative Concentration-Response Modelling of Gene Co-expression Networks to Unlock Hepatotoxic Mechanisms for Next Generation Chemical Safety Assessment

**DOI:** 10.1101/2023.02.01.526628

**Authors:** Steven J. Kunnen, Emma Arnesdotter, Christian Tobias Willenbockel, Mathieu Vinken, Bob van de Water

**Affiliations:** Leiden University, Leiden Academic Centre for Drug Research, Division of Drug Discovery and Safety, Leiden, The Netherlands; Vrije Universiteit Brussel, Department of Pharmaceutical and Pharmacological Sciences, Brussels, Belgium; German Federal Institute for Risk Assessment (BfR), Department Pesticides Safety, Berlin, Germany

**Keywords:** benchmark concentration-response modelling, gene co-expression networks, next-generation risk assessment (NGRA), transcriptomic point of departure (tPOD), liver toxicity, hepatocyte

## Abstract

Next generation risk assessment of chemicals revolves around the use of mechanistic information without animal experimentation. In this regard, toxicogenomics has proven to be a useful tool to elucidate the underlying mechanisms of adverse effects of xenobiotics. In the present study, two widely used human *in vitro* hepatocyte culture systems, namely primary human hepatocytes (PHH) and human hepatoma HepaRG cells, were exposed to liver toxicants known to induce liver cholestasis, steatosis or necrosis. Benchmark concentration-response modelling was applied to transcriptomics gene co-expression networks (modules) in order to derive benchmark concentrations (BMCs) and to gain mechanistic insight into the hepatotoxic effects. BMCs derived by concentration-response modelling of gene co-expression modules recapitulated concentration-response modelling of individual genes. Although PHH and HepaRG cells showed overlap in deregulated genes and modules by the liver toxicants, PHH demonstrated a higher responsiveness, based on the lower BMCs of co-regulated gene modules. Such BMCs can be used as transcriptomics point of departure (tPOD) for assessing module-associated cellular (stress) pathways/processes. This approach identified clear tPODs of around maximum systemic concentration (C_max_) levels for the tested drugs, while for cosmetics ingredients the BMCs were 10-100 fold higher than the estimated plasma concentrations. This approach could serve next generation risk assessment practice to identify early responsive modules at low BMCs, that could be linked to key events in liver adverse outcome pathways. In turn, this can assist in delineating potential hazards of new test chemicals using *in vitro* systems and used in a risk assessment when BMCs are paired with chemical exposure assessment.

## Introduction

Safety testing of chemical compounds has historically been focusing on the hazard identification and as such largely relied on the assessment of apical endpoints in experimental animals (Hayes & Kruger, 2014). However, ethical and legislative constraints have instigated the development and application of animal-free methods, in particular *in vitro* systems, in regulatory chemical safety testing. Major advances have been made in next generation risk assessment (NGRA) using (non-animal) new approach methodologies (NAMs), that allow testing at the cellular and even molecular level, thereby facilitating a better understanding of the mechanisms leading to adverse effects, including the conceptualisation of adverse outcome pathways (AOP) (Choudhuri et al., 2018). Such mechanistic information enables more accurate prediction of biological responses and, in combination with exposure assessment, the risks associated with a defined exposure to a certain chemical compound.

The liver is a primary target organ for toxicity due to its pivotal role in the metabolism of xenobiotics (Gu & Manautou, 2012). It acts as a central regulator of lipid homeostasis, which is closely regulated by complex interactions between hormones, nuclear receptors, and transcription factors (Bechmann et al., 2012; Nguyen et al., 2008). Thus, drug- or chemical-induced injury to the liver may involve multiple and complex mechanisms (Jaeschke et al., 2012; Russmann et al., 2009; Vinken, Maes, et al., 2013). Changes in parameters related to hepatic steatosis (*i.e.* accumulation of fatty acids) (Ipsen et al., 2018) and cholestasis (*i.e.* accumulation of bile acids) (Chatterjee & Annaert, 2018) are two of the most frequent toxic manifestations seen in oral repeated dose toxicity data included in safety evaluation reports of cosmetic ingredients and in drug-induced liver injury cases (Gustafson et al., 2020; Kralj et al., 2021; Vinken et al., 2012). Additionally, hepatocellular injury/death may be one of the most relevant cellular key events (KEs) in various liver AOPs, which can be investigated using *in vitro* test systems for the prediction of hepatotoxicity (Arnesdotter et al., 2021). Currently, the gold standard *in vitro* test system remains cultures of primary human hepatocytes (PHH), but also the human hepatoma HepaRG cell line has been explored extensively for chemical safety testing, since it retains relatively high metabolic capacity (Andersson et al., 2012). However, despite the many advancements in cell culture technologies in recent years, replicating complex physiological processes and organ-specific functions *in vitro* (*i.e.* without the use of intact animals) is still a challenging task.

In the context of non-animal NGRA, toxicogenomics (TXG) plays a pivotal role in revolutionising the approach to evaluating chemical safety without relying on traditional animal testing methods. TXG allows comprehensive analysis of changes in cells, tissues and organisms at the molecular level, and is therefore a promising tool in mechanism-based risk assessment (Z. Liu et al., 2019). TXG can be used to detect a putative mechanism of action (MoA) (Berggren et al., 2017), substantiate disease mechanisms (AbdulHameed et al., 2019) or form the basis for defining or validating key events (KEs) in an AOP (Arnesdotter et al., 2022; Vinken, 2019). There is a growing interest to apply TXG to determine a transcriptomic point of departure (tPOD) for use in human health risk assessment (Farmahin et al., 2017; Friedman et al., 2020; Thomas et al., 2019; Thomas, Philbert, et al., 2013). However, to date, tPOD derivation from TXG data is not yet standard practice in regulatory risk assessment. Presently, TXG approaches predominantly focus on analysis of differentially expressed genes (DEGs) or enrichment analysis tools using gene annotations (Barel & Herwig, 2018). These methods depend on ontologies with a high degree of redundancy, which can result in a bias towards well-annotated genes and result in a flawed interpretation of mechanisms of toxicity applied to specific test systems (Callegaro et al., 2021; Vahle et al., 2018). Moreover, there is currently no consensus regarding the selection of individual genes for the prediction of adverse effects (Farmahin et al., 2017). Instead, gene co-expression analysis can be used to identify sets of genes expressed downstream of a (stress-responsive) common control mechanism, such as transcription factors. In the context of NGRA, analysis of co-expressed gene sets may provide mechanistic insights into observed adverse effects following exposure to toxicants or specific patterns of drug toxicity (Callegaro et al., 2021; Podtelezhnikov et al., 2020; Sutherland et al., 2018; Yin et al., 2021) and possibly serve as a solid basis for PoD derivation. Importantly, reproducibility between gene expression is higher when data are compared on the pathway level rather than at the gene level (Fan et al., 2010; Guo et al., 2006; Wang et al., 2014), thereby rendering co-expression networks better suited for inclusion in risk assessment (Sutherland et al., 2016). Several methods have been developed to identify and quantify co-expression networks, of which weighted gene co-expression analysis (WGCNA) is often applied and used for the development of the TXG-MAPr tools for mechanistic analysis of TXG data (Callegaro et al., 2021). The module eigengene score (EGS) provides a quantitative measure of the activity of a gene co-expression network (module), based on the log2 fold change expression of the module genes as described previously (Sutherland et al., 2016, 2018).

The aim of the present study was to systematically compare the temporal transcriptional responses in collagen sandwich cultures of PHH and HepaRG cells cultured on collagen in a conventional monolayer. These liver-based *in vitro* systems were exposed to chemical substances known and/or suspected to induce selected liver adverse effects (*i.e.* steatosis, cholestasis or necrosis). Benchmark concentration (BMC) modelling was applied on individual genes as well as gene co-expression networks from the PHH TXG-MAPr tool to derive *in vitro* transcriptomics benchmark concentrations (BMCs) and assess their suitability to be used as tPOD in chemical risk assessment.

## Materials and methods

### Cell cultures

PHH (10-donor, LIVERPOOL Cryoplateable Hepatocytes (5 male and 5 female, age 0 till 70 years); BioIVT, #X008001-P, LOT: KCB) were thawed in 25 mL Sekisui XenoTech OptiThaw Hepatocyte Kit (Tebu-bio, #K8000) and centrifuged at 100 x *g* for 10 minutes. Cell pellet was resuspended in INVITROGRO CP medium (BioIVT, #Z99029) supplemented with TORPEDO antibiotics mix (BioIVT, #Z990007) and seeded at a density of 70,000 cells per well in BioCoat Collagen-I-coated 96-wells plates (Corning, #354407). After 6 hours, PHH were washed with Dulbecco’s phosphate-buffered saline (DPBS) to remove dead cells. PHH were overlaid with 100 µl 0.25 mg/mL cooled Matrigel™ (BD Biosciences, #354230) in INVITROGRO HI medium (BioIVT, #Z99009) supplemented with TORPEDO antibiotics mix (BioIVT, #Z990007) to create a collagen-Matrigel™ sandwich. The next day, 24 hours after plating, PHH were ready to use for chemical exposures.

Cryopreserved differentiated HepaRG® cells (Biopredic International, #HPR116-TA08) were thawed and seeded (day 1) according to the manufacturer’s protocol. In short, 72,000 viable cells per well were seeded in collagen-coated (Corning #354236, 0.1 mg/ml in 0.02N acetic acid) 96-well plates (Falcon® #353072) in 100 µL basal hepatic cell medium (Biopredic International, #32551) supplemented with HepaRG® Thawing/Plating/General Purpose Medium (Biopredic International, #ADD670). On the 2^nd^ day of culture, the medium was changed to 100 µL basal hepatic cell medium with HepaRG® Maintenance/Metabolism Medium (Biopredic International, #ADD620) and subsequently renewed on days 4 and 7. A total of six different batches of HepaRG cells were used, three batches for drugs and three for cosmetic ingredients (HPR116-295; HPR116-294; HPR116-250; HPR116-301; HPR116-284; HPR116-291; Biopredic International).

### Chemicals and exposure

PHH were exposed to the compounds in 100 µL INVITROGRO HI medium (BioIVT, #Z99009) supplemented with TORPEDO antibiotics mix (BioIVT, #Z990007) 24 hours after plating. On day 8 in culture, HepaRG® cells were exposed to the compounds in 100 µL basal hepatic cell medium with serum-free HepaRG® Induction Medium (Biopredic International, #ADD650). Compound information and exposure concentrations were based on a range around the (estimated) C_max_ (defined as maximum chemical concentration in the blood after administration) of the chemicals **(Table S1)**. For the cosmetics ingredients, the estimated C_max_ values were calculated based on the systemic absorption described in reports issued by the Scientific Committee on Consumer Safety (SCCS) and assuming 100% bioavailability without any metabolism or clearance (SCCS, 2009, 2010, 2021). These assumptions are conservative by providing a maximum estimate of the systemic concentration when C_max_ values are not available. Cell lysates were collected at four different time points after exposure (*i.e.* 8, 24, 48 or 72 hours) (**Fig. 1a**). Cell cultures exposed for 48 and 72 hours were subjected to daily renewal of cell culture media, including the chemical compounds. Medium of PHH exposures was collected at every refreshment step and just before sample collection for RNA sequencing to determine the cytotoxicity using LDH assay. Briefly, 50 µL medium was collected in v-bottom plates, centrifuged for 5 min at 1000 rpm and stored at 4 °C until measurement for LDH activity according to the manufacturers protocol (Roche, LDH Cytotoxicity Detection Kit; 11644793001). For repeated exposures the cumulative cytotoxicity was calculated since medium replacement would wash away any released LDH. Stock solutions of butylated hydroxytoluene (BHT) (Sigma-Aldrich, #B1378), 2,7-naphthalediol (NPT) (Sigma-Aldrich, #8208510100), cyclosporine A (CSA) (Sigma-Aldrich, #C30024), and triclosan (TCS) (Sigma-Aldrich, #93453) were made in dimethyl sulfoxide (DMSO) (Sigma-Aldrich, #D8418) and stored at 4°C or -20°C. Fresh stock solutions of valproic acid (VPA) (Sigma-Aldrich, #P4543) and acetaminophen (APAP) (Sigma-Aldrich, #A3035) were made daily in cell culture medium. The final solutions were prepared *ex tempore* by diluting the stock solutions with cell culture medium. During exposure to TCS, BHT and NPT, plates were covered with QuickSeal Gas Perm^TM^ film (IST Scientific, #124-080SS) to prevent evaporation. All exposures were repeated three times on separate days, which serve as three independent biological replicates.

**Fig. 1.**
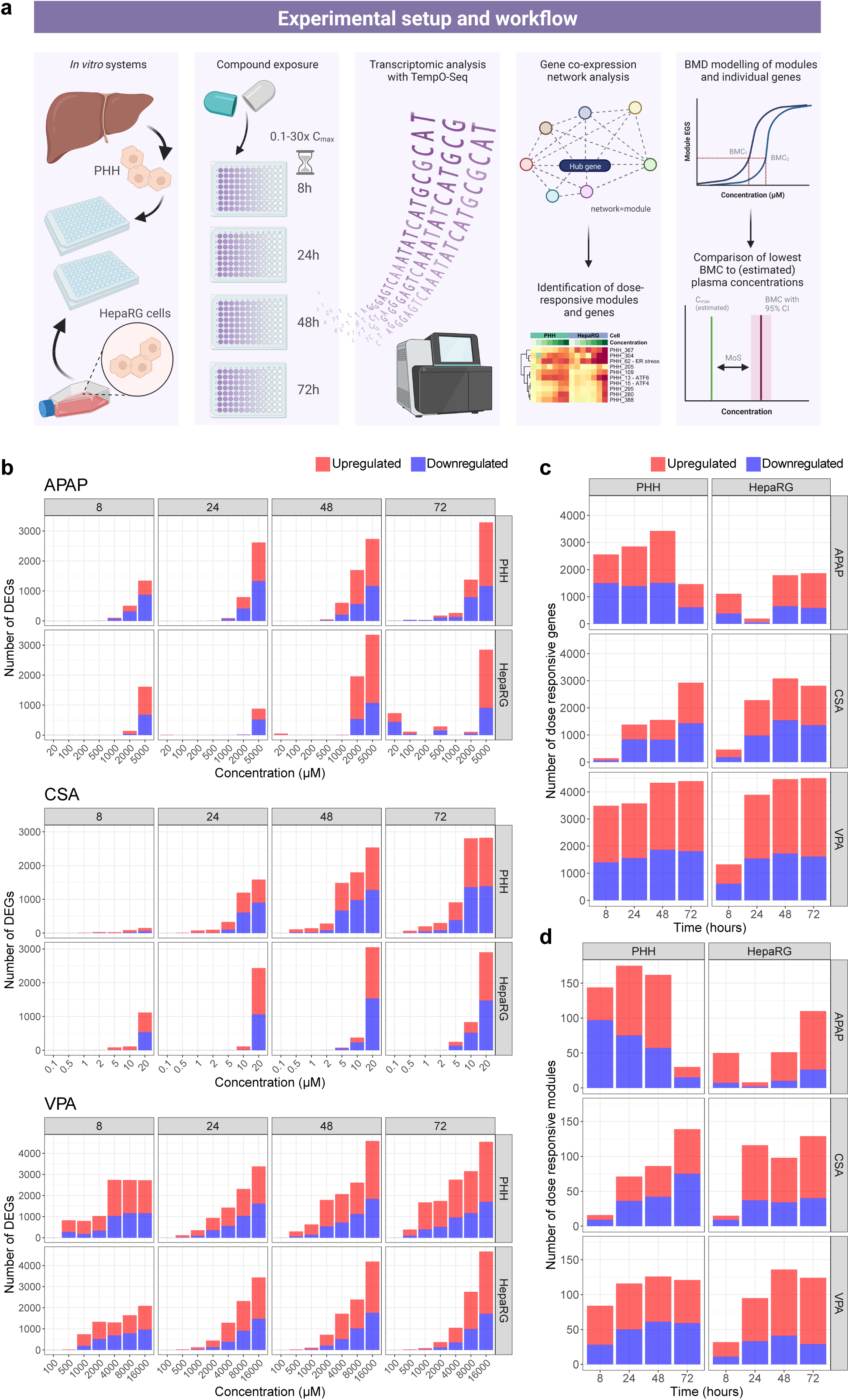
Experimental setup of transcriptomic analysis following chemical exposures in PHH and HepaRG cells. (A) Experimental setup of the study describing different steps of sample treatment, data analysis, concentration-response modelling and comparison of benchmark concentrations (BMCs) with (estimated) plasma concentrations (C_max_) to evaluate the margin of safety (MoS). Created with <u>BioRender.com</u>. (B) Differential gene expression analysis revealed a clear concentration and time dependency for all three drugs in PHH and HepaRG cells. Number of genes (C) and modules (D) showing a significant concentration-response using the William’s Trend Test (WTT). Differential gene expression and concentration responses were considered statistically significant after Benjamini-Hochberg multiple testing correction (p-adjust < 0.05).

### RNA sample preparation and sequencing

Following exposure, cell cultures were washed once with DPBS without calcium and magnesium chloride (Sigma-Aldrich, #D8537). Thereafter, cells were lysed with 50 µL lysis buffer (1:1 DPBS and 2x TempO-Seq lysis buffer, BioSpyder, mixed *ex tempore*) for 15 minutes at room temperature and stored at -80°C. Lysates were thawed and transferred to a 96-well conical bottom plate (#249662, Thermo Fischer) and plates were sealed with an aluminium silver seal (Greiner Bio-One, #676090). Lysates stored at -80°C were analysed at BioClavis (Glasgow, UK) using TempO-Seq targeted RNA sequencing technology with the human whole-transcriptome probeset version 2.0 according to their standard protocol (Yeakley et al., 2017). Raw sequencing data, metadata and processed data is deposited in the EMBL-EBI BioStudies genomics database ArrayExpress (E-MTAB-12668 and E-MTAB-12677).

### Data analysis

A count table of the RNA sequencing reads was provided by BioClavis, containing 1152 samples (2 cell types, 6 compounds, 7 concentrations + 1 control, 4 timepoints, 3 replicates) with 22537 measured probes. Samples with less than 500,000 counts were removed as these were outliers from the average of 3,000,000 reads per sample resulting in 1146 samples. Samples were normalised using the DESeq2 package (version 1.36.0) in R (version 4.1.0 or newer) by applying a counts per million (CPM) normalisation (Love et al., 2014; R Core Team, 2022). Low expressed probes (5247 probes in total with CPM < 1 in all treatment conditions) were removed from the raw expression matrix according to the relevance filter of the RNA-seq R-ODAF pipeline (Verheijen et al., 2022). Another 64 probes that showed the highest replicate variability were removed, which was 0.283% of the total probeset and equal to the percentage of PCA variance that showed highest replicate contribution. Next, multiple probes for the same gene were combined by taking the sum of the probe counts resulting in relevant count levels of 15156 unique genes. CPM normalisation was applied again by dividing raw counts by the sizefactors of each sample in the filtered raw expression matrix (1146 samples x 15156 genes) using the DESeq2 package, followed by differential gene expression analysis. For each chemical and time point, a model was built between the treatment concentrations and the time-matched vehicle control.

Significant gene expression was considered for p-adjust < 0.05 after Benjamini-Hochberg multiple testing correction. Log2 fold change (log2FC) threshold was not applied to allow for consideration of all significant fold changes (even small) and comparison of changes in gene expression and module activity. Modules eigengenes scores (EGS) were calculated using the PHH TXG-MAPr tool as described previously (Callegaro et al., 2021). Abs EGS > 2 were considered significant (Sutherland et al., 2018), although for some figures abs EGS > 5 were used as a cutoff to focus on only the highly perturbed modules. To calculate the module EGS per treatment replicate, the log2 fold change was determined per replicate, followed by the calculation of the module EGS using the replicate specific log2FC. BMDExpress (version 2.3) was used for benchmark dose (BMD) response modelling, also called benchmark concentration (BMC) modelling for *in vitro* experiments, on gene and module levels (Phillips et al., 2019). For gene level BMC modelling, the log2 normalised counts per replicate and treatment condition were used. The module EGS per replicate was used for module-level BMC modelling of each treatment condition. First, the William’s Trend Test (WTT) was performed using 10,000 permutations and Benjamini-Hochberg multiple testing correction to determine the significance of the concentration-response for all chemicals and time points. Thereafter, parametric BMC analysis was performed using continuous models (Hill, Power, Linear, Poly2, Exp2, Exp3, Exp4, Exp5) at 0.95 confidence level and benchmark response (BMR) factor of 1 SD for genes (*i.e.* recommended by the EPA for continuous data) and 1 or 2 SD for modules. The reason for deriving module BMCs at two different BMR factors is to compare with gene BMC (BMR factor of 1 SD) and to derive a BMC closer to a significant biological response at a module abs EGS > 2. Concentration-response models were considered significant with a p-adjust < 0.05 of the WTT. Best BMC model was selected based on the Nested Chi Square test, while flagged hill models were excluded and replaced by the next best model. The precision of the BMC calculation of the module EGS and the precision of the BMC calculation of single genes within the module was quantified with the BMDU/BMDL-ratio (also BMD or BMC precision factor), where the BMDU is the benchmark dose upper bound and the BMDL the benchmark dose lower bound of the BMC confidence interval (More et al., 2022). A BMC calculation with a lower BMDU/BMDL-ratio is considered to be more precise. A one-sided one-sample Wilcoxon signed rank test (R-package exactRankTests; p-value < 0.05) was used to test if the median of the BMDU/BMDL-ratios of all gene log2 CPM within a module was significantly higher than the BMDU/BMDL-ratio of the module EGS (Hothorn & Hornik, 2022). Modules with less than 5 genes were excluded from the analysis due to small sizes and lower biological relevance. To check the hypothesis that the BMC confidence intervals of the genes had a good overlap with the BMC confidence interval of the module EGS the base two logarithm of the ratio of overlap of all gene BMC confidence intervals was calculated with respect to the BMC confidence interval of the module, as described previously (Zoupa et al., 2020). When the base two logarithm was taken to the overlap ratio, a ratio of less than zero signalled a good overlap of BMC confidence intervals. It was tested if the median of the base two logarithm of the overlap ratios of the gene BMC confidence intervals with respect to the module BMC confidence interval was significantly lower than zero with a one-sided one-sample Wilcoxon signed rank test (R-package exactRankTests; p-value < 0.05). Overrepresentation analysis (ORA) was done on the concentration responsive genes (WTT p-adjust < 0.05) using the enrichR package (version 3.0) in R for the GO-terms, KEGG_2021_Human and WikiPathway_2021_Human (Kuleshov et al., 2016). All manuscript figures were created with R, and plotting packages including ggplot2, ggVennDiagram and pheatmap, unless indicated differently. Gene log2FC and BMC modelling data were also deposited in the EMBL-EBI BioStudies genomics database ArrayExpress (E-MTAB-12668 and E-MTAB-12677).

## Results

### Transcriptomics analysis of PHH and HepaRG cells exposed to liver toxicants

For the purpose of mechanism-based risk assessment, we investigated temporal concentration-dependent responses in cultures of PHH and HepaRG cells exposed to three drugs, acetaminophen (APAP), cyclosporine A (CSA) and valproic acid (VPA), that have a high liability for drug-induced liver injury but resulting in different adverse outcomes **(Fig. 1a)**. To facilitate the identification of early KEs and visualisation of their development over time, samples were collected at 4 time points, namely 8 and 24 hours (single exposure), 48 and 72 hours (daily repeated exposure), after exposure to a broad concentration range **(Table S1)**. Concentrations were selected based on the reported total C_max_ of each drug. As these are approved drugs currently on the market, they are not expected to induce overt adverse effects after single exposures around C_max_. Therefore, the selected maximum concentration was set at approximately 30x C_max_, whilst the minimum tested concentration was approximately 0.1x C_max_. There was no significant cytotoxicity observed in PHH for the tested drugs, although at the highest concentrations of CSA and VPA there was a trend of minimal (less than 10%) cytotoxicity (**Fig. S1**).

Differential gene expression analysis of the targeted RNA sequencing data (TempO-Seq) revealed a clear concentration and time dependency for all drugs **(Fig. 1b, Table S2)**. In general, significant alteration in gene expression was seen at lower concentrations in PHH compared to HepaRG cells, which may be indicative of higher responsiveness and/or sensitivity to toxicants. Concentration responses of individual genes were investigated by using the William’s Trend Test (WTT). A large number of genes showed a significant (p-adjust < 0.05) concentration-response at different timepoints **(Fig. 1c)**, which was comparable to the number of differentially expressed genes (DEGs) at the highest tested concentration. Overall, there was good overlap in DEGs and concentration responsive genes (CRGs) in PHH and HepaRG cells, mainly at later time points. Yet, considerable differences were observed in unique DEGs and CRGs for both test systems following exposure to APAP, CSA and VPA **(Fig. S2)**. Notably, most genes related to cytochrome P450 (CYP) iso-enzymes show different temporal expression patterns in vehicle-treated PHH and HepaRG cells, with typically higher expression levels in PHH and CYP expression declining for most high abundant CYP isoforms in HepaRG cells **(Fig. S3a)**. This indicates different xenobiotic metabolic capacity between the two liver test systems, which could influence their temporal response.

### Gene perturbation in PHH and HepaRG cells exposed to liver toxicants by gene co-expression networks

Mechanistic interpretation of transcriptomic data and comparison of transcriptional changes between test systems is complicated and time-consuming when a large number of genes exhibit significant differential expression or concentration-responses to controls, while gene set enrichment of DEGs is biased towards known biology (Barel & Herwig, 2018). Rather, the complexity of transcriptomic data can be reduced by analysing gene co-expressing network (module) activation (Sutherland et al., 2018). Therefore, we leveraged the recently published PHH TXG-MAPr tool for mechanism-based risk assessment of the transcriptomic data (Callegaro et al., 2021). Gene log2 fold changes of all treatment conditions were used to calculate module eigengene scores (EGS), which is a quantitative value for the activation or repression of the gene co-expression network **(Table S3)**. We used the module annotations and enrichments provided in the TXG-MAPr tool to gain mechanistic understanding of the module perturbations by the drugs. Using the WTT, numerous modules did exhibit a significant concentration-response **(Fig. 1d)**, which followed a similar pattern as the number of concentration-responsive genes. Obviously, the number of modules is much lower due to the clustering of co-expressed genes. A total of 192 concentration-responsive modules were strongly deregulated (p-adjust < 0.01, absolute EGS > 5) by at least one of the treatment conditions **(Fig. 2)**. These modules were clustered in 10 larger groups that reflected treatment specific responses.

**Fig. 2.**
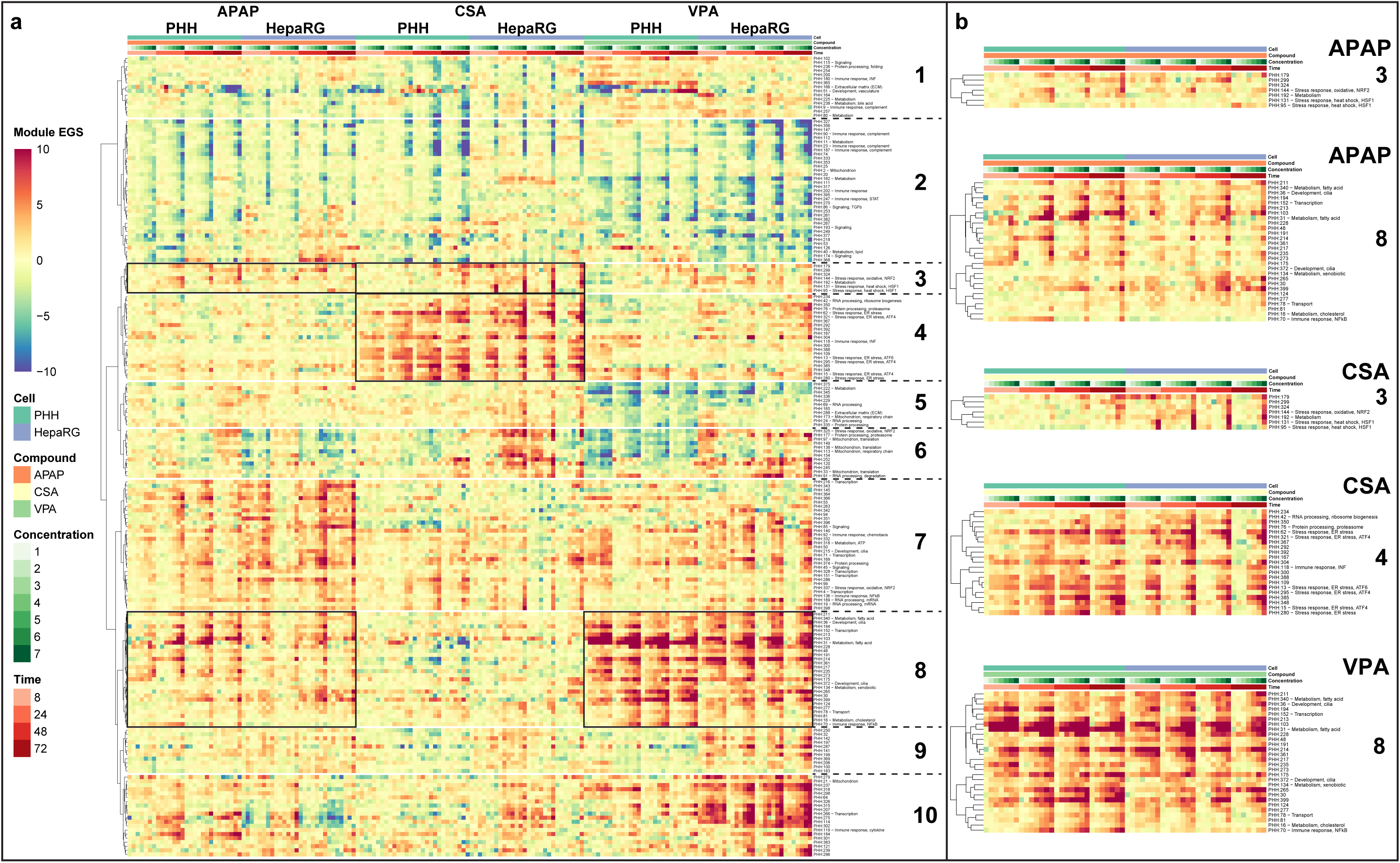
Module EGS heatmap of drug exposures in PHH and HepaRG cells. (A) Heatmap depicting concentration-responsive modules (WTT, p-adjust < 0.05) that were strongly deregulated (absolute EGS > 5) by at least one of the treatment conditions for the drugs. Ten clusters were defined based on Ward’s hierarchical clustering with correlation distance. (B) Specific clusters are highlighted for each drug (right). Module EGS is displayed in a colour gradient from 10 (red) to -10 (blue), although some modules have higher EGS, which are scaled to the most extreme colour.

The main cellular responses to treatment with liver toxicants could be clearly identified with the applied gene co-expression network approach, including well-known stress response activation by the three drugs in both PHH and HepaRG cells. As expected, APAP activates several stress-responsive modules in PHH and HepaRG, including module PHH:144 containing several nuclear factor erythroid 2–related factor 2 (NRF2) target genes, which is involved in an oxidative stress response **(Fig. 2, cluster 3)**. Additionally, fatty acid metabolism modules (PHH:31 and PHH:340) were induced by APAP **(Fig. 2, cluster 8)**. CSA induced several modules regulated by activating transcription factor 4 (ATF4; PHH:15, PHH:295 and PHH:321) and activating transcription factor 6 (ATF6; PHH:13), which are transcription factors involved in endoplasmic reticulum (ER) stress **(Fig. 2, cluster 4)**. Other stress-responsive modules activated by CSA included other ER stress-related (PHH:62 and PHH:280) and a proteasome (PHH:76) module **(Fig. 2, cluster 4)**, while the NRF2 module (PHH:144) and heat shock (HSF1) modules (PHH95 and PHH:131) were mainly activated at later timepoints and show stronger responses in HepaRG cells compared to PHH **(Fig. 2, cluster 3)**. Not surprisingly, modules annotated for fatty acid metabolism (PHH:31 and PHH:340) were strongly activated by VPA, as well as modules involved in xenobiotic metabolism (PHH:134), cholesterol metabolism (PHH:16), transport (PHH:78) and NFκB signalling amongst (PHH:70) other unannotated modules **(Fig. 2, cluster 8)**. All three drugs repressed the activity of a cluster of modules involved in immune responses and metabolism **(Fig. 2, cluster 2)**, which is often seen during cellular stress (Kültz, 2005). Generally, gene network modulation was concordant between PHH and HepaRG, with the latter showing activation at higher concentration (see cluster 3 and 8 for APAP responses and cluster 4 for CSA responses). Interestingly, while for VPA module activation was largely concordant between the two test systems, particular modules in clusters 9 and 10 demonstrated specific modulation by VPA mainly in HepaRG cells.

### Mechanism-based hazard assessment of liver toxicants using concentration-response modelling

To evaluate which genes and modules that are early responsive to liver toxicant treatment and thus most relevant to consider for risk assessment purposes, BMC modelling was performed using BMDExpress **(Tables S4, S5)**. Modules with highly significant concentration responses (WTT p-adjust < 0.01) were sorted by the lowest BMC using the response in PHH as the gold standard to visualise modules that are perturbed at the lowest compound concentration and may as such represent the most sensitive biological effects **(Fig. 3a, Table S6)**. The top five of the most important modules, in particular modules with low BMC and clear biological annotation, were selected per chemical **(Fig. 3a, black and red arrows)**. BMC model fitting of the main stress response modules for APAP, CSA and VPA showed different BMCs at the four tested time points **(Fig. 3b-d)**. APAP module PHH:144 demonstrated the lowest BMC at 24 hr, while for CSA and VPA the lowest BMC for modules PHH:13 and PHH:31, respectively, was observed at 72 hours. The selected stress responses are in line with the main pathways found using overrepresentation analysis (ORA) of differentially expressed genes, including NRF2 for APAP, ER stress or unfolded protein response (UPR) for CSA and lipid metabolism, including PPAR and PXR signalling for VPA **(Fig. S4)**. Concentration-response models on module level are highly representative of BMC models of the underlying genes for the three tested drugs **(Fig. S5)**. In addition, module-derived BMCs show a significant overlap in confidence interval or higher precision (low BMDU/BMDL ratio) compared to single gene-derive BMCs **(Fig. S6, Table S7)**, because the integration of gene co-expression in a weighted average module EGS value reduces experimental variability of individual genes. Accumulation plots of the module and gene level BMC revealed early and late responsive modules and genes with similar trends between the genes and modules **(Fig. 4)**. In general, although depending on the time point, the BMC of the concentration-responsive modules and genes seen in PHH were up to a factor 10 higher in HepaRG cells in particular for APAP and CSA exposure.

**Fig. 3.**
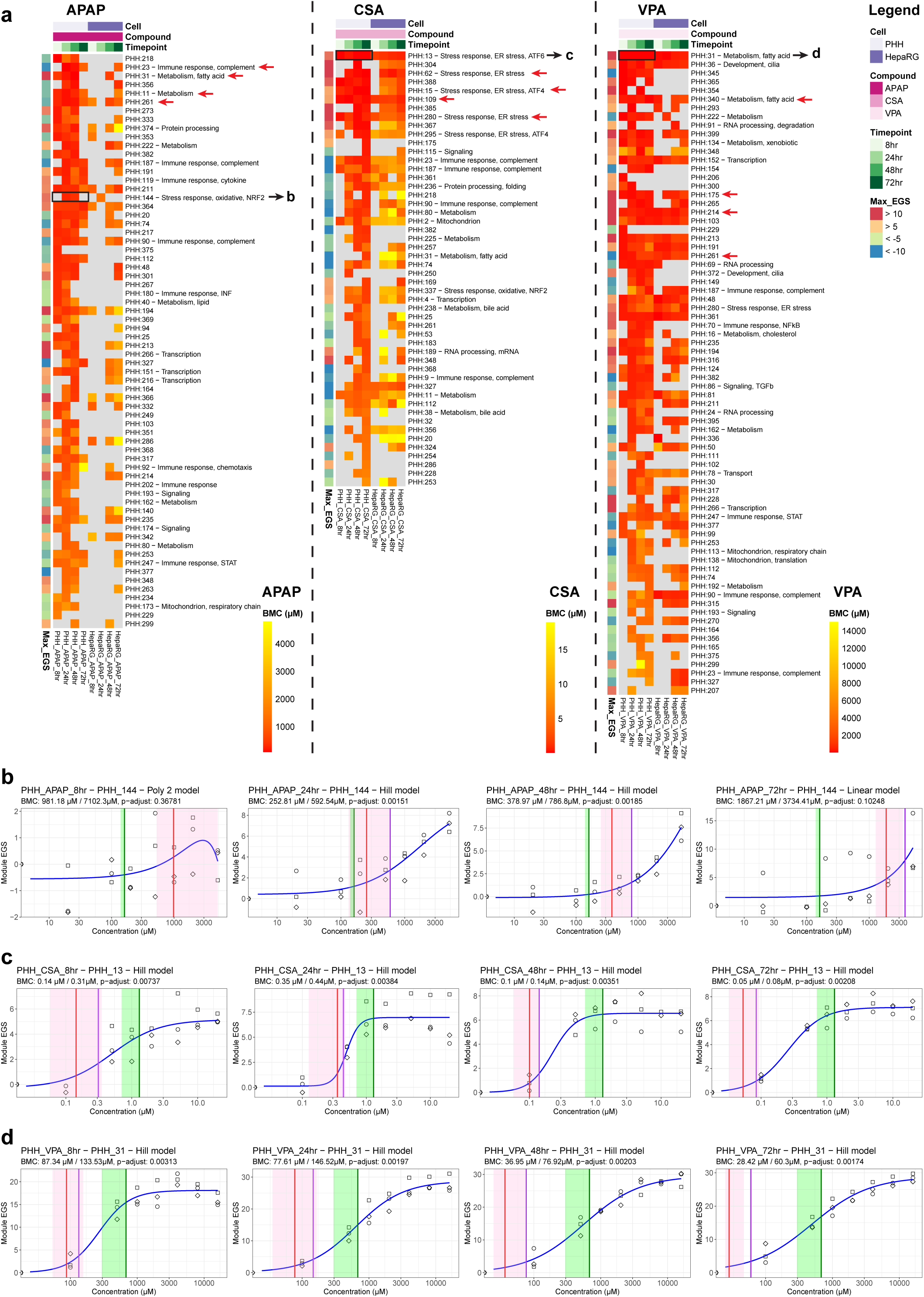
Benchmark concentration modelling of module eigengene score (EGS) responses following drug exposures in PHH and HepaRG cells. (A) Modules with highly significant concentration responses in PHH using the modules EGS (WTT, p-adjust < 0.01, absolute EGS > 5) were sorted by the lowest BMC using a BMR factor of 1 SD. The top five modules with the lowest BMC at most time points or a clear annotation are shown in Fig. 3B-D (black arrow) and Fig. S4 (red arrows). (B-D) BMC model fitting of the main stress response module per drug at the four tested time points. APAP induced the NRF2 module PHH:144 (B), CSA activated ATF6 module PHH:13 (C) and VPA induced fatty acid metabolism module PHH:31 (D). Blue lines display the model fitting of the best BMC model, as indicated in the subfigure’s title. The vertical green lines indicate the C_max_ values of the drugs. The vertical red and purple line indicate the BMC and determined by concentration response modelling using a BMR factor of 1 SD (red) or 2 SD (purple). The shaded pink area indicates the BMC confidence interval (BMDU - BMDL values) determined at BMR factor of 1 SD. The BMC value at BMR factor of 1 SD / 2 SD and the adjusted p-value of the WTT are shown in the subtitles of the figures.

**Fig. 4.**
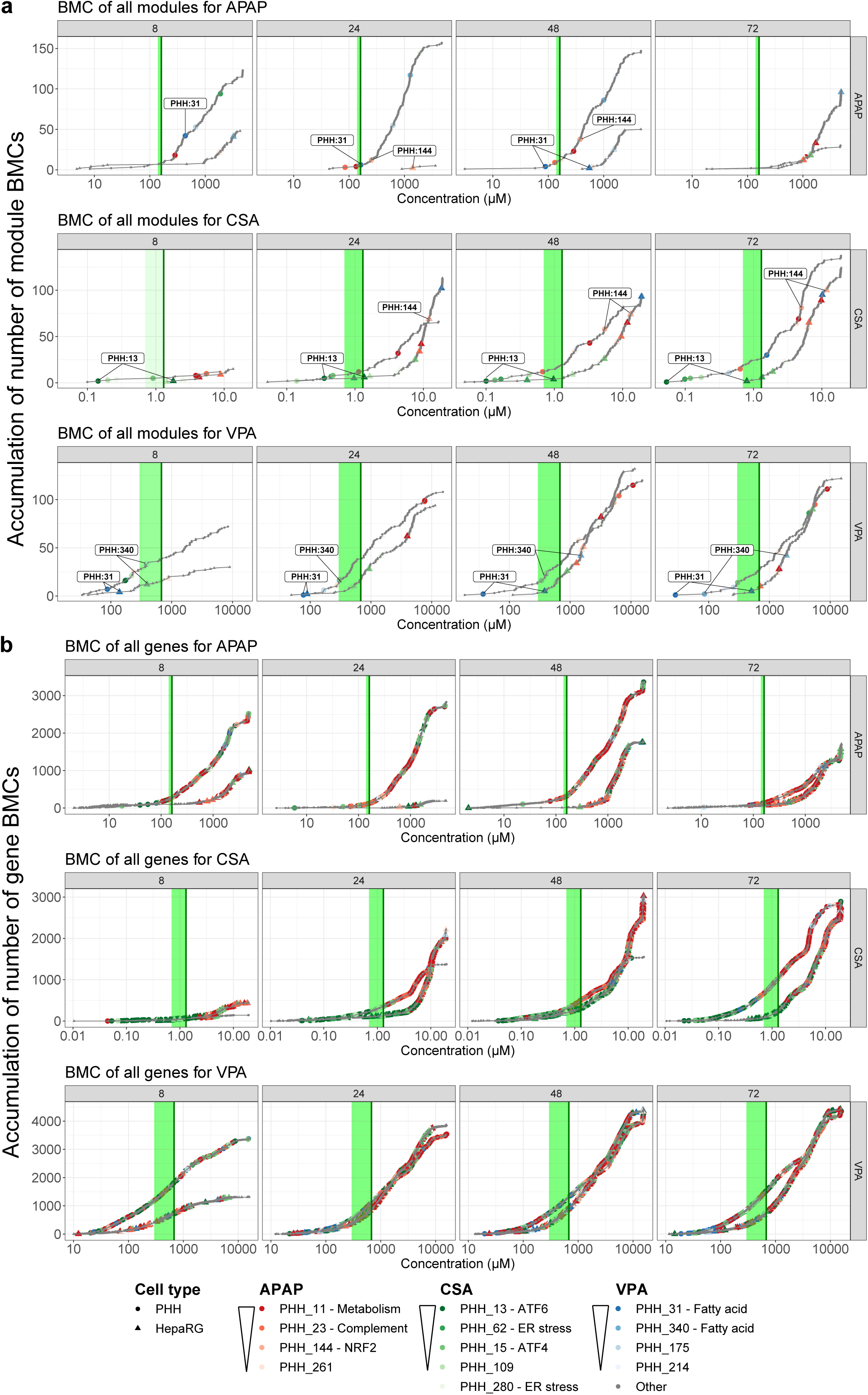
BMC accumulation plots of drugs. Accumulation plots of significant (WTT, p-adjust < 0.05) module (A) and gene level (B) BMCs, which clearly show early and late responsive modules and genes with similar trends between the genes and modules. Selected modules are highlighted with low BMCs following APAP (red colours), CSA (green colours) and VPA (blue colours) exposure.

### Acetaminophen

Acetaminophen (APAP) is a well-known inducer of centrilobular necrosis due to increased production of its deleterious reactive metabolite, known as *N*-acetyl-p-benzoquinone imine (NAPQI), following consumption of an overdose (James et al., 2008). Elevated levels of NAPQI cause rapid depletion of glutathione (GSH) in the liver, leaving free highly reactive NAPQI available to react with sulfhydryl groups to form APAP protein adducts, ultimately leading to oxidative stress and mitochondrial damage (McGill & Jaeschke, 2013; Ramachandran & Jaeschke, 2018). Therefore, the NRF2 module PHH:144 should be considered as the most mechanistically relevant concentration-responsive module. Indeed, PHH:144 showed a concentration-response at 24 and 48 hours following APAP treatment in PHH at a BMC between 250 and 600 µM, depending on the BMR factor, which is a threshold to determine the effect size of the response **(Fig. 3b)**. Thus, the derived BMC for NRF2 activation in PHH is about 2-4 times higher than the reported human C_max_ for APAP (i.e. 139 - 160 µM after ingestion of a standard 1000 mg dose (Farré et al., 2008; Sevilla-Tirado et al., 2003)). However, the BMC is in the range of a mild APAP overdose of 484 µM after taking a dose of approximately 73 mg/kg body weight, thus 50% of the adult toxic dose (Tan & Graudins, 2006). Moreover, in a high overdose situation (10 – 100 g APAP intake) the simulated C_max_ would be in the range of 1 - 2.7 mM (Spyker et al., 2022), which in this study clearly induced the NRF2 response in PHH. Hence, our data suggest that the BMC of the NRF2 response is in the range of a mild overdose and is increasing in activity during a high overdose.

No activation of the NRF2 response was seen at 8 hours, most likely due to the time required to reach glutathione depletion and the subsequent APAP protein adduct formation, which is a critical event in the hepatotoxicity. This is in line with another study that measured APAP protein adducts in APAP exposed HepaRG cell lysates. The authors showed that protein adduct peak levels were obtained already after 6 hours, albeit lactate dehydrogenase leakage did not increase dramatically until several hours later (Xie et al., 2015). Nevertheless, the PHH:144 module is activated 24 hours after APAP exposure in HepaRG cells, but at a higher BMC compared to PHH **(Fig. S5a)**. Well-known NRF2 target genes (*TXNRD1*, *SRXN1*, and *GCLM*) displayed similar concentration responses compared to the module as such and suggests limited bioactivation of APAP to reactive metabolites in HepaRG cells **(Fig. S5a)**. Indeed, lower levels of *CYP2E1,* the enzyme responsible for the metabolism of APAP to its reactive metabolite NAPQI, were observed in the HepaRG cells compared to PHH **(Fig. S3b)**. In addition, *CYP1A2*, equally responsible for metabolism of APAP, was expressed to a much lower extent in HepaRG cells compared to PHH. At 72 hours, the NRF2 concentration-response was still visible in PHH, but insignificant, which indicates attenuation of the adaptive NRF2 response.

It should be noted that module PHH:144 was not displaying the overall lowest BMC during APAP exposure. Multiple modules related to metabolism were perturbed at a lower BMC in APAP exposed cells, including the repressed modules PHH:11 and PHH:23 in PHH at 24 hours after APAP exposure **(Fig. S5a)**. Conversely, in HepaRG cells, the downregulation was non-significant on module and gene level at the early time points, although the trend seemed prominent. Additionally, module PHH:40, associated with biotin metabolism, was repressed while the fatty acid metabolism module PHH:31 was significantly induced in PHH at the earliest time point. Biotin is known to be involved in key metabolic pathways, such as gluconeogenesis, fatty acid synthesis, and amino acid catabolism (Pacheco-Alvarez et al., 2002). This indicates that APAP has a capacity to disturb lipid metabolism, which has been reported previously in mice (Chen et al., 2009). Module PHH:261 was most strongly downregulated at 48 hours both in PHH and HepaRG cells and contains calcium binding genes (*SCGN, S100A2*) **(Fig. S5a)**. Furthermore, numerous other modules showed significant concentration-responses following APAP exposure, including downregulated modules PHH:187 and PHH:90, involved in the complement pathway, and upregulated modules involved in transcription (PHH:151) or with unknown functions (PHH:374, 273, 222, 191, 211, 48, 301, 119, 217, 194, 364, 94, 213 etc.) **(Fig. 3a)**. Genes present in one of the selected modules for APAP with a low module BMC also display a low BMC on gene level **(Fig. 4, red colours)**, all indicating that gene responses show comparable BMCs as their respective module responses.

### Cyclosporine A

CSA is a well-known inducer of cholestasis (Sharanek et al., 2014, 2015; Tazuma, 2006). CSA has been shown to induce cholestasis through multiple mechanisms, including competitive inhibition of ATP-dependent bile salt transporters, especially the bile salt efflux pump (BSEP) and multidrug-resistant protein 2 (MRP2), by disorganisation of pericanalicular F-actin cytoskeleton (Böhme et al., 1994; Kadmon et al., 1993; Román & Coleman, 1994) and through obstruction of bile secretion by increasing canalicular membrane fluidity with no effect on the expression of canalicular transporters (Yasumiba et al., 2001). All but one (MRP2 inhibition) of these mechanisms are included in the AOP on cholestasis available on the AOP-Wiki (AOP ID 27), with BSEP inhibition being the molecular initiating event (Vinken, Landesmann, et al., 2013). Furthermore, CSA has been associated with both ER stress and oxidative stress (Hamon et al., 2014; Rao et al., 2018), which were both mentioned as KEs in an updated AOP on cholestasis (Gijbels et al., 2020). The results from the transcriptomics analysis showed a clear perturbation of multiple modules related to ER stress following exposure of both PHH and HepaRG cells to CSA. An ATF6 regulated module, PHH:13, was strongly induced while having one of the lowest BMC at all tested time points **(Fig. 3a, c and S5b)**. Similarly, another closely related ER stress module, PHH:62, displayed a significant concentration-response at 24 hours, as well as at later time points **(Fig. 3a and S5b)**. ATF4 related modules PHH:15 and PHH:295 showed the lowest BMC at the latest time points, as well as module PHH:280 containing the ATF4 target gene *DDIT3* (CHOP). Genes involved in ER stress or the UPR (*HSP90B, DNAJB9, DNAJB11, PDIA6, SELENOS, SELENOK, CHAC1, DDIT3, HSPA13* amongst others) showed similar concentration-responses and comparable BMCs as their respective modules **(Fig. S5b)**. Moreover, genes present in one of the ER stress related modules displayed the lowest BMC at all four time points, also suggesting that ER stress is the most relevant biological response after CSA exposure **(Fig. 4, green colours)**. The BMCs of ER stress-related modules were between 0.15 and 1.5 µM in PHH, and up to 10 times higher in HepaRG cells, which could be attributed to the higher metabolic clearances in HepaRG cells than in PHH (Bellwon et al., 2015). This was also supported by the higher initial expression of CYP3A4 / 5 / 7 in HepaRG cells **(Fig. S3b)**, which could result in a greater metabolic capacity of CSA, and leading to a lower level of CSA exposure, and thus lower gene expression changes compared to PHH. Interestingly, a module annotated as bile acid metabolism (PHH:38), were downregulated during CSA exposure at the later time points **(Fig. 3a)**, which may be an adaptive response to bile acid accumulation and commonly seen during cholestasis (Gijbels et al., 2019). Additionally, modules involved in metabolism (PHH:11, PHH:31, PHH:32), complement system (PHH:9, PHH:23, PHH:187, PHH:90) and mitochondrial function (PHH:2) were repressed following CSA exposure.

ER stress has been reported as one of the mechanisms of CSA-induced hepatocellular toxicity (Callegaro et al., 2021; Van den Hof et al., 2015), which could be confirmed in this study. However, oxidative stress has also been claimed to be one of the important stress pathways associated with CSA toxicity (Hamon et al., 2014). It may be that during ER stress, disrupted disulphide bond formation and breakage could lead to ROS accumulation and subsequently cause oxidative stress. Additionally, ER stress can cause mitochondrial dysfunction and thereby increase mitochondrial ROS production (Cao & Kaufman, 2014). ER stress and oxidative stress are not mutually exclusive processes, but can be subsequent processes during liver injury (Malhotra & Kaufman, 2007). Indeed, activation of NRF2-regulated oxidative stress (PHH:144 and PHH:337) was observed in the present study, but at higher CSA concentrations and later time points compared to ER stress **(Fig. 3a, 4a)**. So, ER stress is preceding oxidative stress following CSA exposure, which is in line with previous findings (Burban et al., 2018), but in contrast to another study (Gijbels et al., 2020). Nonetheless, the herein presented results identified ER stress as the primary mechanism of CSA induced toxicity, which may be a compound agnostic and early KE in the cholestasis AOP.

### Valproic acid

VPA is a commonly used steatogenic compound consisting of a branched-chain fatty acid that can compete with other fatty acids in hepatocyte metabolic pathways (Schumacher & Guo, 2015). VPA has the potential to inhibit mitochondrial fatty acid oxidation via three molecular initiating events (MIEs): (i) depletion of coenzyme A which is necessary for the oxidation of fatty acids, (ii) depletion of the biomolecule carnitine which transports fatty acids to mitochondria, and (iii) direct enzyme inhibition of β-oxidation (Allen et al., 2014), thereby hampering the β-oxidation of free fatty acids in the liver and causing accumulation of triglycerides in hepatocytes, known as steatosis (Pavlik et al., 2019). Altered fatty acid oxidation is included as a KEs in several described AOPs leading to steatosis in the AOP-Wiki (AOP IDs 318, 213, 232) and elsewhere (Mellor et al., 2016). In the present study, several modules displayed a low BMC at all tested time points of VPA exposure in both PHH and HepaRG cells, of which the fatty acid metabolism modules PHH:31 and PHH:340 were most significant and related to VPA induced steatosis **(Fig. 3, 4 and S5c)**. The BMC of module PHH:31 lowered with time and repeated dosing, suggesting an accumulative effect on fatty acid metabolism **(Fig. 3d)**. PHH:31 contains *ACSL6, ELOVL4* and *SLC27A1* that could be linked to fatty acid metabolism and peroxisome proliferator-activated receptors (PPAR) signalling, which all showed a significant concentration-response comparable to the module itself. The most significantly upregulated genes (*TMEM25 and TUBB2B*) were most similar to the concentration-response of PHH:31 but could, however, not be directly linked to fatty acid metabolism **(Fig. S5c)**. A recent study using the TXG-MAPr tool identified module PHH:31 as a common mechanism following exposure to carboxylic acids with steatotic potential, including VPA (Vrijenhoek et al., 2022). Moreover, VPA-induced module PHH:340 that contains two important genes involved in fatty acid oxidation and storage **(Fig. S5c)**. Specifically, *CPT1*, which encodes a key enzyme involved in the uptake and oxidation of long-chain fatty acids in the outer mitochondrial membrane (Begriche et al., 2011); and *PLIN2*, a gene involved in the coating of cytoplasmic lipid storage droplets (Itabe et al., 2017; Sztalryd & Kimmel, 2014). Other modules strongly activated by VPA in PHH and HepaRG cells, namely PHH:36, PHH:175, PHH:214, PHH:213, PHH:152, PHH:103, PHH:361, PHH:191, PHH:48, PHH:211 and PHH:194, were not clearly annotated in the TXG-MAPr, but correlated with fatty acid metabolism modules, which may suggest involvement in related processes **(Fig. 2 cluster 8, Fig. 3, Fig. S5c)**. Module PHH:261 was most strongly downregulated in both cell types **(Fig. S5c)** and contains calcium binding genes (*SCGN, S100A2*). Modules involved in the complement system (PHH:187, PHH:90) and mitochondrial function (PHH:113, PHH:138) were repressed during VPA exposure, amongst many other modules. *CYP2C9*, the main metabolising enzyme of VPA, showed comparable relatively high gene expression in the two cell types at the earlier time points **(Fig. S3b)**, which is in line with the small differences in transcriptional activation between the two cell systems.

### Characterisation of hazards of cosmetic ingredients in liver test systems using concentration-response modelling of quantitative gene networks

Most chemicals in cosmetic products pose little or no risk to human health. However, some chemicals have been linked to adverse effects (Panico et al., 2019). Therefore, we aimed to evaluate the applicability of the presented risk assessment approach and testing regimen to cosmetic ingredients that lack a defined pharmacological mode-of-action, yet have been, to some degree, associated with liver adversity (Gustafson et al., 2020; SCCS, 2021; Song et al., 2022). Necrosis and steatosis have been observed in the liver of experimental animals following exposure to butylated hydroxytoluene (BHT) (SCCS, 2021) and triclosan (TCS) (Song et al., 2022), respectively. In a review of animal toxicity data, 2,7-naphthalediol (NPT) was found to alter three parameters associated with, and thus possibly indicative of, cholestasis, namely increased serum levels of gamma-glutamyl transferase and bilirubin, together with cellular necrosis (Gustafson et al., 2020). Transcriptomics analysis was performed on PHH and HepaRG cell culture samples collected after exposure to NPT, BHT and TCS **(Table S1)**. There was cytotoxicity in NPT and TCS treated PHH starting at 24 hours and reaching up to 40% cytotoxicity at 48-72 hours (**Fig. S7**), while there was no cytotoxicity observed in PHH after BHT exposures. Differential gene expression analysis demonstrated only a clear concentration and time dependency in PHH following NPT exposure at sub-cytotoxic concentrations, while HepaRG cells had several DEGs mainly at the highest tested concentration **(Fig. S8a, Table S2)**. Numerous genes and modules showed a significant concentration response (WTT, p-adjust < 0.05) following NPT exposure in PHH and HepaRG cells, though PHH showed more gene perturbations at 48 and 72 hours **(Fig. S8b-c)**. Similarly, TCS exposure induced quite some DEGs at 48 hours, but that was the only time point at which there were several concentration-responsive genes and modules **(Fig. S8)**. Conversely, BHT did not induce any DEGs or cause activation of concentration-responsive genes and modules at any time point **(Fig. S8)**. Concentration responsive modules are clearly identified for PHH following NPT and TCS exposure, showing several clusters (8-10) with activated modules, while other clusters (2-4) are repressed **(Fig. S9, Table S3)**. However, due to the lack of concentration-responsive genes and modules in HepaRG cells, instead the data exhibit a mere noisy module EGS activation (data not shown).

BMC modelling identified a higher number of concentration responsive modules with low BMC after NPT, TSC and BHT exposure in PHH compared to HepaRG cells **(Fig. 5a, Tables S4-S6)**. It should be noted that different WTT p-values and max EGS thresholds had to be used for the three cosmetic compounds to facilitate the identification of modules with a concentration response trend for the least potent compound BHT. BMC model fitting of the lowest concentration-response modules for NPT, BHT and TCS identified different BMCs at the four tested time points, which were 10-100 fold higher than the estimated C_max_ values **(Fig. 5b-e)**. Not surprisingly, accumulation plots of the BMCs clearly show a steep increase in the number of BMCs at the highest tested concentrations of NPT and TCS **(Fig. 6)**. A similar trend could be observed for the BMCs of concentration-responsive genes. The transcriptomics results were compared to the response of hepatocytes to the drugs to elucidate any similarities in the MoA.

**Fig. 5.**
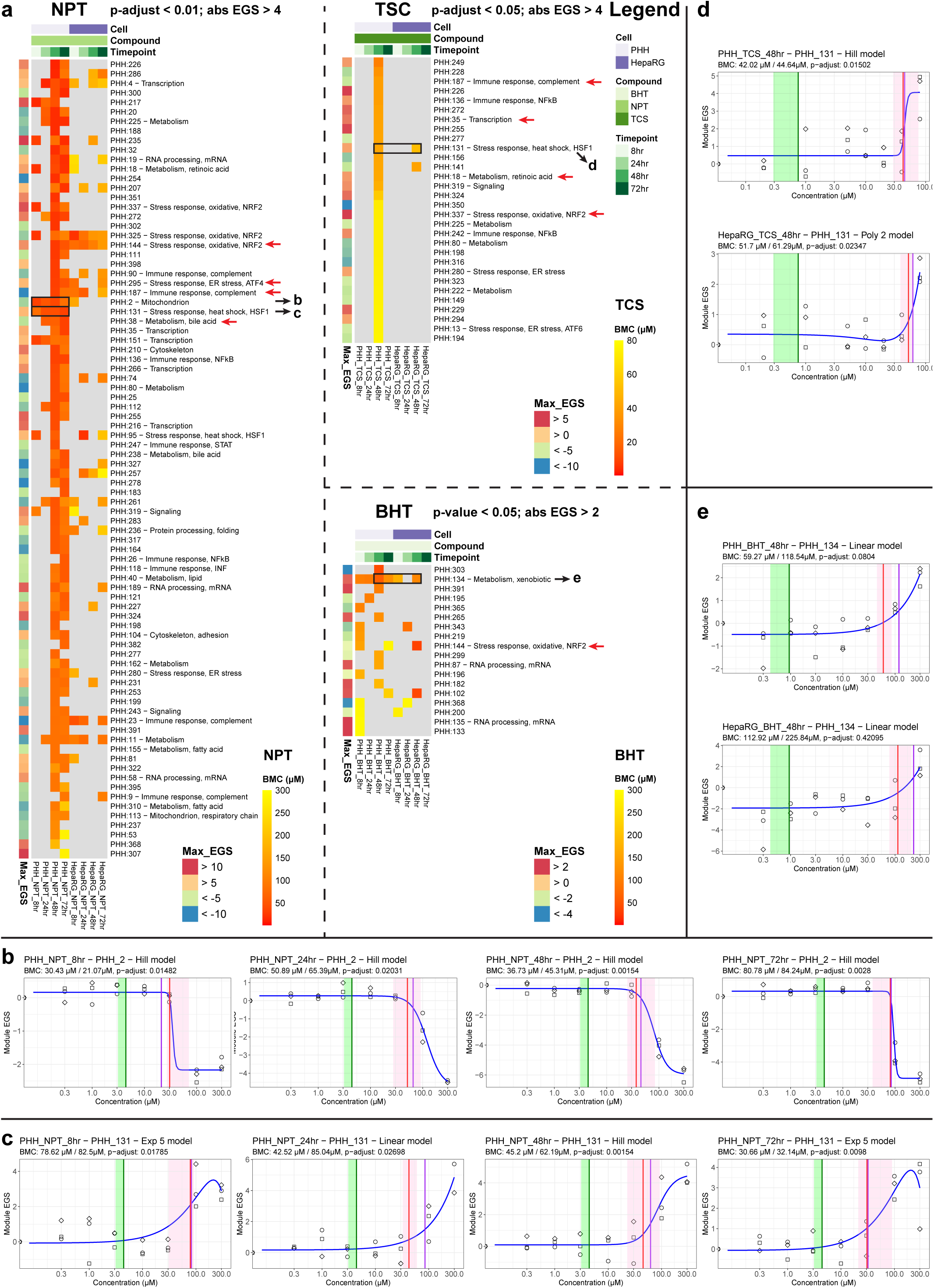
Benchmark concentration modelling of module EGS responses following cosmetic compound exposures in PHH and HepaRG cells. Heatmap depicting concentration-responsive modules using the modules EGS, that were strongly deregulated by at least one of the treatment conditions for the cosmetic compounds, sorted by lowest BMC in PHH using a BMR factor of 1 SD. Note that different WTT p-value and abs. EGS thresholds were used for the three cosmetic compounds. The top five modules with the lowest BMC at most time points or a clear annotation are shown in Fig. 5B-E (black arrows) and Fig. S8 (red arrows). (B-E) BMC model fitting of the repressed mitochondrial module (PHH:2) and activated heat shock module (PHH:131) following NPT exposure at the four tested time points. (D) BMC model fitting of activated heat shock module (PHH:131) following 48 hours TCS exposure in PHH and HepaRG cells. (E) BMC model fitting of activated xenobiotic metabolism module (PHH:134) following 48 hour BHT exposure in PHH and HepaRG cells. Blue lines display the model fitting of the best BMC model, as indicated in the subfigure’s title. The vertical green lines indicate the estimated C_max_ values of the cosmetic compounds. The vertical red and purple line indicate the BMC and determined by concentration response modelling using a BMR factor of 1 SD (red) or 2 SD (purple). The shaded pink area indicates the BMC confidence interval (BMDU - BMDL values) determined at BMR factor of 1 SD. The BMC value at BMR factor of 1 SD / 2 SD and the adjusted p-value of the WTT are shown in the subtitles of the figures.

**Fig. 6.**
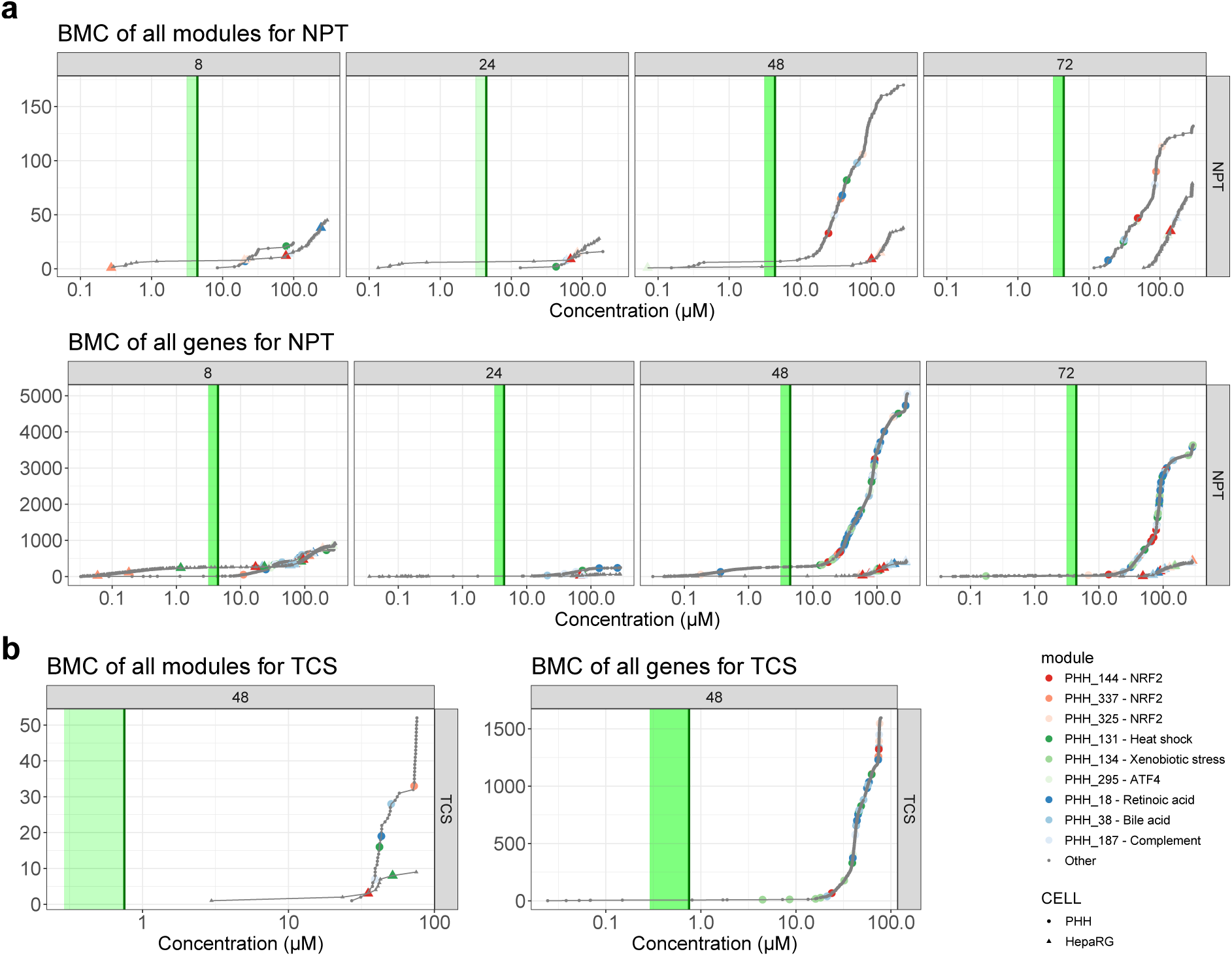
BMC accumulation plots of cosmetic compounds. Accumulation plots of significant (WTT, p-adjust < 0.05) module and gene level BMCs following NPT (A) and TCS (B) exposures in PHH and HepaRG cells. Selected modules are highlighted with NRF2 (red colours), protein folding (green colours) or other (blue colours) annotations.

### 2,7-naphthalenediol

NPT is used as a precursor for hair colours (SCCS, 2010) and has been identified as a potentially cholestasis-inducing compound in historical animal studies (Gustafson et al., 2020; Vinken et al., 2012). In this study, numerous genes and modules were deregulated in a concentration-responsive manner following NPT exposure, though several only upon repeated exposure at 48 and 72 hours **(Fig. 5a and 6)**. Early responsive modules in PHH include deactivation of mitochondrion (PHH:2) and activation of heat shock (PHH:131 and PHH:95), as well as two transcription regulation modules (PHH:4, PHH:151) and a retinoic acid metabolism module (PHH:18), all of which are still deregulated at the late time points **(Fig. 5a-c and S10a)**. Interestingly, in PHH a bile acid metabolism module (PHH:38) was downregulated during NPT exposure at the late time points, similar to CSA response, which may be an adaptive response to bile acid accumulation **(Fig. S10a)**. In contrast to CSA, the ER stress modules were hardly activated by NPT, but only ATF4 module PHH:295, suggesting a different MoA. Two complement system modules (PHH:187 and PHH:90) were repressed in PHH and HepaRG cells at late time points **(Fig. S10a)**. Multiple NRF2 modules (*i.e.* oxidative stress response; PHH:337, PHH:325 and PHH:144) were deregulated in both PHH and HepaRG cells **(Fig. 5a).** In HepaRG cells, the activation of PHH:144 started already at 8 hours, while PHH showed activation only after 48 hours **(Fig. S10a)**, which might suggest that early NRF2 activation in HepaRG cells is more cytoprotective (Baird & Dinkova-Kostova, 2011). These results are in line with the ORA of concentration-responsive genes, where several of these processes and pathways were also enriched **(Fig. S8d)**.

### Triclosan

TCS is a polychlorinated biphenolic antimicrobial used as an antiseptic and preservative in personal care products and medical equipment, which has been associated with liver steatosis (Vinken et al., 2012). Following exposure to TCS, more concentration-responsive modules were identified in PHH compared to HepaRG cells at 48 hours, of which a heat shock module (PHH:131) had the lowest BMC **(Fig. 5a, d)**. Other activated modules have been associated with transcription regulation (PHH:35), immune responses (PHH:136, PHH:242), retinoic acid receptors (PHH:18) and NRF2 response (PHH:337), while a complement system module (PHH:187) was repressed **(Fig. 5a, S10b)**. Notably, the fatty acid metabolism module PHH:340 (containing gene *CPT1* and *PLIN2*), that was clearly induced in VPA exposed cells was also slightly, albeit not significantly, induced. The low potency of TCS to induce transcriptional changes and the high (50-fold) difference between the estimated C_max_ and lowest BMC may suggest a low hazard of liver adversity.

### Butylated hydroxytoluene

BHT is a synthetic antioxidant and lipophilic organic compound. BHT is used across multiple chemical sectors and in numerous products, including food additives, personal/care products, pharmaceuticals, plastics/rubbers and other petroleum products. Upon metabolism by rat CYP2B1 (CYP2B6 in human), BHT can form reactive quinone methide metabolites, which has been associated with hepatocellular necrosis (SCCS, 2021). Unfortunately, the expression of *CYP2B6* was low (**Fig. S3**), which may explain the lack of transcriptional changes. Nevertheless, upregulation of CYP3A enzymes was observed, which complies with previously published data (Price et al., 2008). However, the role of these CYP3A enzymes in BHT metabolism or potential toxicity is unknown. Several modules were identified that showed a concentration-response trend (p-value < 0.05), albeit not significant after p-value adjustment **(Fig. 5a)**. The module with the lowest BMC at most of the time points has been associated with xenobiotic stress (PHH:134) and contains several CYP3A genes **(Fig. 5e and S10c)**. In addition, the NRF2 module PHH:144 (*i.e.* the most mechanistically relevant module for APAP exposure) also showed a slight induction at the highest BHT concentration at 8 hours **(Fig. S10c)**. This may indicate that NRF2 response can be induced by BHT, but only at higher concentrations, which could not be reached *in vitro* due to the limited solubility of BHT.

## Discussion

Current legislation in place in the European Union governing the safety of chemical compounds and cosmetic ingredients restricts and bans the use of animal testing, respectively (European Union, 2006, 2009). Consequently, the field of toxicology strives to develop and make use of methods that assesses the safety of chemicals without the use of experimental animals. In this regard, TXG is a powerful resource capable to inform on underlying mechanisms of possible adverse effects following exposure to exogenous compounds, and to derive toxicity values, such as tPODs, suitable for risk assessment. However, to our knowledge, there is currently no formal guidance on how genes, or groups of genes, should be selected for this purpose. With the growing interest in applying TXG to derive tPODs, exploration of best practices is direly needed. In this study, we present a method to derive BMCs from gene co-expression networks and demonstrate their similarity to BMCs derived from individual genes. We identified BMCs of modules that could be linked to the MoA of drugs after exposure in PHH and HepaRG cells and could be used as tPOD in risk assessment.

In this study, two human-relevant liver *in vitro* models were exposed to compounds with a well-known capacity of inducing common liver adversities, namely necrosis, cholestasis and steatosis. Subsequently, gene co-expression network analysis was combined with concentration-response modelling to gain knowledge of the mechanisms underlying these chemical-induced toxicities and to derive transcriptomics BMCs for such gene networks as well as for the individual genes. By following this strategy, we were able to readily get mechanistic understanding of the transcriptomic perturbation following liver toxicant exposures in PHH and HepaRG cells using module EGS from the PHH TXG-MAPr tool as a measure for the activity of the gene co-expression networks (Callegaro et al., 2021). Identified stress responses included an NRF2 response, ER stress and changes in fatty acid metabolism genes following exposure to APAP, CSA, and VPA, respectively, which were also reported previously (Grünig et al., 2020; Jaeschke et al., 2012; Rao et al., 2018; Xu et al., 2019). In addition to the gene co-expression (WGCNA) approach, these results were also confirmed by ‘traditional’ pathway enrichment analysis.

Subsequently, BMC modelling, which is advised by the European Food Safety Agency to be used over the traditional NOAEL approach (More et al., 2022) was used to derive time-point specific BMCs for gene co-expression modules, thereby demonstrating its capacity to recapitulate BMC modelling of individual genes. Overall, the BMC of modules were a good representation of the individual genes and showed less variability throughout the different timepoints. This suggests that pathway or module-derived BMCs are suitable for use in risk assessment, a finding which is in line with previous conclusions (J. Harrill et al., 2019). The BMC of modules had a clear tendency to be significantly more precise when measured by the BMDU/BMDL-ratio or their BMC confidence intervals exhibited a significant overlap with the BMC confidence intervals of the single genes **(Fig. S6)**. There was tendency for a trade-off between overlap and precision of the module and single gene BMC confidence intervals, which is to be expected because smaller or more precise BMC confidence intervals are less likely to exhibit an overlap. Another advantage of module-derived BMCs is the possibility to link modules to cellular processes or pathways, which could be used to directly investigate the MoA of a chemical or identify KEs in the context of liver adversity. Linking transcriptionally perturbed pathways or modules to KEs in an AOP context have been proposed previously (Saarimäki et al., 2023; Vinken, 2019). However, the impact of a possible KE activation on cellular physiological perturbations should be considered when determining the hazard linked to a chemical exposure, as some KEs may be more adverse than others, including DNA damage/mutation *versus* oxidative stress. This favours network-derived BMCs as, intuitively, the biological significance of the changes in a group of genes is greater than that of individual genes. In this context, BMCs of modules that represent KEs in a liver adversity could be used as tPOD for safety assessment. Nevertheless, it should be stressed that in NGRA it is not expected that a transcriptomic tPOD is used as a stand-alone method in a complete risk assessment, where also functional end points and exposure should be considered.

Following drug exposure, we could clearly identify mechanistically relevant gene networks with low BMCs, indicating that the tPOD of some important processes or KEs are around or below the C_max_ levels of the tested drugs. Increased reactive oxygen species or oxidative stress is a well-known KE in many AOPs leading to cell injury and death (Arnesdotter et al., 2021; Tanabe et al., 2022). Oxidative stress is also implicated in the mechanism of APAP induced hepatotoxicity (Yoon et al., 2016). Indeed, NRF2 module PHH:144 demonstrated a low BMC (using BMR factor of 2 SD) around 600 µM at 24 hr following APAP exposure, which was in the range of the plasma concentration following a mild APAP overdose, while NRF2 activity is increasing during a high overdose (Spyker et al., 2022; Tan & Graudins, 2006). In this case, the BMC derived at a BMR of 2 SD could be considered as a true biological response, since the BMC is often determined at a module abs EGS > 2, which is considered significant (Sutherland et al., 2018). The more conservative BMR factor of 1 SD resulted in a NRF2 module BMC of 250 µM, which is lower than a mild overdose, and thus could be used a tPOD for risk assessment. However, the module BMC is not as conservative as the BMC of 100 µM of the lowest gene *SRXN1*, which is lower than the C_max_ after a single APAP exposure **(Fig. S5a)**. This suggest that module derived BMCs using BMR factor of 1 SD are conservative enough to derive tPODs for safety assessment in case of APAP exposure, while focussing risk assessment on a small number of genes with low BMCs may introduce conservative and/or false positive hazards. Importantly, tPODs derived with different methods can differ several orders of magnitude, which exemplifies the need for reliable methods to derive tPODs (J. A. Harrill et al., 2024; Reardon et al., 2023). There is general scientific consensus that pathway based methods are preferred to determine tPOD, which will also provide mechanistic information that could be linked to KEs in AOPs (Barutcu et al., 2023; Basili et al., 2022; National Toxicology Program, 2018; Ramaiahgari et al., 2019; Thomas, Wesselkamper, et al., 2013), although others argue to determine tPOD as any concerted molecular change, as being more human health protective (Johnson et al., 2022). In this study, we presented such an approach to determine tPODs using modules as surrogate for pathways and/or KEs in AOP, but additional research will be needed to compare the reliability and robustness of this method compared to others.

For CSA and VPA, the lowest BMCs were observed at 72 hours and were both lower than their respective C_max_ levels, suggesting a significant human risk during CSA and VPA exposure for ER stress activation (PHH:13, PHH:15, PHH:62, PHH:295) and fatty acid metabolism (PHH:31 and PHH:340), respectively. ER stress or the unfolded protein response is also recognised as an important KE in the AOP of cholestasis according to recent literature (Burban et al., 2018; Gijbels et al., 2020; X. Liu et al., 2022; Selvaraj et al., 2020). Besides the ER stress response, other stress processes, like oxidative stress and bile acid metabolism, may contribute to cholestasis, but are likely more downstream KEs since these are induced at later time points and higher concentrations. Functional inhibition of the BSEP protein (*i.e.* the MIE of CSA induced cholestasis) can obviously not be measured at transcript level. Interestingly, a decrease of the gene (ABCB11) was seen at transcript levels following CSA exposure at high concentrations above C_max_ levels, which could potentially lead to lower BSEP levels and contribute to cholestasis. This is a counterintuitive response, as bile acids are known to induce ABCB11 expression (Vitale et al., 2023), although bile acids were not added to the medium and were not measured in the present study.

From an AOP perspective it is well-known that an imbalance in fatty acid metabolism, uptake or export can contribute to fatty acid accumulation and finally steatosis (Grefhorst et al., 2021). This is likely recapitulated on transcriptomic level by increased activity of fatty acid metabolism related modules PHH:31 and PHH:340 after VPA exposure, which may also be linked to mitochondrial toxicity (AbdulHameed et al., 2019). Interestingly, module PHH:31 is also induced by APAP, but with a lower BMC than the oxidative stress module, which could be a response to lipid peroxidation or mitochondrial damage, which are well-known KEs in APAP-induced liver adversity (Jaeschke & Ramachandran, 2018; Yoon et al., 2016).

In contrast to the drug exposures, for the cosmetic ingredients the BMCs were 10-100 fold higher than their estimated C_max_ levels, suggesting a reasonable margin of safety. For NPT there was a clear aberration of oxidative stress and bile acid metabolism modules at the late timepoints, which may be common downstream KEs in the cholestasis AOP as identified for CSA. In addition, NPT induced a heat shock response, which is possibly related to unfolded proteins, but NPT didn’t induce a similar ER stress response as CSA exposed PHH. It should be further stressed that the other two cosmetic compounds showed little correlation in the MoA with the drugs based on transcriptomic data. In fact, little gene perturbation was observed for TCS and BHT, even at very high sub-cytotoxic test concentrations. The low potency of cosmetic ingredients to induce transcriptional changes and the high (10-100 fold) difference between the estimated C_max_ and lowest BMC may suggest a much lower risk of liver adversity (Scientific Committee on Consumer Safety, 2021). Although the initial safety assessment can be guided by a transcriptional tPOD, it must be supplemented with other experimental approaches as validation (Rogiers et al., 2020). As this study only evaluated transcriptional changes and not any parameter for liver toxicity, like necrosis, triglyceride or bile acid accumulation, no final conclusions can be drawn in this regard. Thus, follow-up experiments at higher levels of biological organisation must be considered to get a more accurate prediction of the closely overlapping C_max_ and BMC and to confirm potential hazards for liver adversities of chemical exposure. For an oxidative stress response, such tests may include quantification of APAP-protein adducts, lipid peroxidation and oxidation of DNA (Dasgupta & Klein, 2014). For disturbance in fatty acid metabolism and transport (*i.e.* VPA) there are no less than 10 published AOPs related to steatosis in which numerous KEs are included that can be selected for further testing, such as intracellular lipid accumulation (Lichtenstein et al., 2020; Luckert et al., 2018; van Breda et al., 2018). For ER stress, such tests include western blot-based analysis of UPR-induced proteins and protein modifications (Kennedy et al., 2015) or more downstream bile acid accumulation as *in vitro* endpoint test for cholestasis.

The multifaceted role of the liver makes the selection of accurate toxicity testing procedures a daunting task. Notwithstanding multiple drawbacks, PHH are currently seen as the gold standard for *in vitro* hepatotoxicity testing. In any case, the availability and accuracy of the experimental model is of paramount importance to facilitate an ethic and scientifically sound risk assessment. In this light, inadequate metabolic competence in the model of choice can result in mischaracterisation of chemical hazard through both false positive (*i.e.* the compound is *de facto* detoxified *in vivo*) and false negative (*i.e.* the compound is bio-activated *in vivo*) results. The possibility of underestimating a hazard was recognised in the present paper by the (up to ten times) higher BMCs for the main stress responses in HepaRG cells compared to PHH. Multiple factors may contribute to the observed difference in activation and repression of genes, including the hepatocyte differentiation status, different expression of CYP enzymes as well as the fact that the terminally differentiated HepaRG cells used in these experiments have been shown to consist of a 1:1 mixture of hepatocyte-like and cholangiocyte-like cells (Guillouzo et al., 2007). Indeed, following exposure to CSA, the derived BMCs were higher in HepaRG cells compared to those from PHH. This particular difference could be attributed to a higher metabolic clearance in HepaRG cells compared to PHH (Bellwon et al., 2015). APAP exposure showed a difference in cellular activity between PHH and HepaRG as well, which could be explained by lower levels of *CYP1A2* and *CYP2E1,* the enzymes responsible for the metabolism of APAP, including its reactive metabolite NAPQI. HepaRG cells cultured in spheroid formation have shown higher sensitivity than traditional 2D cultures, thus in the future spheroid culture conditions could be considered to improve the performance of this system (Ramaiahgari et al., 2017).

The approach presented in this study allows fast identification of early responsive modules at low BMCs, which help to delineate the tPOD of cellular (stress) responses and potential hazards of chemicals using *in vitro* test systems. The *in vitro*-derived BMCs were compared to the estimated C_max_ levels, which, as expected, indicate much higher hazard for the tested drugs than the cosmetic ingredients. Finally, the *in vitro*-derived and module-based BMCs may be suitable for NGRA purposes, when combined with an *in vivo* dose extrapolation and exposure assessment.

## Abbreviations

AOP: Adverse outcome pathway
ATF4 / 6: Activating transcription factor 4 / 6
APAP: Acetaminophen
BHT: Butylated hydroxytoluene
BMC: Benchmark concentration
BMD: Benchmark dose
C_max_: Maximum concentration
CRGs: Concentration responsive genes
CSA: Cyclosporine A
CYP: Cytochrome P450
DEGs: Differentially expressed genes
EGS: Eigengene score
ER: Endoplasmic reticulum
FC: Fold change
KE: Key event
MIE: Molecular initiating event
MoA: Mechanism of action
MoS: Margin of safety
NAMs: New approach methodologies
NAPQI: N-acetyl-p-benzoquinone imine
NGRA: Next generation risk assessment
NPT: 2,7-naphthalenediol
NRF2: Nuclear factor erythroid 2–related factor 2
ORA: Overrepresentation analysis
PHH: Primary human hepatocytes [plural noun]
TCS: Triclosan
tPOD: Transcriptomic point of departure
TXG: Toxicogenomics
UPR: Unfolded protein response
VPA: Valproic acid
WTT: William’s Trend Test

## Acknowledgements

This project has received funding from Cosmetics Europe as part of the Long Range Science Strategy (LRSS) programme (project AIMT10), the European Chemical Industry Council (CEFIC), the EC Horizon2020 EUToxRisk project (grant number 681002), the EC Horizon2020 RISK-HUNT3R project (grant number 964537; part of the ASPIS cluster), the EC Horizon2020 ONTOX project (grant number 963845; part of the ASPIS cluster), the EC Horizon Europe PARC project (Partnership for the Assessment of Risk from Chemicals; grant number 101057014), the Fund for Scientific Research-Flanders, the University Hospital of the Vrije Universiteit Brussel-Belgium (“Willy Gepts Fonds” UZ-VUB), the Methusalem program of the Flemish government-Belgium, and the EU-EFPIA Innovative Medicines Initiative 2 (IMI2) Joint Undertaking under the TransQST project (grant number 116030). This Joint Undertaking receives support from the European Union’s Horizon 2020 research and innovation program and EFPIA.

The authors would like to acknowledge Gladys Ouédraogo (l’Oreal) and Catherine Mahony (Procter & Gamble) for their thoughtful insights in risk assessment of chemicals and cosmetic ingredients. The authors would like to thank Dr. Nynke Kramer, associate professor in toxicology at Wageningen University and Research, for the many insightful comments and guidance regarding C_max_ values and exposure extrapolation. The authors would like to thank Dr. Marije Niemeijer for her advice on primary human hepatocyte culture conditions and exposure. The authors acknowledge the technical staff at the Vrije Universiteit Brussel, in particular Ms. Dinja De Win, Mr. Steven Branson, Mr. Terry Desmae and Ms. Manon Wery for excellent assistance with cell culture experiments.

---

**Fig. S1. Cytotoxicity of drug exposures in PHH.** LDH cytotoxicity was determined at every sample collection timepoint by measuring LDH release in culture medium. Cumulative cytotoxicity is visualized in boxplots showing the median and quantiles (n=3). Significance of cytotoxicity was assessed using one-way Anova with concentration as independent variable (p < 0.05).

**Fig. S2. Venn diagrams of differential gene expression in PHH and HepaRG cells.** The overlap of the differentially expressed genes (DEGs) and concentration responsive genes (CRGs) in PHH and HepaRG cells are shown in a Venn diagram for the 3 drugs at the tested timepoints.

**Fig. S3. Basal CYP expression in PHH and HepaRG cells.** (A) Heatmap of the Cytochrome P450 (CYP) gene log2 normalized expression (CPM) at the four time points in the vehicle-treated samples. Treatment indicates for which compound this vehicle control was used. There was no apparent difference in basal CYP expression between the vehicle controls, although the was a clear drop in expression in time of some CYP genes, especially in HepaRG cells. (B) Gene log2 normalised expression of the CYP genes involved in the metabolism of the three drugs; APAP is metabolized by CYP1A2 and CYP2E1; CSA is metabolized by CYP3A4, CYP3A5 and CYP3A7; VPA is metabolised by CYP2C9; BHT is metabolized by CYP2B6.

**Fig. S4. Overrepresentation analysis of concentration response genes.** GO term and pathways (KEGG and WikiPathways) enrichment was performed using overrepresentation analysis (ORA) of the concentration responsive genes (WTT, p-adjust < 0.05) following APAP, CSA and VPA treatment. The top 5 most significant terms per timepoint were selected to give an overview of the most enriched GO-terms and pathway terms.

**Fig. S5. BMC modelling of drug-induced responses in PHH and HepaRG cells.** BMC model fitting of top five modules with lowest BMC at most time points and a clear annotation for each drug (left column), *i.e.* (A) APAP, (B) CSA and (C) VPA. BMC model fitting of the gene level BMC for the most significant gene (p-adjust and log2 FC) within each module (middle two columns). Overlap between module BMC (red line) and gene-level BMC (dot) and their confidence intervals, *i.e.* shaded area and black lines, respectively (right column). Blue lines display the model fitting of the best BMC model, as indicated in the subfigure’s title. Vertical lines and shaded area indicate the total C_max_ values (green) of the drugs and the BMCs (red or purple) + confidence interval (BMDU - BMDL values) determined by concentration response modelling using a BMR factor of 1 SD (red) or 2 SD (purple). The BMC value at BMR factor of 1 SD / 2 SD and the adjusted p-value of the WTT are shown in the subtitles of the figures. The lowest adjusted p-values of differentially expressed genes (DEG) is also shown for concentration responsive genes. In the module gene overlap figure the upper and lower confidence intervals of the module BMC are given.

**Fig. S6. Comparison between module and gene derived BMC confidence intervals.** Module EGS-derived BMCs show a significant overlap in confidence interval with gene log2FC derived BMCs, significant higher precision (low BMDU/BMDL ratio) or both significant overlap and higher precision compared to single gene-derive BMCs. Non-significant indicates that module EGS derived BMC didn’t show a significantly higher precision and didn’t show a good overlap with most genes in a module.

**Fig. S7. Cytotoxicity of cosmetic compound exposures in PHH.** LDH cytotoxicity was determined at every sample collection timepoint by measuring LDH release in culture medium. Cumulative cytotoxicity is visualized in boxplots showing the median and quantiles (n=3). Significance of cytotoxicity was assessed using one-way Anova with concentration as independent variable (p < 0.05).

**Fig. S8. Differential and concentration-responsive gene (co-)expression following cosmetic compound exposure in PHH and HepaRG cells.** (A) Number of differentially expressed genes in treatments following cosmetic compound exposures in PHH and HepaRG cells. Number of genes (B) and modules (C) showing a significant concentration-response using the William’s Trend Test (WTT). Differential gene expression and concentration responses were considered statistically significant after Benjamini Hochberg multiple testing correction (p-adjust < 0.05). (D) GO term and pathways (KEGG and WikiPathways) enrichment was performed using overrepresentation analysis (ORA) of the concentration responsive genes (WTT, p-adjust < 0.05) following NPT treatment. The top 5 most significant terms per timepoint were selected to give an overview of the most enriched GO-terms and pathway terms.

**Fig. S9. Module EGS heatmap of cosmetic exposures in PHH.** Heatmap depicting concentration-responsive modules (WTT, p-adjust < 0.05) that were strongly deregulated (absolute EGS > 4) by at least one of the treatment conditions for the cosmetic compounds in PHH. Ten clusters were defined based on Ward’s hierarchical clustering with correlation distance. Module EGS is displayed in a colour gradient from 10 (red) to -10 (blue), although some modules have higher EGS.

**Fig. S10. BMC modelling of cosmetics-induced responses in PHH and HepaRG cells.** BMC model fitting of the top modules (left column) with lowest BMC at most time points and a clear annotation for each cosmetic ingredient, *i.e.* (A) NPT, (B) TCS and (C) BHT. BMC model fitting of the gene level BMC for the most significant gene (p-adjust and log2 FC) within each module (middle two columns). Overlap between module BMC (red line) and gene-level BMC (dot) and their confidence intervals, *i.e.* shaded area and black lines, respectively (right column). Blue lines display the model fitting of the best BMC model, as indicated in the subfigure’s title. Vertical lines and shaded area indicate the estimated C_max_ values (green) of the cosmetic ingredients and the BMCs (red or purple) + confidence interval (BMDU - BMDL values) determined by concentration response modelling using a BMR factor of 1 SD (red) or 2 SD (purple). The BMC value at BMR factor of 1 SD / 2 SD and the adjusted p-value of the WTT are shown in the subtitles of the figures. The lowest adjusted p-values of differentially expressed genes (DEG) is also shown for concentration responsive genes. In the module gene overlap figure the upper and lower confidence intervals of the module BMC are given.

## Supplementary tables

**Table S1.** Compound information and experimental treatment conditions used for transcriptomic analysis in primary human hepatocytes (PHH) and HepaRG cells. Total C_max_ values for drugs were obtained from literature (see reference). Estimated C_max_ values were calculated for the cosmetic ingredients based on the absorption (see reference).

**Table S2.** Significant gene log2 fold changes of compound exposures in PHH and HepaRG cells.

**Table S3.** Module eigengene scores (EGS) of compound exposures in PHH and HepaRG cells.

**Table S4.** Significant gene-level BMC modelling output from BMDExpress (p < 0.05).

**Table S5.** Module-level BMC modelling output from BMDExpress.

**Table S6.** Benchmark concentrations of significant (p-adjust < 0.05) concentration responsive co-expression modules depicted in Fig. 3 and 5 derived from BMD modelling with BMR factor of 1 SD. Colours are based on the highest (estimated) Cmax values of the chemicals, where values < Cmax have orange/red colours, and values >= Cmax have yellow/green colours.

**Table S7.** Concentration-responsive module BMCs compared with gene-derived BMC confidence intervals. Module-derived BMCs show a significant overlap in confidence interval (Wilcoxon p-value, normal and log2 scale), significant higher precision (low BMDU/BMDL ratio) or both significant overlap and higher precision compared to single gene-derive BMCs.

## References

AbdulHameed, M. D. M., Pannala, V. R., & Wallqvist, A. (2019). Mining Public Toxicogenomic Data Reveals Insights and Challenges in Delineating Liver Steatosis Adverse Outcome Pathways. Frontiers in Genetics, 10. 10.3389/fgene.2019.01007

Allen, T. E. H., Goodman, J. M., Gutsell, S., & Russell, P. J. (2014). Defining Molecular Initiating Events in the Adverse Outcome Pathway Framework for Risk Assessment. Chemical Research in Toxicology, 27(12), 2100–2112. 10.1021/tx500345j

Andersson, T. B., Kanebratt, K. P., & Kenna, J. G. (2012). The HepaRG cell line: a unique in vitro tool for understanding drug metabolism and toxicology in human. Expert Opinion on Drug Metabolism & Toxicology, 8(7), 909–920. 10.1517/17425255.2012.685159

Arnesdotter, E., Gijbels, E., dos Santos Rodrigues, B., Vilas-Boas, V., & Vinken, M. (2022). Adverse Outcome Pathways as Versatile Tools in Liver Toxicity Testing (pp. 521–535). 10.1007/978-1-0716-1960-5_20

Arnesdotter, E., Spinu, N., Firman, J., Ebbrell, D., Cronin, M. T. D., Vanhaecke, T., & Vinken, M. (2021). Derivation, characterisation and analysis of an adverse outcome pathway network for human hepatotoxicity. Toxicology, 459, 152856. 10.1016/j.tox.2021.152856

Baird, L., & Dinkova-Kostova, A. T. (2011). The cytoprotective role of the Keap1–Nrf2 pathway. Archives of Toxicology, 85(4), 241–272. 10.1007/s00204-011-0674-5

Barel, G., & Herwig, R. (2018). Network and pathway analysis of toxicogenomics data. Frontiers in Genetics, 9(OCT). 10.3389/fgene.2018.00484

Barutcu, A. R., Black, M. B., & Nong, A. (2023). Mining toxicogenomic data for dose-responsive pathways: implications in advancing next-generation risk assessment. Frontiers in Toxicology, 5. 10.3389/ftox.2023.1272364

Basili, D., Reynolds, J., Houghton, J., Malcomber, S., Chambers, B., Liddell, M., Muller, I., White, A., Shah, I., Everett, L. J., Middleton, A., & Bender, A. (2022). Latent Variables Capture Pathway-Level Points of Departure in High-Throughput Toxicogenomic Data. Chemical Research in Toxicology, 35(4), 670–683. 10.1021/acs.chemrestox.1c00444

Bechmann, L. P., Hannivoort, R. A., Gerken, G., Hotamisligil, G. S., Trauner, M., & Canbay, A. (2012). The interaction of hepatic lipid and glucose metabolism in liver diseases. In Journal of Hepatology (Vol. 56, Issue 4). 10.1016/j.jhep.2011.08.025

Begriche, K., Massart, J., Robin, M.-A., Borgne-Sanchez, A., & Fromenty, B. (2011). Drug-induced toxicity on mitochondria and lipid metabolism: Mechanistic diversity and deleterious consequences for the liver. Journal of Hepatology, 54(4), 773–794. 10.1016/j.jhep.2010.11.006

Bellwon, P., Truisi, G. L., Bois, F. Y., Wilmes, A., Schmidt, T., Savary, C. C., Parmentier, C., Hewitt, P. G., Schmal, O., Josse, R., Richert, L., Guillouzo, A., Mueller, S. O., Jennings, P., Testai, E., & Dekant, W. (2015). Kinetics and dynamics of cyclosporine A in three hepatic cell culture systems. Toxicology in Vitro, 30(1). 10.1016/j.tiv.2015.07.016

Berggren, E., White, A., Ouedraogo, G., Paini, A., Richarz, A. N., Bois, F. Y., Exner, T., Leite, S., Grunsven, L. A. van, Worth, A., & Mahony, C. (2017). Ab initio chemical safety assessment: A workflow based on exposure considerations and non-animal methods. Computational Toxicology, 4, 31–44. 10.1016/j.comtox.2017.10.001

Böhme, M., Müller, M., Leier, I., Jedlitschky, G., & Keppler, D. (1994). Cholestasis caused by inhibition of the adenosine triphosphate-dependent bile salt transport in rat liver. Gastroenterology, 107(1), 255–265. 10.1016/0016-5085(94)90084-1

Burban, A., Sharanek, A., Guguen-Guillouzo, C., & Guillouzo, A. (2018). Endoplasmic reticulum stress precedes oxidative stress in antibiotic-induced cholestasis and cytotoxicity in human hepatocytes. Free Radical Biology and Medicine, 115, 166–178. 10.1016/j.freeradbiomed.2017.11.017

Callegaro, G., Kunnen, S. J., Trairatphisan, P., Grosdidier, S., Niemeijer, M., den Hollander, W., Guney, E., Piñero Gonzalez, J., Furlong, L., Webster, Y. W., Saez-Rodriguez, J., Sutherland, J. J., Mollon, J., Stevens, J. L., & van de Water, B. (2021). The human hepatocyte TXG-MAPr: gene co-expression network modules to support mechanism-based risk assessment. Archives of Toxicology, 95(12), 3745–3775. 10.1007/s00204-021-03141-w

Cao, S. S., & Kaufman, R. J. (2014). Endoplasmic reticulum stress and oxidative stress in cell fate decision and human disease. In Antioxidants and Redox Signaling (Vol. 21, Issue 3). 10.1089/ars.2014.5851

Chatterjee, S., & Annaert, P. (2018). Drug-induced Cholestasis: Mechanisms, Models, and Markers. Current Drug Metabolism, 19(10). 10.2174/1389200219666180427165035

Chen, C., Krausz, K. W., Shah, Y. M., Idle, J. R., & Gonzalez, F. J. (2009). Serum metabolomics reveals irreversible inhibition of fatty acid β-oxidation through the suppression of PPARα activation as a contributing mechanism of acetaminophen-induced hepatotoxicity. Chemical Research in Toxicology, 22(4), 699–707. 10.1021/tx800464q

Choudhuri, S., Patton, G. W., Chanderbhan, R. F., Mattia, A., & Klaassen, C. D. (2018). From classical toxicology to Tox21: Some critical conceptual and technological advances in the molecular understanding of the toxic response beginning from the last Quarter of the 20th century. Toxicological Sciences, 161(1). 10.1093/toxsci/kfx186

Dasgupta, A., & Klein, K. (2014). Methods for Measuring Oxidative Stress in the Laboratory. In Antioxidants in Food, Vitamins and Supplements (pp. 19–40). Elsevier. 10.1016/B978-0-12-405872-9.00002-1

European Union. (2006). Regulation (EC) No 1907/2006 of the European Parliament and of the Council of 18 December 2006 concerning the Registration, Evaluation, Authorisation and Restriction of Chemicals (REACH), establishing a European Chemicals Agency, amending Directive 1999/45/EC and repealing Council Regulation (EEC) No 793/93 and Commission Regulation (EC) No 1488/94 as well as Council Directive 76/769/EEC and Commission Directives 91/155/EEC, 93/67/EEC, 93/105/EC and 2000/21/EC. OJ L, 396, 1–849.

European Union. (2009). Regulation (EC) No 1223/2009 of the European Parliament and of the Council of 30 November 2009 on cosmetic products. OJ L, 342, 59–209.

Fan, X., Lobenhofer, E. K., Chen, M., Shi, W., Huang, J., Luo, J., Zhang, J., Walker, S. J., Chu, T. M., Li, L., Wolfinger, R., Bao, W., Paules, R. S., Bushel, P. R., Li, J., Shi, T., Nikolskaya, T., Nikolsky, Y., Hong, H., … Tong, W. (2010). Consistency of predictive signature genes and classifiers generated using different microarray platforms. Pharmacogenomics Journal, 10(4), 247–257. 10.1038/tpj.2010.34

Farmahin, R., Williams, A., Kuo, B., Chepelev, N. L., Thomas, R. S., Barton-Maclaren, T. S., Curran, I. H., Nong, A., Wade, M. G., & Yauk, C. L. (2017). Recommended approaches in the application of toxicogenomics to derive points of departure for chemical risk assessment. Archives of Toxicology, 91(5), 2045–2065. 10.1007/s00204-016-1886-5

Farré, M., Roset, P. N., Abanades, S., Menoyo, E., Alvarez, Y., Rovira, M., & Baena, A. (2008). Study of paracetamol 1g oral solution bioavailability. Methods and Findings in Experimental and Clinical Pharmacology, 30(1), 37. 10.1358/mf.2008.30.1.1159648

Friedman, K. P., Gagne, M., Loo, L. H., Karamertzanis, P., Netzeva, T., Sobanski, T., Franzosa, J. A., Richard, A. M., Lougee, R. R., Gissi, A., Lee, J. Y. J., Angrish, M., Dorne, J. lou, Foster, S., Raffaele, K., Bahadori, T., Gwinn, M. R., Lambert, J., Whelan, M., … Thomas, R. S. (2020). Utility of in Vitro Bioactivity as a Lower Bound Estimate of in Vivo Adverse Effect Levels and in Risk-Based Prioritization. Toxicological Sciences, 173(1), 202–225. 10.1093/toxsci/kfz201

Gijbels, E., Vilas-Boas, V., Annaert, P., Vanhaecke, T., Devisscher, L., & Vinken, M. (2020). Robustness testing and optimization of an adverse outcome pathway on cholestatic liver injury. Archives of Toxicology, 94(4), 1151–1172. 10.1007/s00204-020-02691-9

Gijbels, E., Vilas-Boas, V., Deferm, N., Devisscher, L., Jaeschke, H., Annaert, P., & Vinken, M. (2019). Mechanisms and in vitro models of drug-induced cholestasis. In Archives of Toxicology (Vol. 93, Issue 5). 10.1007/s00204-019-02437-2

Grefhorst, A., van de Peppel, I. P., Larsen, L. E., Jonker, J. W., & Holleboom, A. G. (2021). The Role of Lipophagy in the Development and Treatment of Non-Alcoholic Fatty Liver Disease. Frontiers in Endocrinology, 11. 10.3389/fendo.2020.601627

Grünig, D., Szabo, L., Marbet, M., & Krähenbühl, S. (2020). Valproic acid affects fatty acid and triglyceride metabolism in HepaRG cells exposed to fatty acids by different mechanisms. Biochemical Pharmacology, 177. 10.1016/j.bcp.2020.113860

Gu, X., & Manautou, J. E. (2012). Molecular mechanisms underlying chemical liver injury. Expert Reviews in Molecular Medicine, 14, e4. 10.1017/S1462399411002110

Guillouzo, A., Corlu, A., Aninat, C., Glaise, D., Morel, F., & Guguen-Guillouzo, C. (2007). The human hepatoma HepaRG cells: A highly differentiated model for studies of liver metabolism and toxicity of xenobiotics. Chemico-Biological Interactions, 168(1), 66–73. 10.1016/j.cbi.2006.12.003

Guo, L., Lobenhofer, E. K., Wang, C., Shippy, R., Harris, S. C., Zhang, L., Mei, N., Chen, T., Herman, D., Goodsaid, F. M., Hurban, P., Phillips, K. L., Xu, J., Deng, X., Sun, Y. A., Tong, W., Dragan, Y. P., & Shi, L. (2006). Rat toxicogenomic study reveals analytical consistency across microarray platforms. Nature Biotechnology, 24(9), 1162–1169. 10.1038/nbt1238

Gustafson, E., Debruyne, C., De Troyer, O., Rogiers, V., Vinken, M., & Vanhaecke, T. (2020). Screening of repeated dose toxicity data in safety evaluation reports of cosmetic ingredients issued by the Scientific Committee on Consumer Safety between 2009 and 2019. Archives of Toxicology, 94(11), 3723–3735. 10.1007/s00204-020-02868-2

Hamon, J., Jennings, P., & Bois, F. Y. (2014). Systems biology modeling of omics data: Effect of cyclosporine a on the Nrf2 pathway in human renal cells. BMC Systems Biology, 8(1). 10.1186/1752-0509-8-76

Harrill, J. A., Everett, L. J., Haggard, D. E., Bundy, J. L., Willis, C. M., Shah, I., Friedman, K. P., Basili, D., Middleton, A., & Judson, R. S. (2024). Exploring the effects of experimental parameters and data modeling approaches on in vitro transcriptomic point-of-departure estimates. Toxicology, 501, 153694. 10.1016/j.tox.2023.153694

Harrill, J., Shah, I., Setzer, R. W., Haggard, D., Auerbach, S., Judson, R., & Thomas, R. S. (2019). Considerations for strategic use of high-throughput transcriptomics chemical screening data in regulatory decisions. Current Opinion in Toxicology, 15, 64–75. 10.1016/j.cotox.2019.05.004

Hayes, A. W., & Kruger, C. L. (2014). Hayes’ Principles and Methods of Toxicology. In Hayes’ Principles and Methods of Toxicology (Sixth Edition). CRC Press, Taylor & Francis Group. 10.1201/b17359

Hothorn, T., & Hornik, K. (2022). exactRankTests: Exact Distributions for Rank and Permutation Tests. https://CRAN.R-project.org/package=exactRankTests

Ipsen, D. H., Lykkesfeldt, J., & Tveden-Nyborg, P. (2018). Molecular mechanisms of hepatic lipid accumulation in non-alcoholic fatty liver disease. Cellular and Molecular Life Sciences, 75(18), 3313–3327. 10.1007/s00018-018-2860-6

Itabe, H., Yamaguchi, T., Nimura, S., & Sasabe, N. (2017). Perilipins: a diversity of intracellular lipid droplet proteins. Lipids in Health and Disease, 16(1), 83. 10.1186/s12944-017-0473-y

Jaeschke, H., McGill, M. R., & Ramachandran, A. (2012). Oxidant stress, mitochondria, and cell death mechanisms in drug-induced liver injury: Lessons learned from acetaminophen hepatotoxicity. In Drug Metabolism Reviews (Vol. 44, Issue 1). 10.3109/03602532.2011.602688

Jaeschke, H., & Ramachandran, A. (2018). Oxidant Stress and Lipid Peroxidation in Acetaminophen Hepatotoxicity. *Reactive Oxygen Species (Apex*, N.C.), 5(15), 145–158.

James, L. P., Capparelli, E. v., Simpson, P. M., Letzig, L., Roberts, D., Hinson, J. A., Kearns, G. L., Blumer, J. L., & Sullivan, J. E. (2008). Acetaminophen-associated hepatic injury: Evaluation of acetaminophen protein adducts in children and adolescents with acetaminophen overdose. Clinical Pharmacology and Therapeutics, 84(6). 10.1038/clpt.2008.190

Johnson, K. J., Auerbach, S. S., Stevens, T., Barton-Maclaren, T. S., Costa, E., Currie, R. A., Dalmas Wilk, D., Haq, S., Rager, J. E., Reardon, A. J. F., Wehmas, L., Williams, A., O’Brien, J., Yauk, C., LaRocca, J. L., & Pettit, S. (2022). A Transformative Vision for an Omics-Based Regulatory Chemical Testing Paradigm. Toxicological Sciences, 190(2), 127–132. 10.1093/toxsci/kfac097

Kadmon, M., Klünemann, C., Böhme, M., Ishikawa, T., Gorgas, K., Otto, G., Herfarth, C., & Keppler, D. (1993). Inhibition by cyclosporin A of Adenosine triphosphate-dependent transport from the hepatocyte into bile. Gastroenterology, 104(5), 1507–1514. 10.1016/0016-5085(93)90363-H

Kennedy, D., Samali, A., & Jäger, R. (2015). Methods for Studying ER Stress and UPR Markers in Human Cells (pp. 3–18). 10.1007/978-1-4939-2522-3_1

Kralj, T., Brouwer, K. L. R., & Creek, D. J. (2021). Analytical and Omics-Based Advances in the Study of Drug-Induced Liver Injury. Toxicological Sciences, 183(1), 1–13. 10.1093/toxsci/kfab069

Kuleshov, M. V., Jones, M. R., Rouillard, A. D., Fernandez, N. F., Duan, Q., Wang, Z., Koplev, S., Jenkins, S. L., Jagodnik, K. M., Lachmann, A., McDermott, M. G., Monteiro, C. D., Gundersen, G. W., & Ma’ayan, A. (2016). Enrichr: a comprehensive gene set enrichment analysis web server 2016 update. Nucleic Acids Research, 44(W1), W90–W97. 10.1093/nar/gkw377

Kültz, D. (2005). Molecular and evolutionary basis of the cellular stress response. Annual Review of Physiology, 67, 225–257. 10.1146/annurev.physiol.67.040403.103635

Lichtenstein, D., Luckert, C., Alarcan, J., de Sousa, G., Gioutlakis, M., Katsanou, E. S., Konstantinidou, P., Machera, K., Milani, E. S., Peijnenburg, A., Rahmani, R., Rijkers, D., Spyropoulou, A., Stamou, M., Stoopen, G., Sturla, S. J., Wollscheid, B., Zucchini-Pascal, N., Braeuning, A., & Lampen, A. (2020). An adverse outcome pathway-based approach to assess steatotic mixture effects of hepatotoxic pesticides in vitro. Food and Chemical Toxicology, 139, 111283. 10.1016/j.fct.2020.111283

Liu, X., Taylor, S. A., Celaj, S., Levitsky, J., & Green, R. M. (2022). Expression of unfolded protein response genes in post-transplantation liver biopsies. BMC Gastroenterology, 22(1), 380. 10.1186/s12876-022-02459-8

Liu, Z., Huang, R., Roberts, R., & Tong, W. (2019). Toxicogenomics: A 2020 Vision. Trends in Pharmacological Sciences, 40(2), 92–103. 10.1016/j.tips.2018.12.001

Love, M. I., Huber, W., & Anders, S. (2014). Moderated estimation of fold change and dispersion for RNA-seq data with DESeq2. Genome Biology, 15(12), 550. 10.1186/s13059-014-0550-8

Luckert, C., Braeuning, A., de Sousa, G., Durinck, S., Katsanou, E. S., Konstantinidou, P., Machera, K., Milani, E. S., Peijnenburg, A. A. C. M., Rahmani, R., Rajkovic, A., Rijkers, D., Spyropoulou, A., Stamou, M., Stoopen, G., Sturla, S., Wollscheid, B., Zucchini-Pascal, N., & Lampen, A. (2018). Adverse Outcome Pathway-Driven Analysis of Liver Steatosis *in Vitro* : A Case Study with Cyproconazole. Chemical Research in Toxicology, 31(8), 784–798. 10.1021/acs.chemrestox.8b00112

Malhotra, J. D., & Kaufman, R. J. (2007). Endoplasmic Reticulum Stress and Oxidative Stress: A Vicious Cycle or a Double-Edged Sword? Antioxidants & Redox Signaling, 9(12), 2277–2294. 10.1089/ars.2007.1782

McGill, M. R., & Jaeschke, H. (2013). Metabolism and disposition of acetaminophen: Recent advances in relation to hepatotoxicity and diagnosis. In Pharmaceutical Research (Vol. 30, Issue 9). 10.1007/s11095-013-1007-6

Mellor, C. L., Steinmetz, F. P., & Cronin, M. T. D. (2016). The identification of nuclear receptors associated with hepatic steatosis to develop and extend adverse outcome pathways. Critical Reviews in Toxicology, 46(2), 138–152. 10.3109/10408444.2015.1089471

More, S. J., Bampidis, V., Benford, D., Bragard, C., Halldorsson, T. I., Hernández-Jerez, A. F., Bennekou, S. H., Koutsoumanis, K., Lambré, C., Machera, K., Mennes, W., Mullins, E., Nielsen, S. S., Schrenk, D., Turck, D., Younes, M., Aerts, M., Edler, L., Sand, S., … Schlatter, J. (2022). Guidance on the use of the benchmark dose approach in risk assessment. EFSA Journal, 20(10). 10.2903/j.efsa.2022.7584

National Toxicology Program. (2018). NTP Research Report on National Toxicology Program Approach to Genomic Dose-Response Modeling. 10.22427/NTP-RR-5

Nguyen, P., Leray, V., Diez, M., Serisier, S., le Bloc’H, J., Siliart, B., & Dumon, H. (2008). Liver lipid metabolism. Journal of Animal Physiology and Animal Nutrition, 92(3). 10.1111/j.1439-0396.2007.00752.x

Pacheco-Alvarez, D., Solórzano-Vargas, R. S., & del Río, A. L. (2002). Biotin in metabolism and its relationship to human disease. In Archives of Medical Research (Vol. 33, Issue 5). 10.1016/S0188-4409(02)00399-5

Panico, A., Serio, F., Bagordo, F., Grassi, T., Idolo, A., de Giorgi, M., Guido, M., Congedo, M., & de Donno, A. (2019). Skin safety and health prevention: An overview of chemicals in cosmetic products. Journal of Preventive Medicine and Hygiene, 60(1), E50–E57. 10.15167/2421-4248/jpmh2019.60.1.1080

Pavlik, L., Regev, A., Ardayfio, P. A., & Chalasani, N. P. (2019). Drug-Induced Steatosis and Steatohepatitis: The Search for Novel Serum Biomarkers Among Potential Biomarkers for Non-Alcoholic Fatty Liver Disease and Non-Alcoholic Steatohepatitis. Drug Safety, 42(6), 701–711. 10.1007/s40264-018-00790-2

Phillips, J. R., Svoboda, D. L., Tandon, A., Patel, S., Sedykh, A., Mav, D., Kuo, B., Yauk, C. L., Yang, L., Thomas, R. S., Gift, J. S., Davis, J. A., Olszyk, L., Merrick, B. A., Paules, R. S., Parham, F., Saddler, T., Shah, R. R., & Auerbach, S. S. (2019). BMDExpress 2: enhanced transcriptomic dose-response analysis workflow. Bioinformatics, 35(10), 1780–1782. 10.1093/bioinformatics/bty878

Podtelezhnikov, A. A., Monroe, J. J., Aslamkhan, A. G., Pearson, K., Qin, C., Tamburino, A. M., Loboda, A. P., Glaab, W. E., Sistare, F. D., & Tanis, K. Q. (2020). Quantitative Transcriptional Biomarkers of Xenobiotic Receptor Activation in Rat Liver for the Early Assessment of Drug Safety Liabilities. Toxicological Sciences, 175(1), 98–112. 10.1093/toxsci/kfaa026

Price, R. J., Scott, M. P., Giddings, A. M., Walters, D. G., Stierum, R. H., Meredith, C., & Lake, B. G. (2008). Effect of butylated hydroxytoluene, curcumin, propyl gallate and thiabendazole on cytochrome P450 forms in cultured human hepatocytes. Xenobiotica, 38(6), 574–586. 10.1080/00498250802008615

R Core Team. (2022). R: A Language and Environment for Statistical Computing. https://www.R-project.org/

Ramachandran, A., & Jaeschke, H. (2018). Acetaminophen toxicity: Novel insights into mechanisms and future perspectives. In Gene Expression (Vol. 18, Issue 1). 10.3727/105221617X15084371374138

Ramaiahgari, S. C., Auerbach, S. S., Saddler, T. O., Rice, J. R., Dunlap, P. E., Sipes, N. S., DeVito, M. J., Shah, R. R., Bushel, P. R., Merrick, B. A., Paules, R. S., & Ferguson, S. S. (2019). The Power of Resolution: Contextualized Understanding of Biological Responses to Liver Injury Chemicals Using High-throughput Transcriptomics and Benchmark Concentration Modeling. Toxicological Sciences, 169(2), 553–566. 10.1093/toxsci/kfz065

Ramaiahgari, S. C., Waidyanatha, S., Dixon, D., DeVito, M. J., Paules, R. S., & Ferguson, S. S. (2017). From the Cover: Three-Dimensional (3D) HepaRG Spheroid Model With Physiologically Relevant Xenobiotic Metabolism Competence and Hepatocyte Functionality for Liver Toxicity Screening. Toxicological Sciences, 159(1), 124–136. 10.1093/toxsci/kfx122

Rao, S. R., Ajitkumar, S., Subbarayan, R., & Girija, D. M. (2018). Cyclosporine-A induces endoplasmic reticulum stress in human gingival fibroblasts – An in vitro study. Journal of Oral Biology and Craniofacial Research, 8(3), 165–167. 10.1016/j.jobcr.2016.11.002

Reardon, A. J. F., Farmahin, R., Williams, A., Meier, M. J., Addicks, G. C., Yauk, C. L., Matteo, G., Atlas, E., Harrill, J., Everett, L. J., Shah, I., Judson, R., Ramaiahgari, S., Ferguson, S. S., & Barton-Maclaren, T. S. (2023). From vision toward best practices: Evaluating in vitro transcriptomic points of departure for application in risk assessment using a uniform workflow. Frontiers in Toxicology, 5. 10.3389/ftox.2023.1194895

Rogiers, V., Benfenati, E., Bernauer, U., Bodin, L., Carmichael, P., Chaudhry, Q., Coenraads, P. J., Cronin, M. T. D., Dent, M., Dusinska, M., Ellison, C., Ezendam, J., Gaffet, E., Galli, C. L., Goebel, C., Granum, B., Hollnagel, H. M., Kern, P. S., Kosemund-Meynen, K., … Worth, A. (2020). The way forward for assessing the human health safety of cosmetics in the EU - Workshop proceedings. Toxicology, 436, 152421. 10.1016/j.tox.2020.152421

Román, I. D., & Coleman, R. (1994). Disruption of canalicular function in isolated rat hepatocyte couplets caused by cyclosporin A. Biochemical Pharmacology, 48(12), 2181–2188. 10.1016/0006-2952(94)90352-2

Russmann, S., Kullak-Ublick, G., & Grattagliano, I. (2009). Current Concepts of Mechanisms in Drug-Induced Hepatotoxicity. Current Medicinal Chemistry, 16(23), 3041–3053. 10.2174/092986709788803097

Saarimäki, L. A., Morikka, J., Pavel, A., Korpilähde, S., Del Giudice, G., Federico, A., Fratello, M., Serra, A., & Greco, D. (2023). Toxicogenomics Data for Chemical Safety Assessment and Development of New Approach Methodologies: An Adverse Outcome Pathway-Based Approach. *Advanced Science (Weinheim, Baden-Wurttemberg*, Germany), 10(2), e2203984. 10.1002/advs.202203984

SCCS. (2009). Scientific Committee on Consumer Safety, opinion on Triclosan, addemdum to the SCCP opinion on Triclosan (SCCP/1192/08). 10.2772/96027

SCCS. (2010). Scientific Committee on Consumer Safety, opinion on 2,7-Naphthalenediol, COLIPA n° A19. 10.2772/27368

SCCS. (2021). Scientific Committee on Consumer Safety, opinion on Butylated Hydroxytoluene (BHT). 10.2875/53206

Schumacher, J., & Guo, G. (2015). Mechanistic Review of Drug-Induced Steatohepatitis. Toxicology and Applied Pharmacology, 289(1), 40–47. 10.1016/J.TAAP.2015.08.022

Scientific Committee on Consumer Safety. (2021). The SCCS Notes of Guidance for the Testing of Cosmetic Ingredients and their Safety Evaluation, 11th revision. 10.2875/273162

Selvaraj, S., Oh, J.-H., & Borlak, J. (2020). An adverse outcome pathway for immune-mediated and allergic hepatitis: a case study with the NSAID diclofenac. Archives of Toxicology, 94(8), 2733– 2748. 10.1007/s00204-020-02767-6

Sevilla-Tirado, F. J., Gonzalez-Vallejo, E. B., Leary, A. C., Breedt, H. J., Hyde, V. J., & Fernandez-Hernando, N. (2003). Bioavailability of two new formulations of paracetamol, compared with three marketed formulations, in healthy volunteers. Methods and Findings in Experimental and Clinical Pharmacology, 25(7), 531. 10.1358/mf.2003.25.7.778092

Sharanek, A., Azzi, P. B.-E., Al-Attrache, H., Savary, C. C., Humbert, L., Rainteau, D., Guguen-Guillouzo, C., & Guillouzo, A. (2014). Different Dose-Dependent Mechanisms Are Involved in Early Cyclosporine A-Induced Cholestatic Effects in HepaRG Cells. Toxicological Sciences, 141(1), 244–253. 10.1093/toxsci/kfu122

Sharanek, A., Burban, A., Humbert, L., Bachour-El Azzi, P., Felix-Gomes, N., Rainteau, D., & Guillouzo, A. (2015). Cellular Accumulation and Toxic Effects of Bile Acids in Cyclosporine A-Treated HepaRG Hepatocytes. Toxicological Sciences, 147(2), 573–587. 10.1093/toxsci/kfv155

Song, Y., Zhang, C., Lei, H., Qin, M., Chen, G., Wu, F., Chen, C., Cao, Z., Zhang, C., Wu, M., Chen, X., & Zhang, L. (2022). Characterization of triclosan-induced hepatotoxicity and triclocarban-triggered enterotoxicity in mice by multiple omics screening. Science of The Total Environment, 838, 156570. 10.1016/J.SCITOTENV.2022.156570

Spyker, D. A., Dart, R. C., Yip, L., Reynolds, K., Brittain, S., & Yarema, M. (2022). Population pharmacokinetic analysis of acetaminophen overdose with immediate release, extended release and modified release formulations. Clinical Toxicology, 60(10), 1113–1121. 10.1080/15563650.2022.2114361

Sutherland, J. J., Jolly, R. A., Goldstein, K. M., & Stevens, J. L. (2016). Assessing Concordance of Drug-Induced Transcriptional Response in Rodent Liver and Cultured Hepatocytes. PLoS Computational Biology, 12(3), e1004847. 10.1371/journal.pcbi.1004847

Sutherland, J. J., Webster, Y. W., Willy, J. A., Searfoss, G. H., Goldstein, K. M., Irizarry, A. R., Hall, D. G., & Stevens, J. L. (2018). Toxicogenomic module associations with pathogenesis: a network-based approach to understanding drug toxicity. The Pharmacogenomics Journal, 18(3), 377–390. 10.1038/tpj.2017.17

Sztalryd, C., & Kimmel, A. R. (2014). Perilipins: Lipid droplet coat proteins adapted for tissue-specific energy storage and utilization, and lipid cytoprotection. Biochimie, 96, 96–101. 10.1016/j.biochi.2013.08.026

Tan, C., & Graudins, A. (2006). Comparative pharmacokinetics of Panadol Extend and immediate-release paracetamol in a simulated overdose model. Emergency Medicine Australasia, 18(4), 398–403. 10.1111/j.1742-6723.2006.00873.x

Tanabe, S., O’Brien, J., Tollefsen, K. E., Kim, Y., Chauhan, V., Yauk, C., Huliganga, E., Rudel, R. A., Kay, J. E., Helm, J. S., Beaton, D., Filipovska, J., Sovadinova, I., Garcia-Reyero, N., Mally, A., Poulsen, S. S., Delrue, N., Fritsche, E., Luettich, K., … FitzGerald, R. (2022). Reactive Oxygen Species in the Adverse Outcome Pathway Framework: Toward Creation of Harmonized Consensus Key Events. Frontiers in Toxicology, 4. 10.3389/ftox.2022.887135

Tazuma, S. (2006). Cyclosporin A and cholestasis: Its mechanism(s) and clinical relevancy. Hepatology Research, 34(3), 135–136. 10.1016/j.hepres.2005.12.009

Thomas, R. S., Bahadori, T., Buckley, T. J., Cowden, J., Deisenroth, C., Dionisio, K. L., Frithsen, J. B., Grulke, C. M., Gwinn, M. R., Harrill, J. A., Higuchi, M., Houck, K. A., Hughes, M. F., Sidney Hunter, E., Isaacs, K. K., Judson, R. S., Knudsen, T. B., Lambert, J. C., Linnenbrink, M., … Williams, A. J. (2019). The next generation blueprint of computational toxicology at the U.S. Environmental protection agency. Toxicological Sciences, 169(2), 317–332. 10.1093/toxsci/kfz058

Thomas, R. S., Philbert, M. A., Auerbach, S. S., Wetmore, B. A., Devito, M. J., Cote, I., Rowlands, J. C., Whelan, M. P., Hays, S. M., Andersen, M. E., Meek, M. E. B., Reiter, L. W., Lambert, J. C., Clewell, H. J., Stephens, M. L., Zhao, Q. J., Wesselkamper, S. C., Flowers, L., Carney, E. W., … Nong, A. (2013). Incorporating new technologies into toxicity testing and risk assessment: Moving from 21st century vision to a data-driven framework. Toxicological Sciences, 136(1), 4–18. 10.1093/toxsci/kft178

Thomas, R. S., Wesselkamper, S. C., Wang, N. C. Y., Zhao, Q. J., Petersen, D. D., Lambert, J. C., Cote, I., Yang, L., Healy, E., Black, M. B., Clewell, H. J., Allen, B. C., & Andersen, M. E. (2013). Temporal Concordance Between Apical and Transcriptional Points of Departure for Chemical Risk Assessment. Toxicological Sciences, 134(1), 180–194. 10.1093/toxsci/kft094

Vahle, J. L., Anderson, U., Blomme, E. A. G., Hoflack, J.-C., & Stiehl, D. P. (2018). Use of toxicogenomics in drug safety evaluation: Current status and an industry perspective. Regulatory Toxicology and Pharmacology, 96, 18–29. 10.1016/j.yrtph.2018.04.011

van Breda, S. G. J., Claessen, S. M. H., van Herwijnen, M., Theunissen, D. H. J., Jennen, D. G. J., de Kok, T. M. C. M., & Kleinjans, J. C. S. (2018). Integrative omics data analyses of repeated dose toxicity of valproic acid in vitro reveal new mechanisms of steatosis induction. Toxicology, 393, 160–170. 10.1016/j.tox.2017.11.013

Van den Hof, W. F. P. M., Ruiz-Aracama, A., Van Summeren, A., Jennen, D. G. J., Gaj, S., Coonen, M. L. J., Brauers, K., Wodzig, W. K. W. H., van Delft, J. H. M., & Kleinjans, J. C. S. (2015). Integrating multiple omics to unravel mechanisms of Cyclosporin A induced hepatotoxicity in vitro. Toxicology in Vitro, 29(3), 489–501. 10.1016/j.tiv.2014.12.016

Verheijen, M. CT., Meier, M. J., Asensio, J. O., Gant, T. W., Tong, W., Yauk, C. L., & Caiment, F. (2022). R-ODAF: Omics data analysis framework for regulatory application. Regulatory Toxicology and Pharmacology, 131, 105143. 10.1016/j.yrtph.2022.105143

Vinken, M. (2019). Omics-based input and output in the development and use of adverse outcome pathways. Current Opinion in Toxicology, 18, 8–12. 10.1016/J.COTOX.2019.02.006

Vinken, M., Landesmann, B., Goumenou, M., Vinken, S., Shah, I., Jaeschke, H., Willett, C., Whelan, M., & Rogiers, V. (2013). Development of an Adverse Outcome Pathway From Drug-Mediated Bile Salt Export Pump Inhibition to Cholestatic Liver Injury. Toxicological Sciences, 136(1), 97–106. 10.1093/toxsci/kft177

Vinken, M., Maes, M., Vanhaecke, T., & Rogiers, V. (2013). Drug-Induced Liver Injury: Mechanisms, Types and Biomarkers. Current Medicinal Chemistry, 20(24), 3011–3021. 10.2174/0929867311320240006

Vinken, M., Pauwels, M., Ates, G., Vivier, M., Vanhaecke, T., & Rogiers, V. (2012). Screening of repeated dose toxicity data present in SCC(NF)P/SCCS safety evaluations of cosmetic ingredients. Archives of Toxicology, 86(3), 405–412. 10.1007/s00204-011-0769-z

Vitale, G., Mattiaccio, A., Conti, A., Berardi, S., Vero, V., Turco, L., Seri, M., & Morelli, M. C. (2023). Molecular and Clinical Links between Drug-Induced Cholestasis and Familial Intrahepatic Cholestasis. International Journal of Molecular Sciences, 24(6), 5823. 10.3390/ijms24065823

Vrijenhoek, N., Wehr, M. M., Kunnen, S. J., Wijaya, L. S., Callegaro, G., Moné, M., Escher, S. E., & van de Water, B. (2022). Application of high-throughput transcriptomics for mechanism-based biological read-across of short-chain carboxylic acid analogues of valproic acid. ALTEX. 10.14573/altex.2107261

Wang, C., Gong, B., Bushel, P. R., Thierry-Mieg, J., Thierry-Mieg, D., Xu, J., Fang, H., Hong, H., Shen, J., Su, Z., Meehan, J., Li, X., Yang, L., Li, H., Łabaj, P. P., Kreil, D. P., Megherbi, D., Gaj, S., Caiment, F., … Tong, W. (2014). The concordance between RNA-seq and microarray data depends on chemical treatment and transcript abundance. Nature Biotechnology, 32(9), 926–932. 10.1038/nbt.3001

Xie, Y., McGill, M. R., Cook, S. F., Sharpe, M. R., Winefield, R. D., Wilkins, D. G., Rollins, D. E., & Jaeschke, H. (2015). Time course of acetaminophen-protein adducts and acetaminophen metabolites in circulation of overdose patients and in HepaRG cells. Xenobiotica, 45(10), 1574–1583. 10.3109/00498254.2015.1026426

Xu, S., Chen, Y., Ma, Y., Liu, T., Zhao, M., Wang, Z., & Zhao, L. (2019). Lipidomic Profiling Reveals Disruption of Lipid Metabolism in Valproic Acid-Induced Hepatotoxicity. Frontiers in Pharmacology, 10. 10.3389/fphar.2019.00819

Yasumiba, S., Tazuma, S., Ochi, H., Chayama, K., & Kajiyama, G. (2001). Cyclosporin A reduces canalicular membrane fluidity and regulates transporter function in rats. Biochemical Journal, 354(3), 591–596. 10.1042/0264-6021:3540591

Yeakley, J. M., Shepard, P. J., Goyena, D. E., VanSteenhouse, H. C., McComb, J. D., & Seligmann, B. E. (2017). A trichostatin A expression signature identified by TempO-Seq targeted whole transcriptome profiling. PLOS ONE, 12(5), e0178302. 10.1371/journal.pone.0178302

Yin, W., Mendoza, L., Monzon-Sandoval, J., Urrutia, A. O., & Gutierrez, H. (2021). Emergence of co-expression in gene regulatory networks. PLoS ONE, 16(4), e0247671. 10.1371/journal.pone.0247671

Yoon, E., Babar, A., Choudhary, M., Kutner, M., & Pyrsopoulos, N. (2016). Acetaminophen-Induced Hepatotoxicity: a Comprehensive Update. Journal of Clinical and Translational Hepatology, 4(2), 131–142. 10.14218/JCTH.2015.00052

Zoupa, M., Zwart, E. P., Gremmer, E. R., Nugraha, A., Compeer, S., Slob, W., & van der Ven, L. T. M. (2020). Dose addition in chemical mixtures inducing craniofacial malformations in zebrafish (Danio rerio) embryos. Food and Chemical Toxicology, 137, 111117. 10.1016/j.fct.2020.111117

