## Supplementary Figures for "Qualitative and Quantitative Concentration-Response Modelling of Gene Co-expression Networks to Unlock Hepatotoxic Mechanisms for Next Generation Chemical Safety Assessment"

**Figure S1**

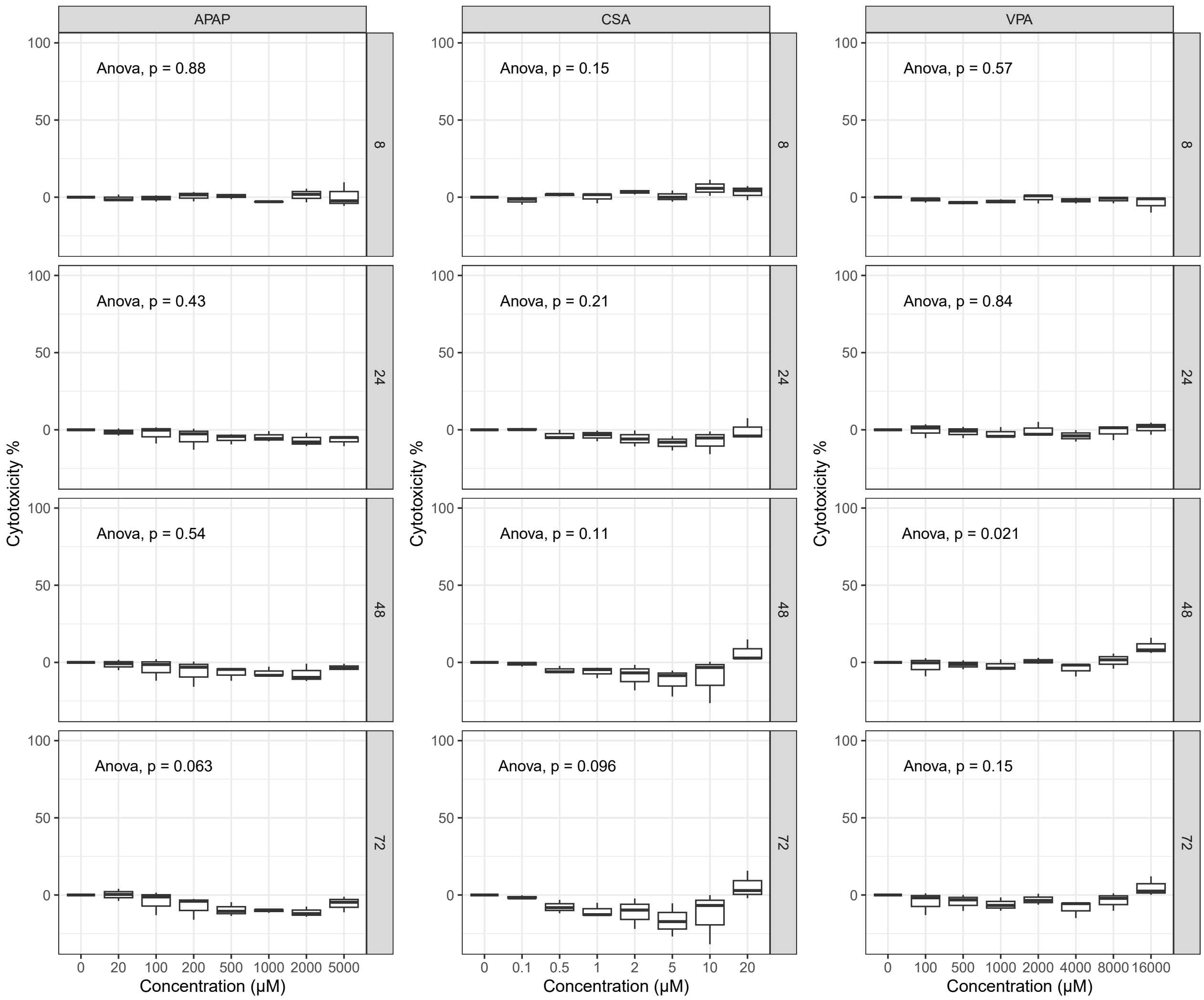

APAP
CSA
VPA
Anova, p = 0.88
Anova, p = 0.15
Anova, p = 0.57
Anova, p = 0.43
Anova, p = 0.21
Anova, p = 0.84
Anova, p = 0.54
Anova, p = 0.11
Anova, p = 0.021
Anova, p = 0.063
Anova, p = 0.096
Anova, p = 0.15
8
24
48
72
Cytotoxicity %
100
50
0
0 20 100 200 500 1000 2000 5000
0 0.1 0.5 1 2 5 10 20
0 100 500 1000 2000 4000 8000 16000
Concentration (µM)

#### Figure S2

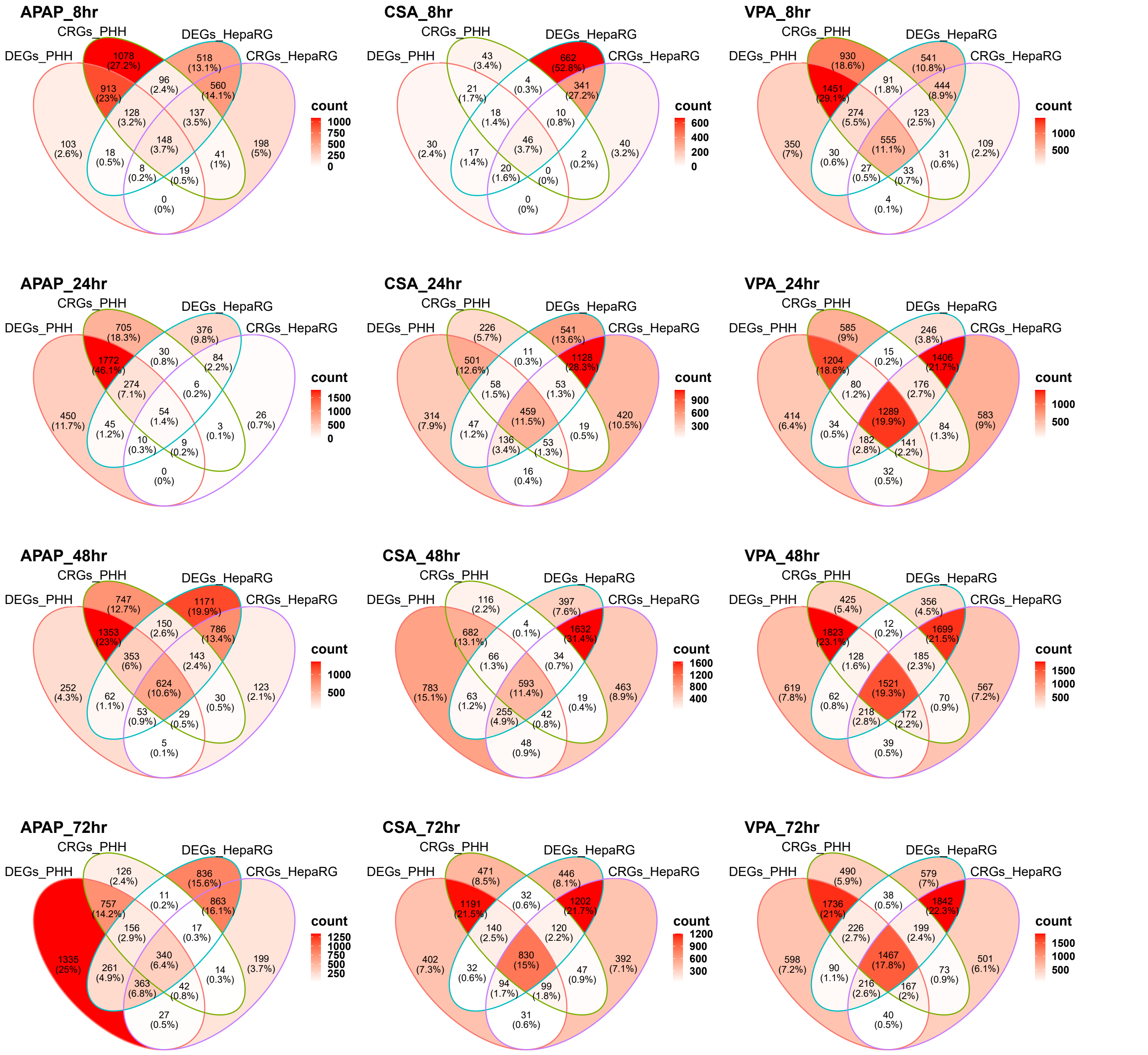
APAP_8hr
CSA_8hr
VPA_8hr
APAP_24hr
CSA_24hr
VPA_24hr
APAP_48hr
CSA_48hr
VPA_48hr
APAP_72hr
CSA_72hr
VPA_72hr
DEGs_PHH
CRGs_PHH
DEGs_HepaRG
CRGs_HepaRG
count

Figure S3

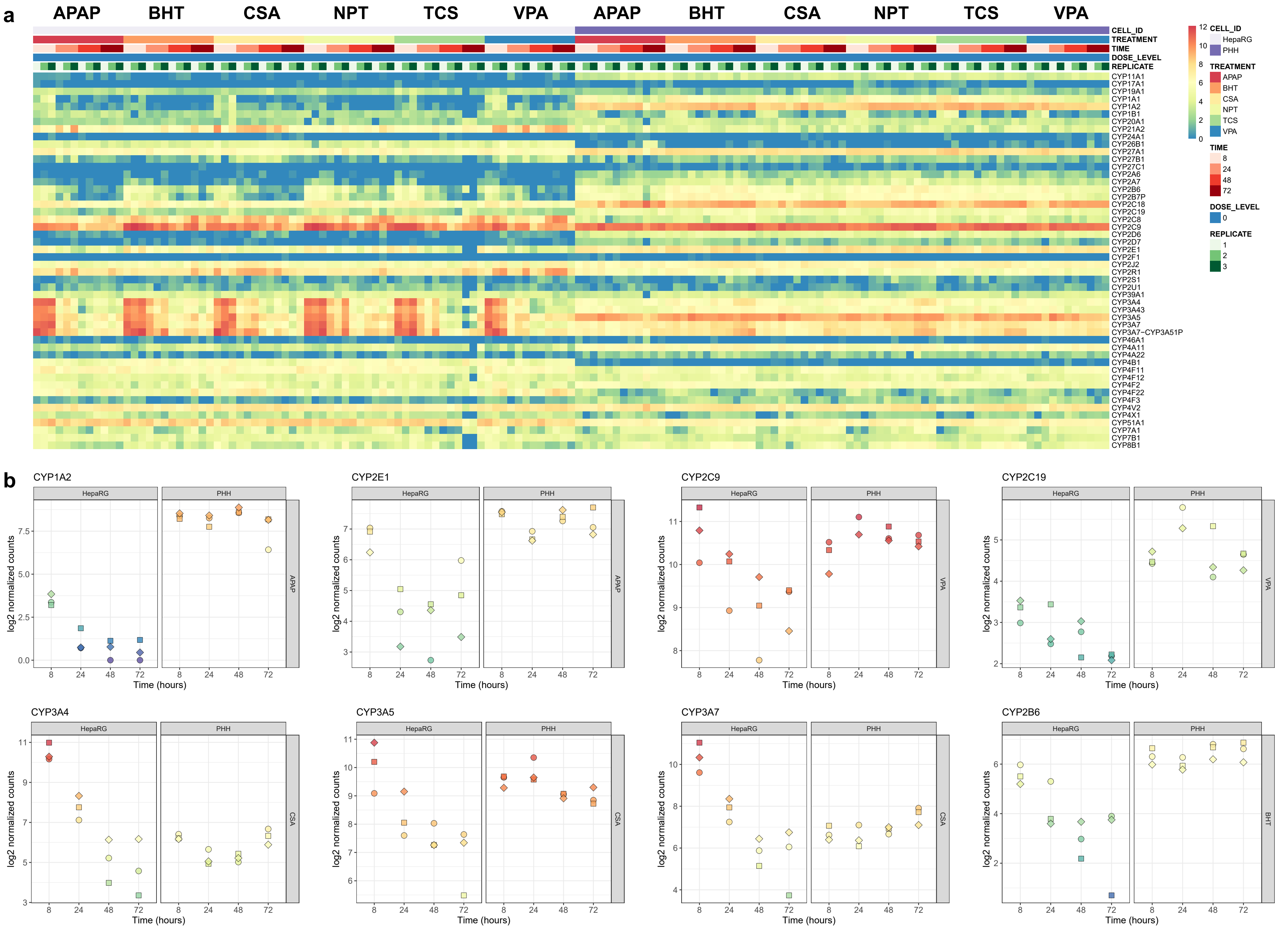
a
APAP
BHT
CSA
NPT
TCS
VPA
APAP
BHT
CSA
NPT
TCS
VPA
CELL_ID
TREATMENT
TIME
DOSE_LEVEL
REPLICATE
CYP11A1
CYP17A1
CYP19A1
CYP1A1
CYP1A2
CYP1B1
CYP20A1
CYP21A2
CYP24A1
CYP26B1
CYP27A1
CYP27B1
CYP27C1
CYP2A6
CYP2A7
CYP2B6
CYP2B7P
CYP2C18
CYP2C19
CYP2C8
CYP2C9
CYP2D6
CYP2D7
CYP2E1
CYP2F1
CYP2J2
CYP2R1
CYP2S1
CYP2U1
CYP39A1
CYP3A4
CYP3A43
CYP3A5
CYP3A7
CYP3A7−CYP3A51P
CYP46A1
CYP4A11
CYP4A22
CYP4B1
CYP4F11
CYP4F12
CYP4F2
CYP4F22
CYP4F3
CYP4V2
CYP4X1
CYP51A1
CYP7A1
CYP7B1
CYP8B1
12
10
8
6
4
2
0
CELL_ID
HepaRG
PHH
TREATMENT
APAP
BHT
CSA
NPT
TCS
VPA
TIME
8
24
48
72
DOSE_LEVEL
0
REPLICATE
1
2
3
b
CYP1A2
CYP2E1
CYP2C9
CYP2C19
CYP3A4
CYP3A5
CYP3A7
CYP2B6
HepaRG
PHH
APAP
VPA
CSA
BHT
log2 normalized counts
Time (hours)
8
24
48
72

#### Figure S4

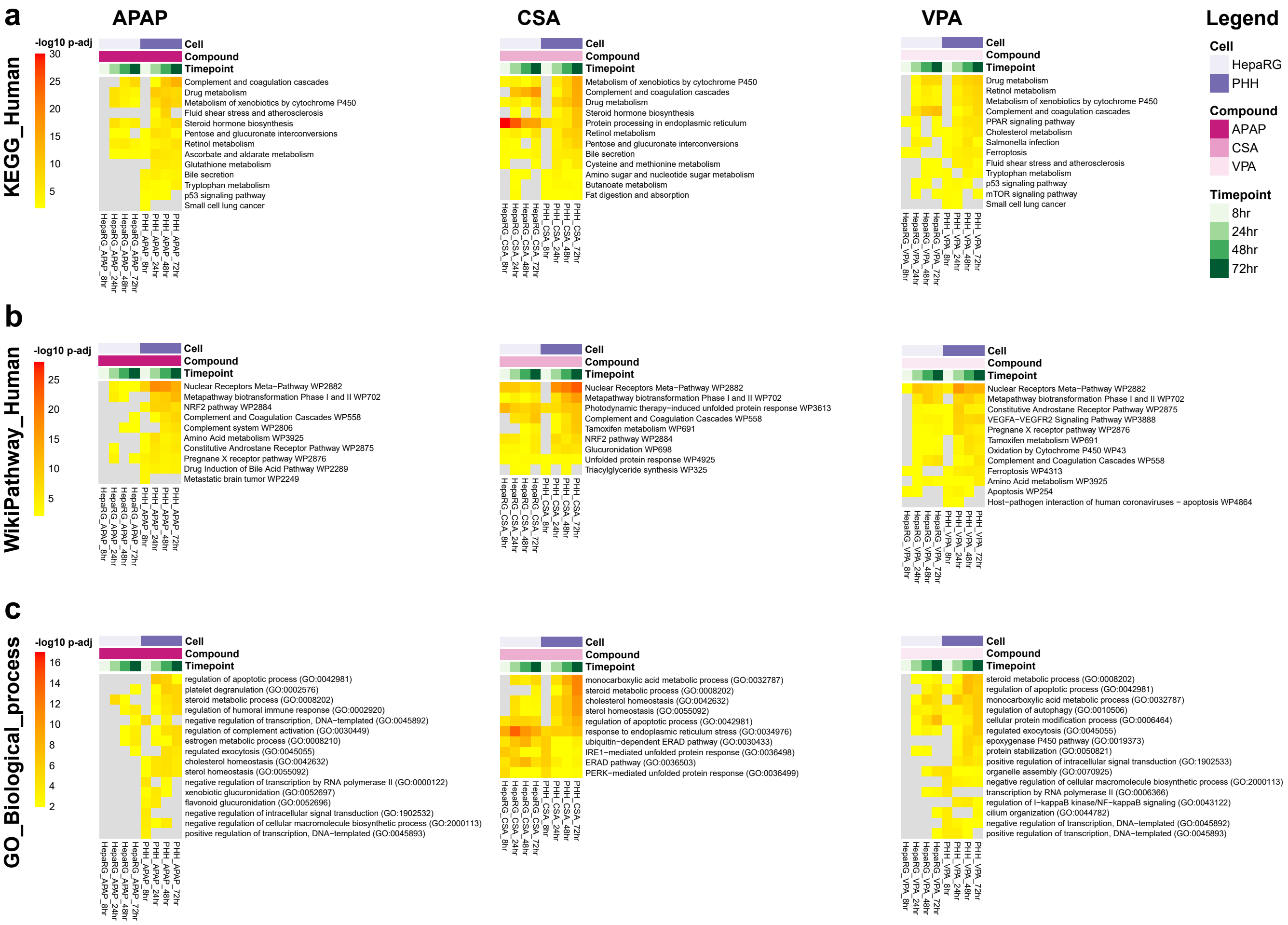

a
APAP
CSA
VPA
Legend
KEGG_Human
-log10 p-adj
Cell
Compound
Timepoint
Complement and coagulation cascades
Drug metabolism
Metabolism of xenobiotics by cytochrome P450
Fluid shear stress and atherosclerosis
Steroid hormone biosynthesis
Pentose and glucuronate interconversions
Retinol metabolism
Ascorbate and aldarate metabolism
Glutathione metabolism
Bile secretion
Tryptophan metabolism
p53 signaling pathway
Small cell lung cancer
Protein processing in endoplasmic reticulum
Cysteine and methionine metabolism
Amino sugar and nucleotide sugar metabolism
Butanoate metabolism
Fat digestion and absorption
PPAR signaling pathway
Cholesterol metabolism
Salmonella infection
Ferroptosis
mTOR signaling pathway
HepaRG
PHH
APAP
CSA
VPA
8hr
24hr
48hr
72hr
b
WikiPathway_Human
Nuclear Receptors Meta−Pathway WP2882
Metapathway biotransformation Phase I and II WP702
NRF2 pathway WP2884
Complement and Coagulation Cascades WP558
Complement system WP2806
Amino Acid metabolism WP3925
Constitutive Androstane Receptor Pathway WP2875
Pregnane X receptor pathway WP2876
Drug Induction of Bile Acid Pathway WP2289
Metastatic brain tumor WP2249
Photodynamic therapy−induced unfolded protein response WP3613
Tamoxifen metabolism WP691
Glucuronidation WP698
Unfolded protein response WP4925
Triacylglyceride synthesis WP325
VEGFA−VEGFR2 Signaling Pathway WP3888
Oxidation by Cytochrome P450 WP43
Ferroptosis WP4313
Apoptosis WP254
Host−pathogen interaction of human coronaviruses − apoptosis WP4864
c
GO_Biological_process
regulation of apoptotic process (GO:0042981)
platelet degranulation (GO:0002576)
steroid metabolic process (GO:0008202)
regulation of humoral immune response (GO:0002920)
negative regulation of transcription, DNA−templated (GO:0045892)
regulation of complement activation (GO:0030449)
estrogen metabolic process (GO:0008210)
regulated exocytosis (GO:0045055)
cholesterol homeostasis (GO:0042632)
sterol homeostasis (GO:0055092)
negative regulation of transcription by RNA polymerase II (GO:0000122)
xenobiotic glucuronidation (GO:0052697)
flavonoid glucuronidation (GO:0052696)
negative regulation of intracellular signal transduction (GO:1902532)
negative regulation of cellular macromolecule biosynthetic process (GO:2000113)
positive regulation of transcription, DNA−templated (GO:0045893)
monocarboxylic acid metabolic process (GO:0032787)
response to endoplasmic reticulum stress (GO:0034976)
ubiquitin−dependent ERAD pathway (GO:0030433)
IRE1−mediated unfolded protein response (GO:0036498)
ERAD pathway (GO:0036503)
PERK−mediated unfolded protein response (GO:0036499)
regulation of autophagy (GO:0010506)
cellular protein modification process (GO:0006464)
epoxygenase P450 pathway (GO:0019373)
protein stabilization (GO:0050821)
positive regulation of intracellular signal transduction (GO:1902533)
organelle assembly (GO:0070925)
transcription by RNA polymerase II (GO:0006366)
regulation of I−kappaB kinase/NF−kappaB signaling (GO:0043122)
cilium organization (GO:0044782)

#### Figure S5a

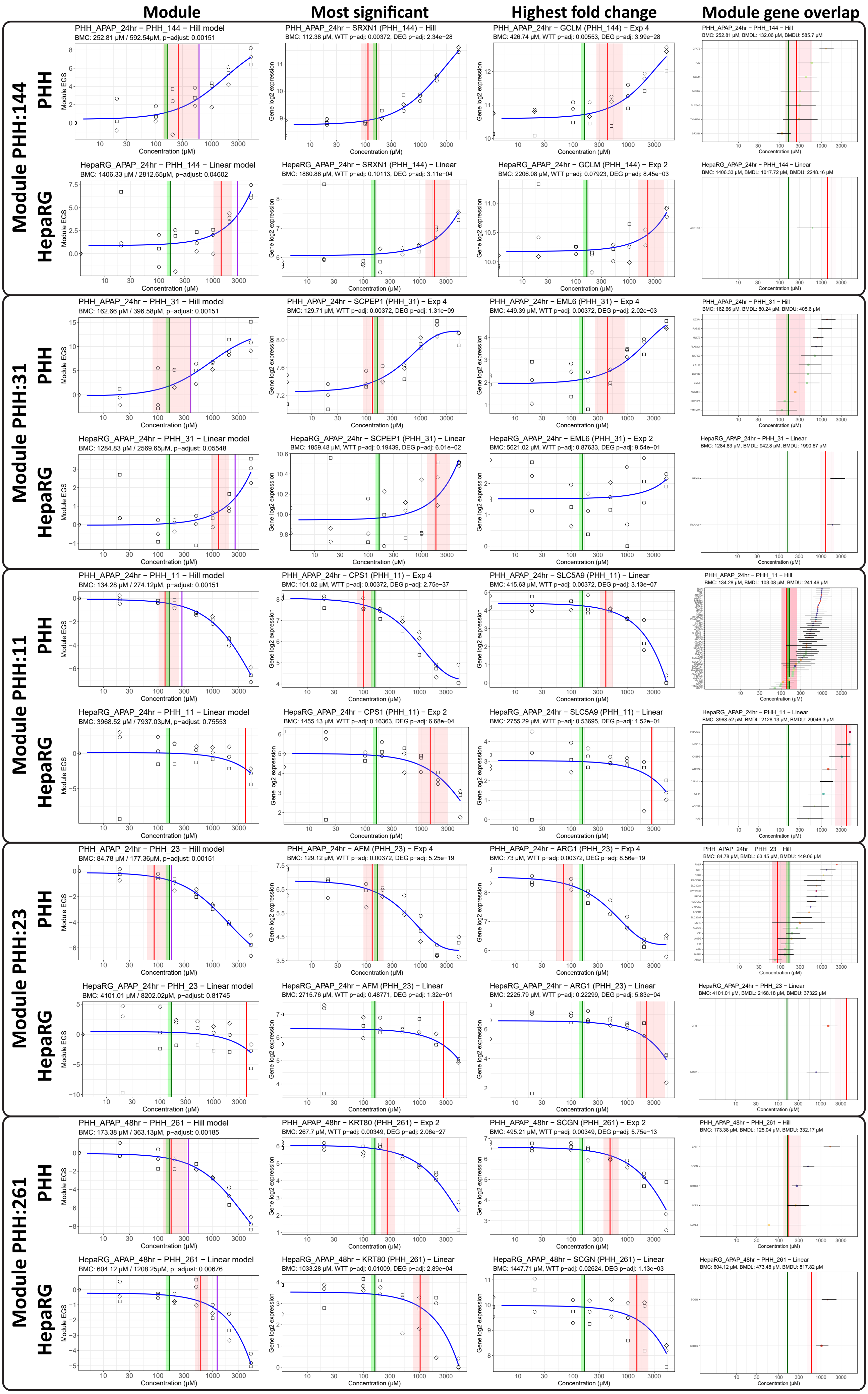

Module
Most significant
Highest fold change
Module gene overlap
Module PHH:144
PHH
HepaRG
PHH_APAP_24hr − PHH_144 − Hill model
BMC: 252.81 μM / 592.54μM, p-adjust: 0.00151
PHH_APAP_24hr − SRXN1 (PHH_144) − Hill
BMC: 112.38 μM, WTT p-adj: 0.00372, DEG p-adj: 2.34e−28
PHH_APAP_24hr − GCLM (PHH_144) − Exp 4
BMC: 426.74 μM, WTT p-adj: 0.00553, DEG p-adj: 3.99e−28
PHH_APAP_24hr − PHH_144 − Hill
BMC: 252.81 μM, BMDL: 132.06 μM, BMDU: 585.7 μM
HepaRG_APAP_24hr − PHH_144 − Linear model
BMC: 1406.33 μM / 2812.65μM, p-adjust: 0.04602
HepaRG_APAP_24hr − SRXN1 (PHH_144) − Linear
BMC: 1880.86 μM, WTT p-adj: 0.10113, DEG p-adj: 3.11e−04
HepaRG_APAP_24hr − GCLM (PHH_144) − Exp 2
BMC: 2206.08 μM, WTT p-adj: 0.07923, DEG p-adj: 8.45e−03
HepaRG_APAP_24hr − PHH_144 − Linear
BMC: 1406.33 μM, BMDL: 1017.72 μM, BMDU: 2248.16 μM
Module PHH:31
PHH_APAP_24hr − PHH_31 − Hill model
BMC: 162.66 μM / 396.58μM, p-adjust: 0.00151
PHH_APAP_24hr − SCPEP1 (PHH_31) − Exp 4
BMC: 129.71 μM, WTT p-adj: 0.00372, DEG p-adj: 1.31e−09
PHH_APAP_24hr − EML6 (PHH_31) − Exp 4
BMC: 449.39 μM, WTT p-adj: 0.00372, DEG p-adj: 2.02e−03
PHH_APAP_24hr − PHH_31 − Hill
BMC: 162.66 μM, BMDL: 80.24 μM, BMDU: 405.6 μM
HepaRG_APAP_24hr − PHH_31 − Linear model
BMC: 1284.83 μM / 2569.65μM, p-adjust: 0.05548
HepaRG_APAP_24hr − SCPEP1 (PHH_31) − Linear
BMC: 1859.48 μM, WTT p-adj: 0.19439, DEG p-adj: 6.01e−02
HepaRG_APAP_24hr − EML6 (PHH_31) − Exp 2
BMC: 5621.02 μM, WTT p-adj: 0.87633, DEG p-adj: 9.54e−01
HepaRG_APAP_24hr − PHH_31 − Linear
BMC: 1284.83 μM, BMDL: 942.8 μM, BMDU: 1990.67 μM
Module PHH:11
PHH_APAP_24hr − PHH_11 − Hill model
BMC: 134.28 μM / 274.12μM, p-adjust: 0.00151
PHH_APAP_24hr − CPS1 (PHH_11) − Exp 4
BMC: 101.02 μM, WTT p-adj: 0.00372, DEG p-adj: 2.75e−37
PHH_APAP_24hr − SLC5A9 (PHH_11) − Linear
BMC: 415.63 μM, WTT p-adj: 0.00372, DEG p-adj: 3.13e−07
PHH_APAP_24hr − PHH_11 − Hill
BMC: 134.28 μM, BMDL: 103.08 μM, BMDU: 241.46 μM
HepaRG_APAP_24hr − PHH_11 − Linear model
BMC: 3968.52 μM / 7937.03μM, p-adjust: 0.75553
HepaRG_APAP_24hr − CPS1 (PHH_11) − Exp 2
BMC: 1455.13 μM, WTT p-adj: 0.16363, DEG p-adj: 6.68e−04
HepaRG_APAP_24hr − SLC5A9 (PHH_11) − Linear
BMC: 2755.29 μM, WTT p-adj: 0.53695, DEG p-adj: 1.52e−01
HepaRG_APAP_24hr − PHH_11 − Linear
BMC: 3968.52 μM, BMDL: 2128.13 μM, BMDU: 29046.3 μM
Module PHH:23
PHH_APAP_24hr − PHH_23 − Hill model
BMC: 84.78 μM / 177.36μM, p-adjust: 0.00151
PHH_APAP_24hr − AFM (PHH_23) − Exp 4
BMC: 129.12 μM, WTT p-adj: 0.00372, DEG p-adj: 5.25e−19
PHH_APAP_24hr − ARG1 (PHH_23) − Exp 4
BMC: 73 μM, WTT p-adj: 0.00372, DEG p-adj: 8.56e−19
PHH_APAP_24hr − PHH_23 − Hill
BMC: 84.78 μM, BMDL: 63.45 μM, BMDU: 149.06 μM
HepaRG_APAP_24hr − PHH_23 − Linear model
BMC: 4101.01 μM / 8202.02μM, p-adjust: 0.81745
HepaRG_APAP_24hr − AFM (PHH_23) − Linear
BMC: 2715.76 μM, WTT p-adj: 0.48771, DEG p-adj: 1.32e−01
HepaRG_APAP_24hr − ARG1 (PHH_23) − Linear
BMC: 2225.79 μM, WTT p-adj: 0.22299, DEG p-adj: 5.83e−04
HepaRG_APAP_24hr − PHH_23 − Linear
BMC: 4101.01 μM, BMDL: 2168.18 μM, BMDU: 37322 μM
Module PHH:261
PHH_APAP_48hr − PHH_261 − Hill model
BMC: 173.38 μM / 363.13μM, p-adjust: 0.00185
PHH_APAP_48hr − KRT80 (PHH_261) − Exp 2
BMC: 267.7 μM, WTT p-adj: 0.00349, DEG p-adj: 2.06e−27
PHH_APAP_48hr − SCGN (PHH_261) − Exp 2
BMC: 495.21 μM, WTT p-adj: 0.00349, DEG p-adj: 5.75e−13
PHH_APAP_48hr − PHH_261 − Hill
BMC: 173.38 μM, BMDL: 125.04 μM, BMDU: 332.17 μM
HepaRG_APAP_48hr − PHH_261 − Linear model
BMC: 604.12 μM / 1208.25μM, p-adjust: 0.00676
HepaRG_APAP_48hr − KRT80 (PHH_261) − Linear
BMC: 1033.28 μM, WTT p-adj: 0.01009, DEG p-adj: 2.89e−04
HepaRG_APAP_48hr − SCGN (PHH_261) − Linear
BMC: 1447.71 μM, WTT p-adj: 0.02624, DEG p-adj: 1.13e−03
HepaRG_APAP_48hr − PHH_261 − Linear
BMC: 604.12 μM, BMDL: 473.48 μM, BMDU: 817.82 μM
Module EGS
Gene log2 expression
Concentration (μM)

**Figure S5b**

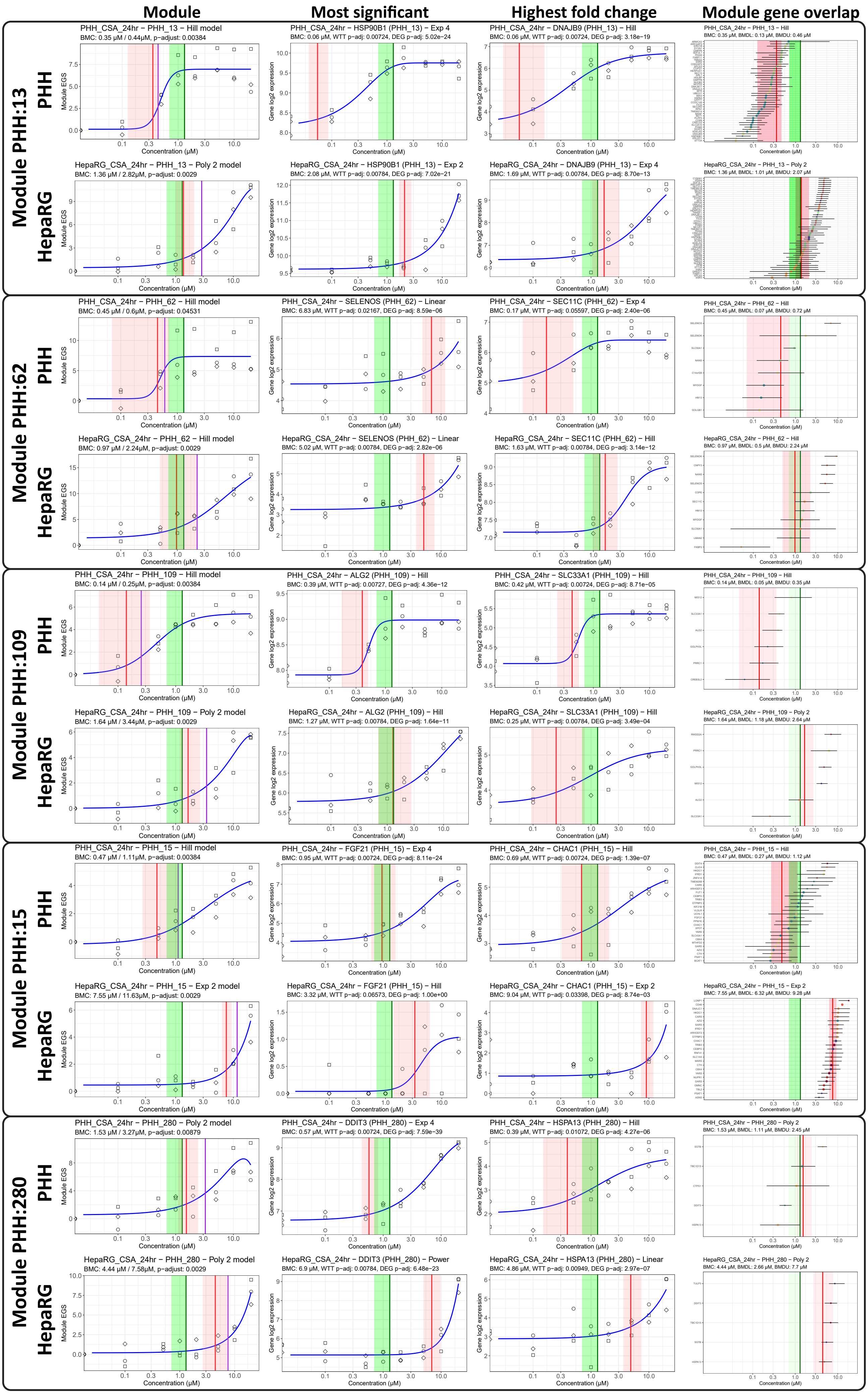

Module
Most significant
Highest fold change
Module gene overlap
Module PHH:13
Module PHH:62
Module PHH:109
Module PHH:15
Module PHH:280
PHH
HepaRG
PHH_CSA_24hr – PHH_13 – Hill model
BMC: 0.35 μM / 0.44μM, p-adjust: 0.00384
PHH_CSA_24hr – HSP90B1 (PHH_13) – Exp 4
BMC: 0.06 μM, WTT p-adj: 0.00724, DEG p-adj: 5.02e–24
PHH_CSA_24hr – DNAJB9 (PHH_13) – Hill
BMC: 0.06 μM, WTT p-adj: 0.00724, DEG p-adj: 3.18e–19
PHH_CSA_24hr – PHH_13 – Hill
BMC: 0.35 μM, BMDL: 0.13 μM, BMDU: 0.46 μM
HepaRG_CSA_24hr – PHH_13 – Poly 2 model
BMC: 1.36 μM / 2.82μM, p-adjust: 0.0029
HepaRG_CSA_24hr – HSP90B1 (PHH_13) – Exp 2
BMC: 2.08 μM, WTT p-adj: 0.00784, DEG p-adj: 7.02e–21
HepaRG_CSA_24hr – DNAJB9 (PHH_13) – Exp 4
BMC: 1.69 μM, WTT p-adj: 0.00784, DEG p-adj: 8.70e–13
HepaRG_CSA_24hr – PHH_13 – Poly 2
BMC: 1.36 μM, BMDL: 1.01 μM, BMDU: 2.07 μM
PHH_CSA_24hr – PHH_62 – Hill model
BMC: 0.45 μM / 0.6μM, p-adjust: 0.04531
PHH_CSA_24hr – SELENOS (PHH_62) – Linear
BMC: 6.83 μM, WTT p-adj: 0.02167, DEG p-adj: 8.59e–06
PHH_CSA_24hr – SEC11C (PHH_62) – Exp 4
BMC: 0.17 μM, WTT p-adj: 0.05597, DEG p-adj: 2.40e–06
PHH_CSA_24hr – PHH_62 – Hill
BMC: 0.45 μM, BMDL: 0.07 μM, BMDU: 0.72 μM
HepaRG_CSA_24hr – PHH_62 – Hill model
BMC: 0.97 μM / 2.24μM, p-adjust: 0.0029
HepaRG_CSA_24hr – SELENOS (PHH_62) – Linear
BMC: 5.02 μM, WTT p-adj: 0.00784, DEG p-adj: 2.82e–06
HepaRG_CSA_24hr – SEC11C (PHH_62) – Hill
BMC: 1.63 μM, WTT p-adj: 0.00784, DEG p-adj: 3.14e–12
HepaRG_CSA_24hr – PHH_62 – Hill
BMC: 0.97 μM, BMDL: 0.5 μM, BMDU: 2.24 μM
PHH_CSA_24hr – PHH_109 – Hill model
BMC: 0.14 μM / 0.25μM, p-adjust: 0.00384
PHH_CSA_24hr – ALG2 (PHH_109) – Hill
BMC: 0.39 μM, WTT p-adj: 0.00727, DEG p-adj: 4.36e–12
PHH_CSA_24hr – SLC33A1 (PHH_109) – Hill
BMC: 0.42 μM, WTT p-adj: 0.00724, DEG p-adj: 8.71e–05
PHH_CSA_24hr – PHH_109 – Hill
BMC: 0.14 μM, BMDL: 0.05 μM, BMDU: 0.35 μM
HepaRG_CSA_24hr – PHH_109 – Poly 2 model
BMC: 1.64 μM / 3.44μM, p-adjust: 0.0029
HepaRG_CSA_24hr – ALG2 (PHH_109) – Hill
BMC: 1.27 μM, WTT p-adj: 0.00784, DEG p-adj: 1.64e–11
HepaRG_CSA_24hr – SLC33A1 (PHH_109) – Hill
BMC: 0.25 μM, WTT p-adj: 0.00784, DEG p-adj: 3.49e–04
HepaRG_CSA_24hr – PHH_109 – Poly 2
BMC: 1.64 μM, BMDL: 1.18 μM, BMDU: 2.64 μM
PHH_CSA_24hr – PHH_15 – Hill model
BMC: 0.47 μM / 1.11μM, p-adjust: 0.00384
PHH_CSA_24hr – FGF21 (PHH_15) – Exp 4
BMC: 0.95 μM, WTT p-adj: 0.00724, DEG p-adj: 8.11e–24
PHH_CSA_24hr – CHAC1 (PHH_15) – Hill
BMC: 0.69 μM, WTT p-adj: 0.00724, DEG p-adj: 1.39e–07
PHH_CSA_24hr – PHH_15 – Hill
BMC: 0.47 μM, BMDL: 0.27 μM, BMDU: 1.12 μM
HepaRG_CSA_24hr – PHH_15 – Exp 2 model
BMC: 7.55 μM / 11.63μM, p-adjust: 0.0029
HepaRG_CSA_24hr – FGF21 (PHH_15) – Hill
BMC: 3.32 μM, WTT p-adj: 0.06573, DEG p-adj: 1.00e+00
HepaRG_CSA_24hr – CHAC1 (PHH_15) – Exp 2
BMC: 9.04 μM, WTT p-adj: 0.03398, DEG p-adj: 8.74e–03
HepaRG_CSA_24hr – PHH_15 – Exp 2
BMC: 7.55 μM, BMDL: 6.32 μM, BMDU: 9.28 μM
PHH_CSA_24hr – PHH_280 – Poly 2 model
BMC: 1.53 μM / 3.27μM, p-adjust: 0.00879
PHH_CSA_24hr – DDIT3 (PHH_280) – Exp 4
BMC: 0.57 μM, WTT p-adj: 0.00724, DEG p-adj: 7.59e–39
PHH_CSA_24hr – HSPA13 (PHH_280) – Hill
BMC: 0.39 μM, WTT p-adj: 0.01072, DEG p-adj: 4.27e–06
PHH_CSA_24hr – PHH_280 – Poly 2
BMC: 1.53 μM, BMDL: 1.11 μM, BMDU: 2.45 μM
HepaRG_CSA_24hr – PHH_280 – Poly 2 model
BMC: 4.44 μM / 7.58μM, p-adjust: 0.0029
HepaRG_CSA_24hr – DDIT3 (PHH_280) – Power
BMC: 6.9 μM, WTT p-adj: 0.00784, DEG p-adj: 6.48e–23
HepaRG_CSA_24hr – HSPA13 (PHH_280) – Linear
BMC: 4.86 μM, WTT p-adj: 0.00949, DEG p-adj: 2.97e–07
HepaRG_CSA_24hr – PHH_280 – Poly 2
BMC: 4.44 μM, BMDL: 2.66 μM, BMDU: 7.7 μM
Module EGS
Gene log2 expression
Concentration (μM)

### Figure S5c

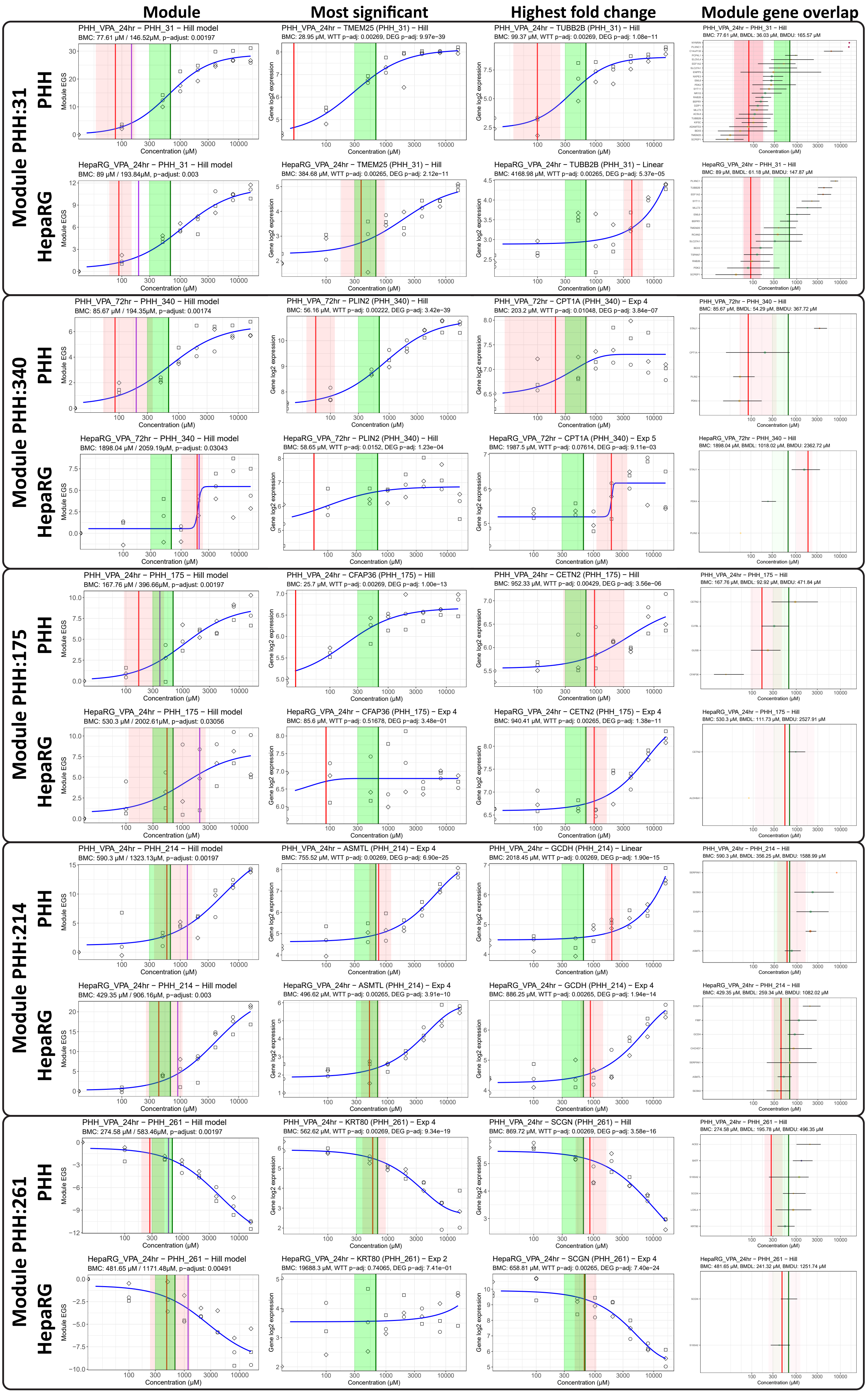

Module
Most significant
Highest fold change
Module gene overlap
Module PHH:31
PHH
HepaRG
PHH_VPA_24hr – PHH_31 – Hill model
BMC: 77.61 μM / 146.52μM, p-adjust: 0.00197
PHH_VPA_24hr – TMEM25 (PHH_31) – Hill
BMC: 28.95 μM, WTT p-adj: 0.00269, DEG p-adj: 9.97e–39
PHH_VPA_24hr – TUBB2B (PHH_31) – Hill
BMC: 99.37 μM, WTT p-adj: 0.00269, DEG p-adj: 1.08e–11
PHH_VPA_24hr – PHH_31 – Hill
BMC: 77.61 μM, BMDL: 36.03 μM, BMDU: 165.57 μM
HepaRG_VPA_24hr – PHH_31 – Hill model
BMC: 89 μM / 193.84μM, p-adjust: 0.003
HepaRG_VPA_24hr – TMEM25 (PHH_31) – Hill
BMC: 384.68 μM, WTT p-adj: 0.00265, DEG p-adj: 2.12e–11
HepaRG_VPA_24hr – TUBB2B (PHH_31) – Linear
BMC: 4168.98 μM, WTT p-adj: 0.00265, DEG p-adj: 5.37e–05
HepaRG_VPA_24hr – PHH_31 – Hill
BMC: 89 μM, BMDL: 61.18 μM, BMDU: 147.87 μM
Module PHH:340
PHH_VPA_72hr – PHH_340 – Hill model
BMC: 85.67 μM / 194.35μM, p-adjust: 0.00174
PHH_VPA_72hr – PLIN2 (PHH_340) – Hill
BMC: 56.16 μM, WTT p-adj: 0.00222, DEG p-adj: 3.42e–39
PHH_VPA_72hr – CPT1A (PHH_340) – Exp 4
BMC: 203.2 μM, WTT p-adj: 0.01048, DEG p-adj: 3.84e–07
PHH_VPA_72hr – PHH_340 – Hill
BMC: 85.67 μM, BMDL: 54.29 μM, BMDU: 367.72 μM
HepaRG_VPA_72hr – PHH_340 – Hill model
BMC: 1898.04 μM / 2059.19μM, p-adjust: 0.03043
HepaRG_VPA_72hr – PLIN2 (PHH_340) – Hill
BMC: 58.65 μM, WTT p-adj: 0.0152, DEG p-adj: 1.23e–04
HepaRG_VPA_72hr – CPT1A (PHH_340) – Exp 5
BMC: 1987.5 μM, WTT p-adj: 0.07614, DEG p-adj: 9.11e–03
HepaRG_VPA_72hr – PHH_340 – Hill
BMC: 1898.04 μM, BMDL: 1018.02 μM, BMDU: 2362.72 μM
Module PHH:175
PHH_VPA_24hr – PHH_175 – Hill model
BMC: 167.76 μM / 396.66μM, p-adjust: 0.00197
PHH_VPA_24hr – CFAP36 (PHH_175) – Hill
BMC: 25.7 μM, WTT p-adj: 0.00269, DEG p-adj: 1.00e–13
PHH_VPA_24hr – CETN2 (PHH_175) – Hill
BMC: 952.33 μM, WTT p-adj: 0.00429, DEG p-adj: 3.56e–06
PHH_VPA_24hr – PHH_175 – Hill
BMC: 167.76 μM, BMDL: 92.92 μM, BMDU: 471.84 μM
HepaRG_VPA_24hr – PHH_175 – Hill model
BMC: 530.3 μM / 2002.61μM, p-adjust: 0.03056
HepaRG_VPA_24hr – CFAP36 (PHH_175) – Exp 4
BMC: 85.6 μM, WTT p-adj: 0.51678, DEG p-adj: 3.48e–01
HepaRG_VPA_24hr – CETN2 (PHH_175) – Exp 4
BMC: 940.41 μM, WTT p-adj: 0.00265, DEG p-adj: 1.38e–11
HepaRG_VPA_24hr – PHH_175 – Hill
BMC: 530.3 μM, BMDL: 111.73 μM, BMDU: 2527.91 μM
Module PHH:214
PHH_VPA_24hr – PHH_214 – Hill model
BMC: 590.3 μM / 1323.13μM, p-adjust: 0.00197
PHH_VPA_24hr – ASMTL (PHH_214) – Exp 4
BMC: 755.52 μM, WTT p-adj: 0.00269, DEG p-adj: 6.90e–25
PHH_VPA_24hr – GCDH (PHH_214) – Linear
BMC: 2018.45 μM, WTT p-adj: 0.00269, DEG p-adj: 1.90e–15
PHH_VPA_24hr – PHH_214 – Hill
BMC: 590.3 μM, BMDL: 356.25 μM, BMDU: 1588.99 μM
HepaRG_VPA_24hr – PHH_214 – Hill model
BMC: 429.35 μM / 906.16μM, p-adjust: 0.003
HepaRG_VPA_24hr – ASMTL (PHH_214) – Exp 4
BMC: 496.62 μM, WTT p-adj: 0.00265, DEG p-adj: 3.91e–10
HepaRG_VPA_24hr – GCDH (PHH_214) – Exp 4
BMC: 886.25 μM, WTT p-adj: 0.00265, DEG p-adj: 1.94e–14
HepaRG_VPA_24hr – PHH_214 – Hill
BMC: 429.35 μM, BMDL: 259.34 μM, BMDU: 1082.02 μM
Module PHH:261
PHH_VPA_24hr – PHH_261 – Hill model
BMC: 274.58 μM / 583.46μM, p-adjust: 0.00197
PHH_VPA_24hr – KRT80 (PHH_261) – Exp 4
BMC: 562.62 μM, WTT p-adj: 0.00269, DEG p-adj: 9.34e–19
PHH_VPA_24hr – SCGN (PHH_261) – Hill
BMC: 869.72 μM, WTT p-adj: 0.00269, DEG p-adj: 3.58e–16
PHH_VPA_24hr – PHH_261 – Hill
BMC: 274.58 μM, BMDL: 195.78 μM, BMDU: 496.35 μM
HepaRG_VPA_24hr – PHH_261 – Hill model
BMC: 481.65 μM / 1171.48μM, p-adjust: 0.00491
HepaRG_VPA_24hr – KRT80 (PHH_261) – Exp 2
BMC: 19688.3 μM, WTT p-adj: 0.74065, DEG p-adj: 7.41e–01
HepaRG_VPA_24hr – SCGN (PHH_261) – Exp 4
BMC: 658.81 μM, WTT p-adj: 0.00265, DEG p-adj: 7.40e–24
HepaRG_VPA_24hr – PHH_261 – Hill
BMC: 481.65 μM, BMDL: 241.32 μM, BMDU: 1251.74 μM
Module EGS
Gene log2 expression
Concentration (μM)

**Figure S6**

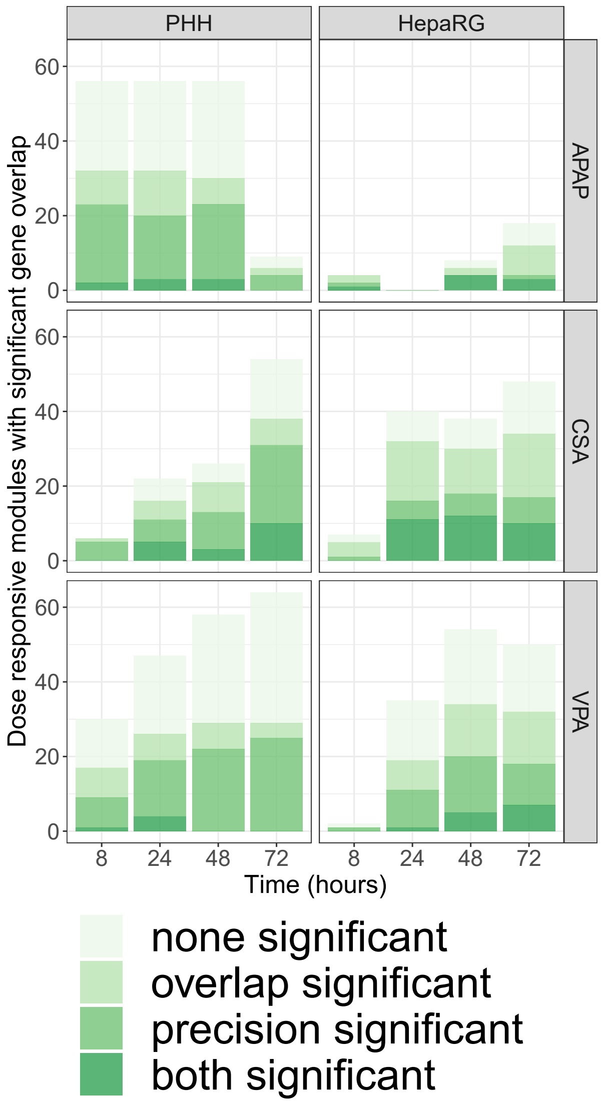
PHH
HepaRG
APAP
CSA
VPA
Dose responsive modules with significant gene overlap
60
40
20
0
8
24
48
72
Time (hours)
none significant
overlap significant
precision significant
both significant

**Figure S7**

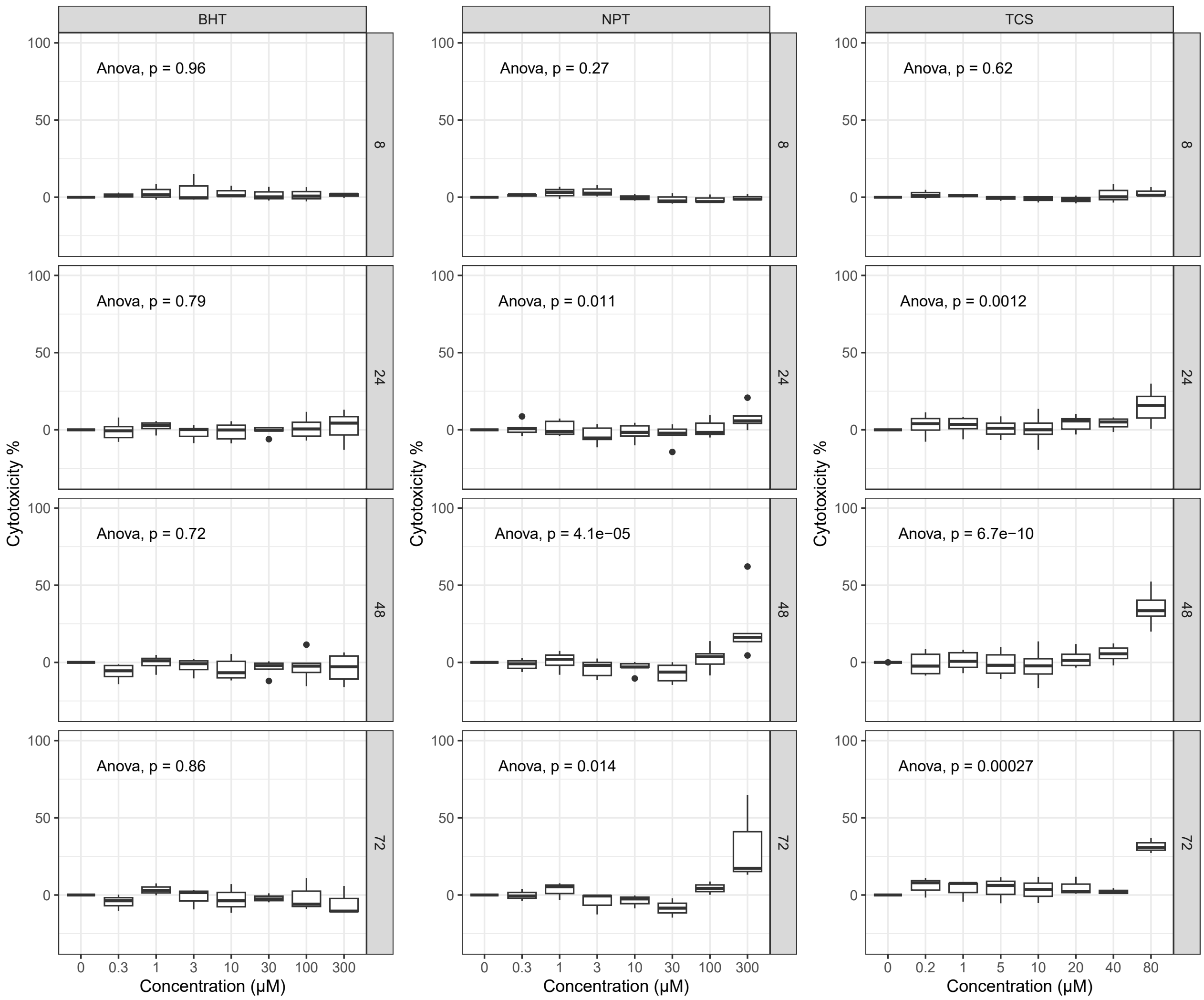
BHT
NPT
TCS
Anova, p = 0.96
Anova, p = 0.27
Anova, p = 0.62
Anova, p = 0.79
Anova, p = 0.011
Anova, p = 0.0012
Anova, p = 0.72
Anova, p = 4.1e−05
Anova, p = 6.7e−10
Anova, p = 0.86
Anova, p = 0.014
Anova, p = 0.00027
8
24
48
72
Cytotoxicity %
Concentration (μM)
0 0.3 1 3 10 30 100 300
0 0.2 1 5 10 20 40 80

#### Figure S8

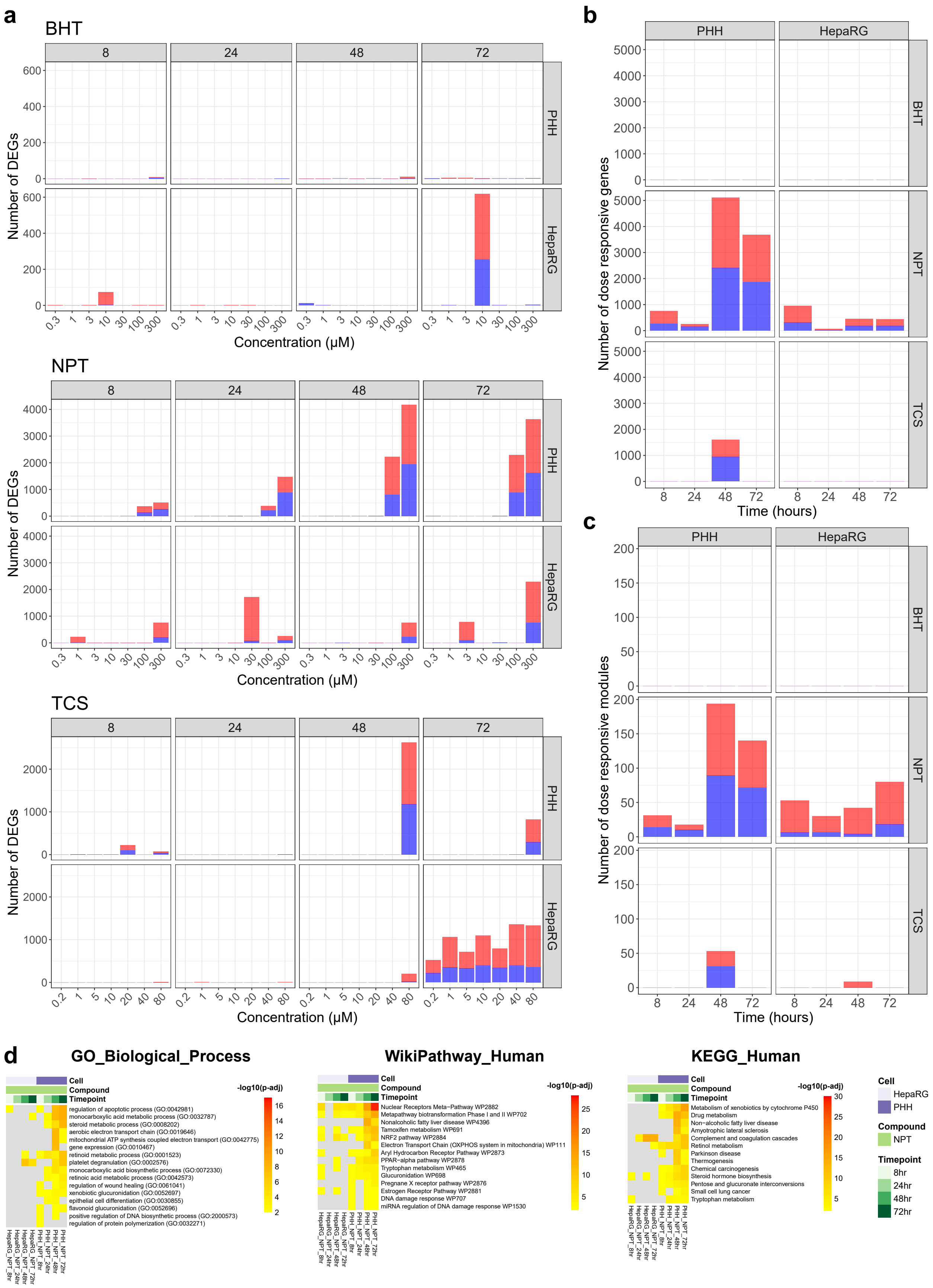
a
BHT
8
24
48
72
PHH
HepaRG
Number of DEGs
Concentration (μM)
NPT
TCS
b
PHH
HepaRG
BHT
NPT
TCS
Number of dose responsive genes
Time (hours)
c
Number of dose responsive modules
d
GO_Biological_Process
Cell
Compound
Timepoint
-log10(p-adj)
regulation of apoptotic process (GO:0042981)
monocarboxylic acid metabolic process (GO:0032787)
steroid metabolic process (GO:0008202)
aerobic electron transport chain (GO:0019646)
mitochondrial ATP synthesis coupled electron transport (GO:0042775)
gene expression (GO:0010467)
retinoid metabolic process (GO:0001523)
platelet degranulation (GO:0002576)
monocarboxylic acid biosynthetic process (GO:0072330)
retinoic acid metabolic process (GO:0042573)
regulation of wound healing (GO:0061041)
xenobiotic glucuronidation (GO:0052697)
epithelial cell differentiation (GO:0030855)
flavonoid glucuronidation (GO:0052696)
positive regulation of DNA biosynthetic process (GO:2000573)
regulation of protein polymerization (GO:0032271)
HepaRG_NPT_8hr
HepaRG_NPT_24hr
HepaRG_NPT_48hr
HepaRG_NPT_72hr
PHH_NPT_8hr
PHH_NPT_24hr
PHH_NPT_48hr
PHH_NPT_72hr
WikiPathway_Human
Nuclear Receptors Meta−Pathway WP2882
Metapathway biotransformation Phase I and II WP702
Nonalcoholic fatty liver disease WP4396
Tamoxifen metabolism WP691
NRF2 pathway WP2884
Electron Transport Chain (OXPHOS system in mitochondria) WP111
Aryl Hydrocarbon Receptor Pathway WP2873
PPAR−alpha pathway WP2878
Tryptophan metabolism WP465
Glucuronidation WP698
Pregnane X receptor pathway WP2876
Estrogen Receptor Pathway WP2881
DNA damage response WP707
miRNA regulation of DNA damage response WP1530
KEGG_Human
Metabolism of xenobiotics by cytochrome P450
Drug metabolism
Non−alcoholic fatty liver disease
Amyotrophic lateral sclerosis
Complement and coagulation cascades
Retinol metabolism
Parkinson disease
Thermogenesis
Chemical carcinogenesis
Steroid hormone biosynthesis
Pentose and glucuronate interconversions
Small cell lung cancer
Tryptophan metabolism
Cell
HepaRG
PHH
Compound
NPT
Timepoint
8hr
24hr
48hr
72hr

### Figure S9

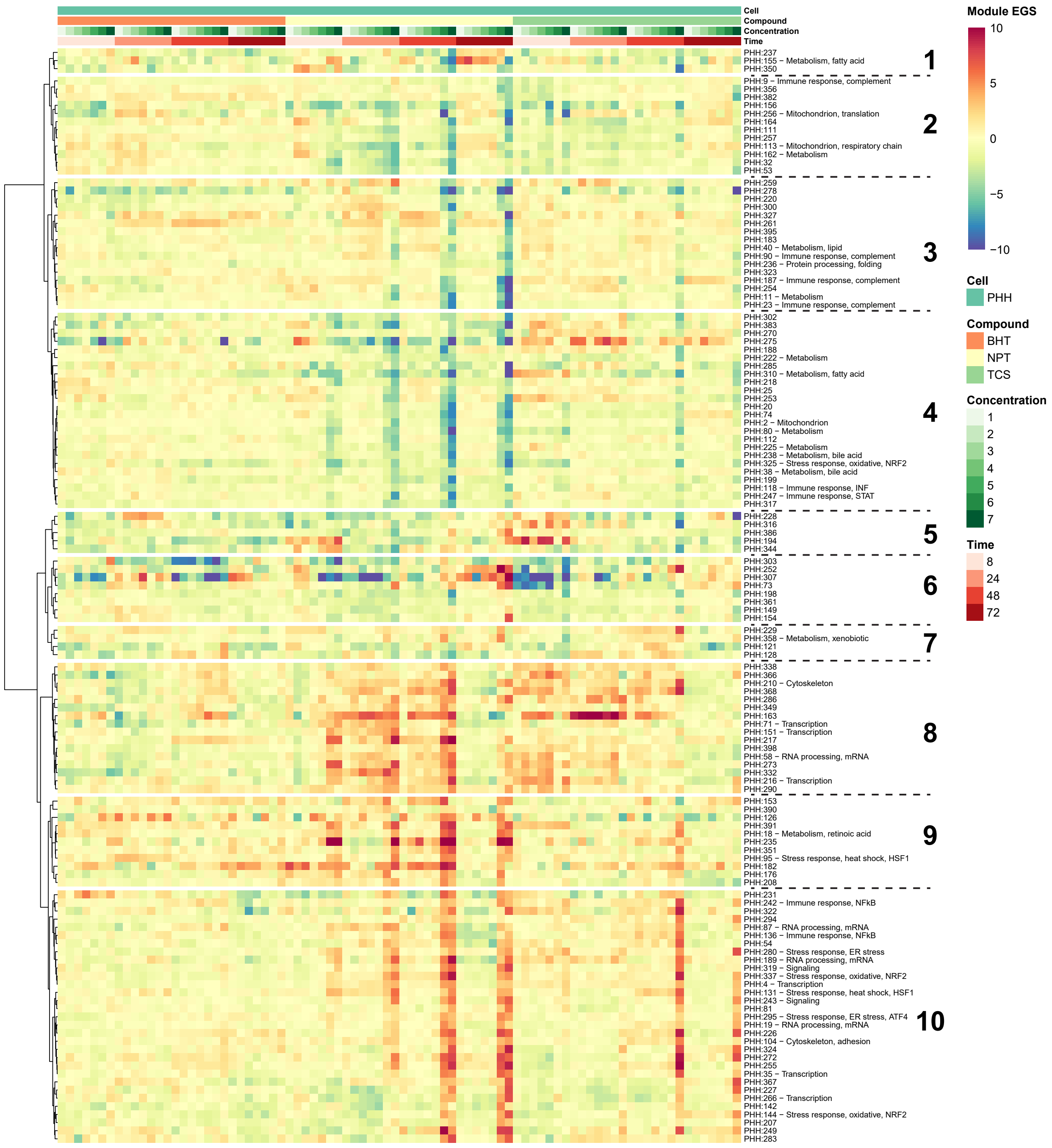

Cell
Compound
Concentration
Time
PHH:237
PHH:155 − Metabolism, fatty acid
PHH:350
1
PHH:9 − Immune response, complement
PHH:356
PHH:382
PHH:156
PHH:256 − Mitochondrion, translation
PHH:164
PHH:111
PHH:257
PHH:113 − Mitochondrion, respiratory chain
PHH:162 − Metabolism
PHH:32
PHH:53
2
PHH:259
PHH:278
PHH:220
PHH:300
PHH:327
PHH:261
PHH:395
PHH:183
PHH:40 − Metabolism, lipid
PHH:90 − Immune response, complement
PHH:236 − Protein processing, folding
PHH:323
PHH:187 − Immune response, complement
PHH:254
PHH:11 − Metabolism
PHH:23 − Immune response, complement
3
PHH:302
PHH:383
PHH:270
PHH:275
PHH:188
PHH:222 − Metabolism
PHH:285
PHH:310 − Metabolism, fatty acid
PHH:218
PHH:25
PHH:253
PHH:20
PHH:74
PHH:2 − Mitochondrion
PHH:80 − Metabolism
PHH:112
PHH:225 − Metabolism
PHH:238 − Metabolism, bile acid
PHH:325 − Stress response, oxidative, NRF2
PHH:38 − Metabolism, bile acid
PHH:199
PHH:118 − Immune response, INF
PHH:247 − Immune response, STAT
PHH:317
4
PHH:228
PHH:316
PHH:386
PHH:194
PHH:344
5
PHH:303
PHH:252
PHH:307
PHH:73
PHH:198
PHH:361
PHH:149
PHH:154
6
PHH:229
PHH:358 − Metabolism, xenobiotic
PHH:121
PHH:128
7
PHH:338
PHH:366
PHH:210 − Cytoskeleton
PHH:368
PHH:286
PHH:349
PHH:163
PHH:71 − Transcription
PHH:151 − Transcription
PHH:217
PHH:398
PHH:58 − RNA processing, mRNA
PHH:273
PHH:332
PHH:216 − Transcription
PHH:290
8
PHH:153
PHH:390
PHH:126
PHH:391
PHH:18 − Metabolism, retinoic acid
PHH:235
PHH:351
PHH:95 − Stress response, heat shock, HSF1
PHH:182
PHH:176
PHH:208
9
PHH:231
PHH:242 − Immune response, NFkB
PHH:322
PHH:294
PHH:87 − RNA processing, mRNA
PHH:136 − Immune response, NFkB
PHH:54
PHH:280 − Stress response, ER stress
PHH:189 − RNA processing, mRNA
PHH:319 − Signaling
PHH:337 − Stress response, oxidative, NRF2
PHH:4 − Transcription
PHH:131 − Stress response, heat shock, HSF1
PHH:243 − Signaling
PHH:81
PHH:295 − Stress response, ER stress, ATF4
PHH:19 − RNA processing, mRNA
PHH:226
PHH:104 − Cytoskeleton, adhesion
PHH:324
PHH:272
PHH:255
PHH:35 − Transcription
PHH:367
PHH:227
PHH:266 − Transcription
PHH:142
PHH:144 − Stress response, oxidative, NRF2
PHH:207
PHH:249
PHH:283
10
Module EGS
10
5
0
−5
−10
Cell
PHH
Compound
BHT
NPT
TCS
Concentration
1
2
3
4
5
6
7
Time
8
24
48
72

**Figure S10a**

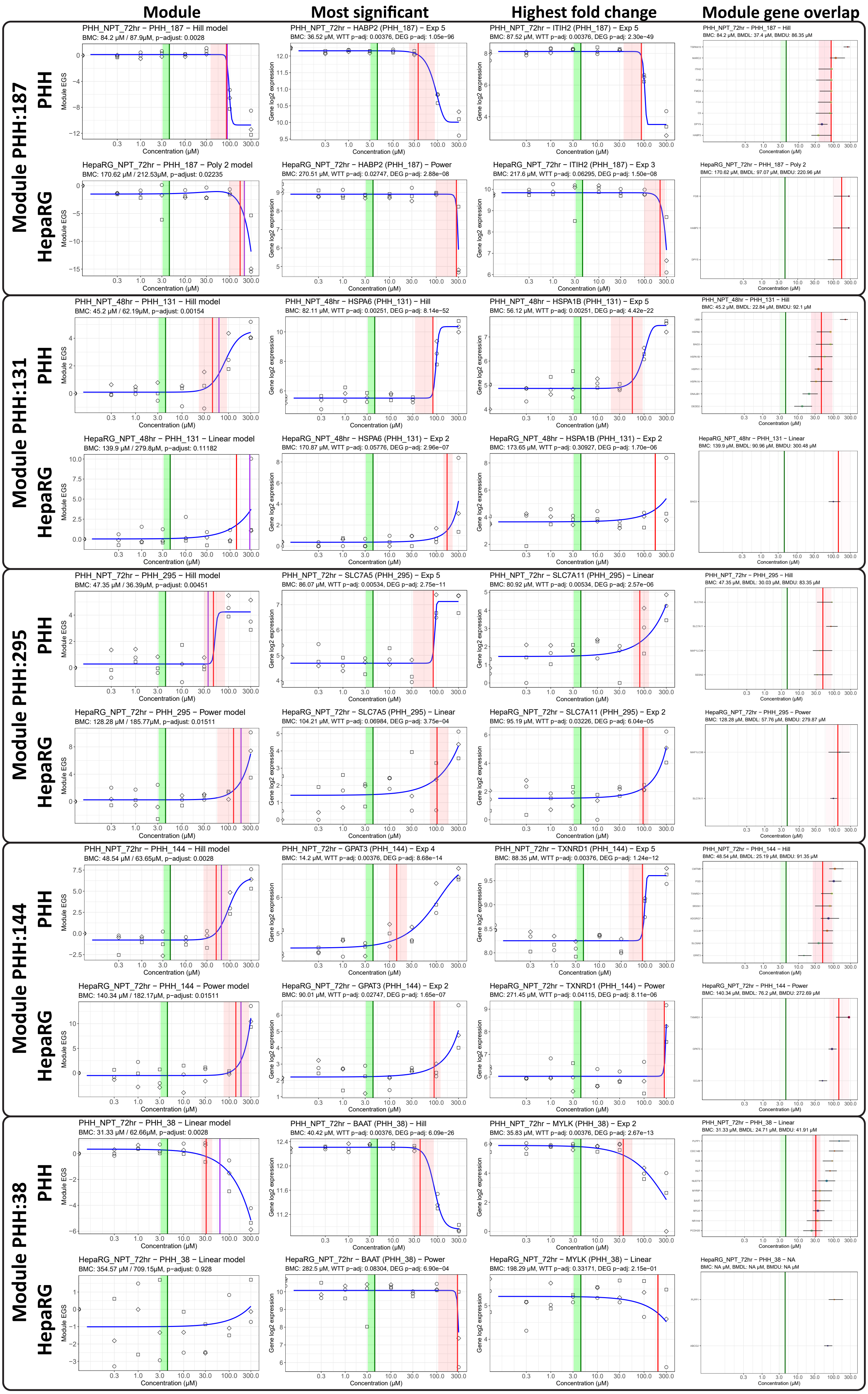

Module
Most significant
Highest fold change
Module gene overlap
Module PHH:187
PHH
HepaRG
PHH_NPT_72hr – PHH_187 – Hill model
BMC: 84.2 μM / 87.9μM, p-adjust: 0.0028
PHH_NPT_72hr – HABP2 (PHH_187) – Exp 5
BMC: 36.52 μM, WTT p-adj: 0.00376, DEG p-adj: 1.05e−96
PHH_NPT_72hr – ITIH2 (PHH_187) – Exp 5
BMC: 87.52 μM, WTT p-adj: 0.00376, DEG p-adj: 2.30e−49
PHH_NPT_72hr – PHH_187 – Hill
BMC: 84.2 μM, BMDL: 37.4 μM, BMDU: 86.35 μM
HepaRG_NPT_72hr – PHH_187 – Poly 2 model
BMC: 170.62 μM / 212.53μM, p-adjust: 0.02235
HepaRG_NPT_72hr – HABP2 (PHH_187) – Power
BMC: 270.51 μM, WTT p-adj: 0.02747, DEG p-adj: 2.88e−08
HepaRG_NPT_72hr – ITIH2 (PHH_187) – Exp 3
BMC: 217.6 μM, WTT p-adj: 0.06295, DEG p-adj: 1.50e−08
HepaRG_NPT_72hr – PHH_187 – Poly 2
BMC: 170.62 μM, BMDL: 97.07 μM, BMDU: 220.96 μM
Module PHH:131
PHH_NPT_48hr – PHH_131 – Hill model
BMC: 45.2 μM / 62.19μM, p-adjust: 0.00154
PHH_NPT_48hr – HSPA6 (PHH_131) – Hill
BMC: 82.11 μM, WTT p-adj: 0.00251, DEG p-adj: 8.14e−52
PHH_NPT_48hr – HSPA1B (PHH_131) – Exp 5
BMC: 56.12 μM, WTT p-adj: 0.00251, DEG p-adj: 4.42e−22
PHH_NPT_48hr – PHH_131 – Hill
BMC: 45.2 μM, BMDL: 22.84 μM, BMDU: 92.1 μM
HepaRG_NPT_48hr – PHH_131 – Linear model
BMC: 139.9 μM / 279.8μM, p-adjust: 0.11182
HepaRG_NPT_48hr – HSPA6 (PHH_131) – Exp 2
BMC: 170.87 μM, WTT p-adj: 0.05776, DEG p-adj: 2.96e−07
HepaRG_NPT_48hr – HSPA1B (PHH_131) – Exp 2
BMC: 173.65 μM, WTT p-adj: 0.30927, DEG p-adj: 1.70e−06
HepaRG_NPT_48hr – PHH_131 – Linear
BMC: 139.9 μM, BMDL: 90.96 μM, BMDU: 300.48 μM
Module PHH:295
PHH_NPT_72hr – PHH_295 – Hill model
BMC: 47.35 μM / 36.39μM, p-adjust: 0.00451
PHH_NPT_72hr – SLC7A5 (PHH_295) – Exp 5
BMC: 86.07 μM, WTT p-adj: 0.00534, DEG p-adj: 2.75e−11
PHH_NPT_72hr – SLC7A11 (PHH_295) – Linear
BMC: 80.92 μM, WTT p-adj: 0.00534, DEG p-adj: 2.57e−06
PHH_NPT_72hr – PHH_295 – Hill
BMC: 47.35 μM, BMDL: 30.03 μM, BMDU: 83.35 μM
HepaRG_NPT_72hr – PHH_295 – Power model
BMC: 128.28 μM / 185.77μM, p-adjust: 0.01511
HepaRG_NPT_72hr – SLC7A5 (PHH_295) – Linear
BMC: 104.21 μM, WTT p-adj: 0.06984, DEG p-adj: 3.75e−04
HepaRG_NPT_72hr – SLC7A11 (PHH_295) – Exp 2
BMC: 95.19 μM, WTT p-adj: 0.03226, DEG p-adj: 6.04e−05
HepaRG_NPT_72hr – PHH_295 – Power
BMC: 128.28 μM, BMDL: 57.76 μM, BMDU: 279.87 μM
Module PHH:144
PHH_NPT_72hr – PHH_144 – Hill model
BMC: 48.54 μM / 63.65μM, p-adjust: 0.0028
PHH_NPT_72hr – GPAT3 (PHH_144) – Exp 4
BMC: 14.2 μM, WTT p-adj: 0.00376, DEG p-adj: 8.68e−14
PHH_NPT_72hr – TXNRD1 (PHH_144) – Exp 5
BMC: 88.35 μM, WTT p-adj: 0.00376, DEG p-adj: 1.24e−12
PHH_NPT_72hr – PHH_144 – Hill
BMC: 48.54 μM, BMDL: 25.19 μM, BMDU: 91.35 μM
HepaRG_NPT_72hr – PHH_144 – Power model
BMC: 140.34 μM / 182.17μM, p-adjust: 0.01511
HepaRG_NPT_72hr – GPAT3 (PHH_144) – Exp 2
BMC: 90.01 μM, WTT p-adj: 0.02747, DEG p-adj: 1.65e−07
HepaRG_NPT_72hr – TXNRD1 (PHH_144) – Power
BMC: 271.45 μM, WTT p-adj: 0.04115, DEG p-adj: 8.11e−06
HepaRG_NPT_72hr – PHH_144 – Power
BMC: 140.34 μM, BMDL: 76.2 μM, BMDU: 272.69 μM
Module PHH:38
PHH_NPT_72hr – PHH_38 – Linear model
BMC: 31.33 μM / 62.66μM, p-adjust: 0.0028
PHH_NPT_72hr – BAAT (PHH_38) – Hill
BMC: 40.42 μM, WTT p-adj: 0.00376, DEG p-adj: 6.09e−26
PHH_NPT_72hr – MYLK (PHH_38) – Exp 2
BMC: 35.83 μM, WTT p-adj: 0.00376, DEG p-adj: 2.67e−13
PHH_NPT_72hr – PHH_38 – Linear
BMC: 31.33 μM, BMDL: 24.71 μM, BMDU: 41.91 μM
HepaRG_NPT_72hr – PHH_38 – Linear model
BMC: 354.57 μM / 709.15μM, p-adjust: 0.928
HepaRG_NPT_72hr – BAAT (PHH_38) – Power
BMC: 282.5 μM, WTT p-adj: 0.08304, DEG p-adj: 6.90e−04
HepaRG_NPT_72hr – MYLK (PHH_38) – Linear
BMC: 198.29 μM, WTT p-adj: 0.33171, DEG p-adj: 2.15e−01
HepaRG_NPT_72hr – PHH_38 – NA
BMC: NA μM, BMDL: NA μM, BMDU: NA μM
Module EGS
Gene log2 expression
Concentration (μM)

**Figure S10b**

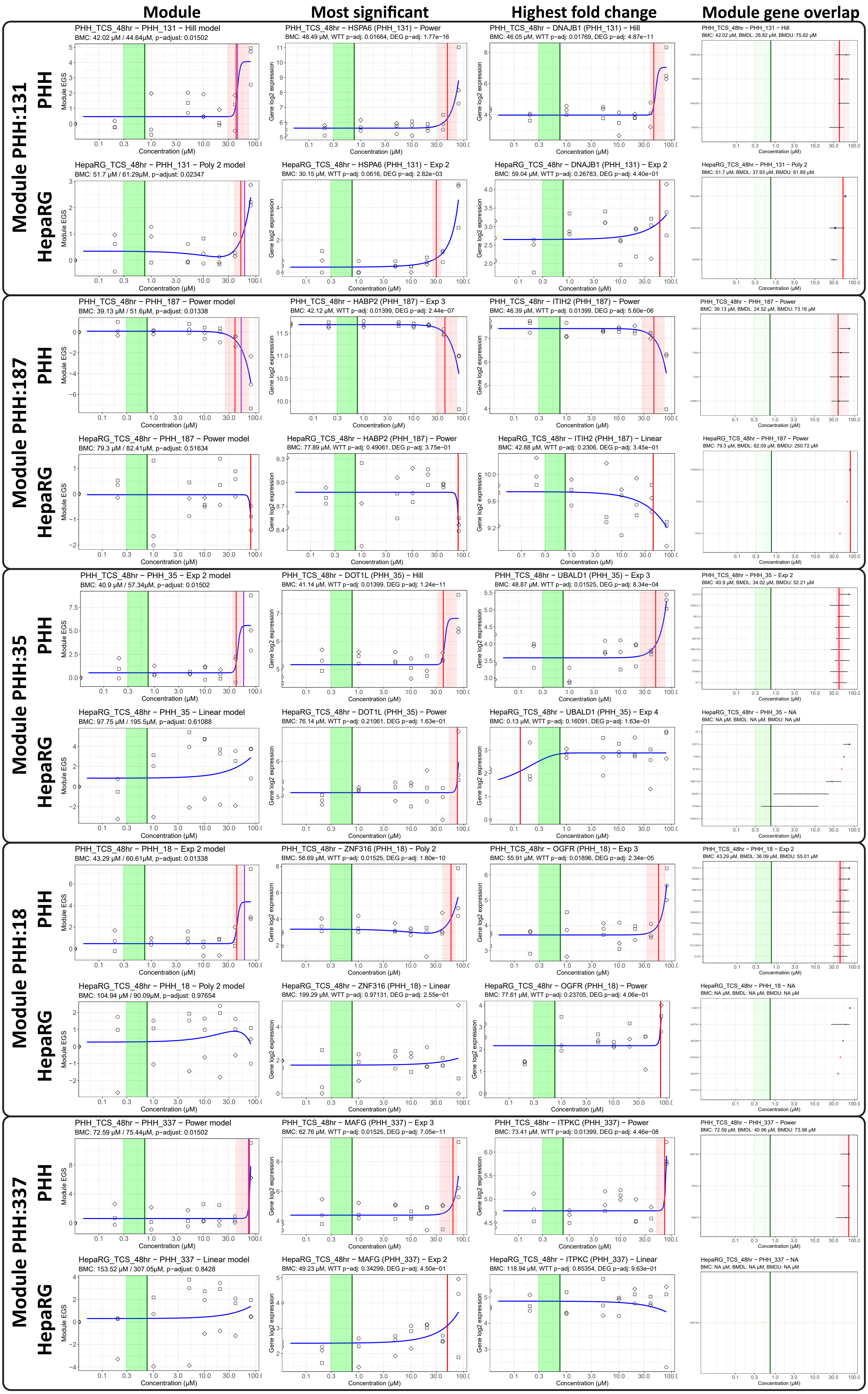
Module
Most significant
Highest fold change
Module gene overlap
Module PHH:131
PHH
HepaRG
PHH_TCS_48hr – PHH_131 – Hill model
BMC: 42.02 µM / 44.64µM, p-adjust: 0.01502
PHH_TCS_48hr – HSPA6 (PHH_131) – Power
BMC: 48.49 µM, WTT p-adj: 0.01664, DEG p-adj: 1.77e−16
PHH_TCS_48hr – DNAJB1 (PHH_131) – Hill
BMC: 46.05 µM, WTT p-adj: 0.01769, DEG p-adj: 4.87e−11
PHH_TCS_48hr – PHH_131 – Hill
BMC: 42.02 µM, BMDL: 28.82 µM, BMDU: 75.62 µM
HepaRG_TCS_48hr – PHH_131 – Poly 2 model
BMC: 51.7 µM / 61.29µM, p-adjust: 0.02347
HepaRG_TCS_48hr – HSPA6 (PHH_131) – Exp 2
BMC: 30.15 µM, WTT p-adj: 0.0616, DEG p-adj: 2.82e−03
HepaRG_TCS_48hr – DNAJB1 (PHH_131) – Exp 2
BMC: 59.04 µM, WTT p-adj: 0.26783, DEG p-adj: 4.40e−01
HepaRG_TCS_48hr – PHH_131 – Poly 2
BMC: 51.7 µM, BMDL: 37.93 µM, BMDU: 61.89 µM
Module PHH:187
PHH_TCS_48hr – PHH_187 – Power model
BMC: 39.13 µM / 51.6µM, p-adjust: 0.01338
PHH_TCS_48hr – HABP2 (PHH_187) – Exp 3
BMC: 42.12 µM, WTT p-adj: 0.01399, DEG p-adj: 2.44e−07
PHH_TCS_48hr – ITIH2 (PHH_187) – Power
BMC: 46.39 µM, WTT p-adj: 0.01399, DEG p-adj: 5.60e−06
PHH_TCS_48hr – PHH_187 – Power
BMC: 39.13 µM, BMDL: 24.52 µM, BMDU: 73.16 µM
HepaRG_TCS_48hr – PHH_187 – Power model
BMC: 79.3 µM / 82.41µM, p-adjust: 0.51634
HepaRG_TCS_48hr – HABP2 (PHH_187) – Power
BMC: 77.89 µM, WTT p-adj: 0.49061, DEG p-adj: 3.75e−01
HepaRG_TCS_48hr – ITIH2 (PHH_187) – Linear
BMC: 42.88 µM, WTT p-adj: 0.2306, DEG p-adj: 3.45e−01
HepaRG_TCS_48hr – PHH_187 – Power
BMC: 79.3 µM, BMDL: 62.09 µM, BMDU: 250.72 µM
Module PHH:35
PHH_TCS_48hr – PHH_35 – Exp 2 model
BMC: 40.9 µM / 57.34µM, p-adjust: 0.01502
PHH_TCS_48hr – DOT1L (PHH_35) – Hill
BMC: 41.14 µM, WTT p-adj: 0.01399, DEG p-adj: 1.24e−11
PHH_TCS_48hr – UBALD1 (PHH_35) – Exp 3
BMC: 48.87 µM, WTT p-adj: 0.01525, DEG p-adj: 8.34e−04
PHH_TCS_48hr – PHH_35 – Exp 2
BMC: 40.9 µM, BMDL: 34.02 µM, BMDU: 52.21 µM
HepaRG_TCS_48hr – PHH_35 – Linear model
BMC: 97.75 µM / 195.5µM, p-adjust: 0.61088
HepaRG_TCS_48hr – DOT1L (PHH_35) – Power
BMC: 76.14 µM, WTT p-adj: 0.21061, DEG p-adj: 1.63e−01
HepaRG_TCS_48hr – UBALD1 (PHH_35) – Exp 4
BMC: 0.13 µM, WTT p-adj: 0.16091, DEG p-adj: 1.63e−01
HepaRG_TCS_48hr – PHH_35 – NA
BMC: NA µM, BMDL: NA µM, BMDU: NA µM
Module PHH:18
PHH_TCS_48hr – PHH_18 – Exp 2 model
BMC: 43.29 µM / 60.61µM, p-adjust: 0.01338
PHH_TCS_48hr – ZNF316 (PHH_18) – Poly 2
BMC: 58.69 µM, WTT p-adj: 0.01525, DEG p-adj: 1.80e−10
PHH_TCS_48hr – OGFR (PHH_18) – Exp 3
BMC: 55.91 µM, WTT p-adj: 0.01896, DEG p-adj: 2.34e−05
PHH_TCS_48hr – PHH_18 – Exp 2
BMC: 43.29 µM, BMDL: 36.09 µM, BMDU: 55.01 µM
HepaRG_TCS_48hr – PHH_18 – Poly 2 model
BMC: 104.94 µM / 90.09µM, p-adjust: 0.97654
HepaRG_TCS_48hr – ZNF316 (PHH_18) – Linear
BMC: 199.29 µM, WTT p-adj: 0.97131, DEG p-adj: 2.55e−01
HepaRG_TCS_48hr – OGFR (PHH_18) – Power
BMC: 77.61 µM, WTT p-adj: 0.23705, DEG p-adj: 4.06e−01
HepaRG_TCS_48hr – PHH_18 – NA
BMC: NA µM, BMDL: NA µM, BMDU: NA µM
Module PHH:337
PHH_TCS_48hr – PHH_337 – Power model
BMC: 72.59 µM / 75.44µM, p-adjust: 0.01502
PHH_TCS_48hr – MAFG (PHH_337) – Exp 3
BMC: 62.76 µM, WTT p-adj: 0.01525, DEG p-adj: 7.05e−11
PHH_TCS_48hr – ITPKC (PHH_337) – Power
BMC: 73.41 µM, WTT p-adj: 0.01399, DEG p-adj: 4.46e−08
PHH_TCS_48hr – PHH_337 – Power
BMC: 72.59 µM, BMDL: 40.96 µM, BMDU: 73.98 µM
HepaRG_TCS_48hr – PHH_337 – Linear model
BMC: 153.52 µM / 307.05µM, p-adjust: 0.8428
HepaRG_TCS_48hr – MAFG (PHH_337) – Exp 2
BMC: 49.23 µM, WTT p-adj: 0.34299, DEG p-adj: 4.50e−01
HepaRG_TCS_48hr – ITPKC (PHH_337) – Linear
BMC: 118.94 µM, WTT p-adj: 0.85354, DEG p-adj: 9.63e−01
HepaRG_TCS_48hr – PHH_337 – NA
BMC: NA µM, BMDL: NA µM, BMDU: NA µM
Module EGS
Gene log2 expression
Concentration (µM)

#### Figure S10c

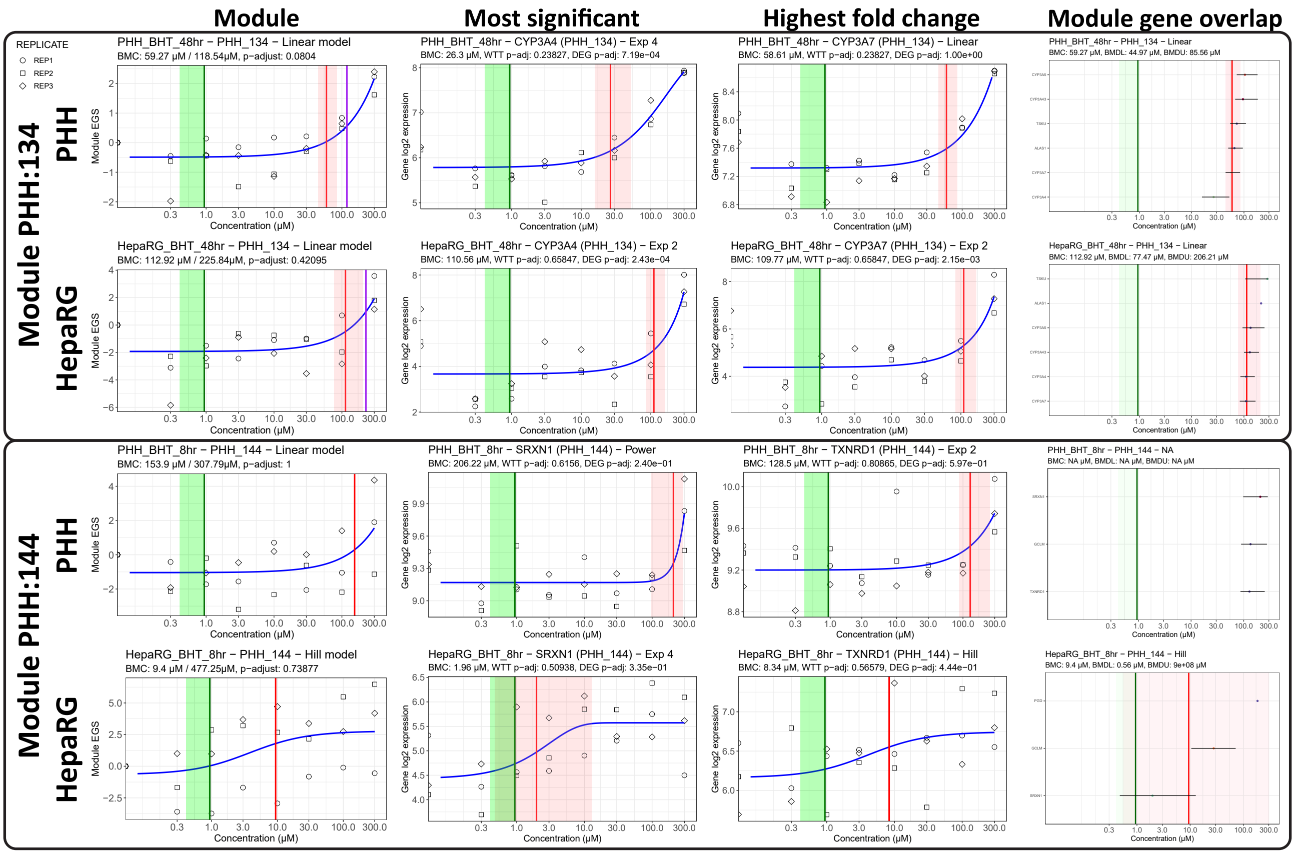

Module
Most significant
Highest fold change
Module gene overlap
REPLICATE
REP1
REP2
REP3
Module PHH:134
PHH
HepaRG
PHH_BHT_48hr − PHH_134 − Linear model
BMC: 59.27 μM / 118.54μM, p−adjust: 0.0804
PHH_BHT_48hr − CYP3A4 (PHH_134) − Exp 4
BMC: 26.3 μM, WTT p−adj: 0.23827, DEG p−adj: 7.19e−04
PHH_BHT_48hr − CYP3A7 (PHH_134) − Linear
BMC: 58.61 μM, WTT p−adj: 0.23827, DEG p−adj: 1.00e+00
PHH_BHT_48hr − PHH_134 − Linear
BMC: 59.27 μM, BMDL: 44.97 μM, BMDU: 85.56 μM
HepaRG_BHT_48hr − PHH_134 − Linear model
BMC: 112.92 μM / 225.84μM, p−adjust: 0.42095
HepaRG_BHT_48hr − CYP3A4 (PHH_134) − Exp 2
BMC: 110.56 μM, WTT p−adj: 0.65847, DEG p−adj: 2.43e−04
HepaRG_BHT_48hr − CYP3A7 (PHH_134) − Exp 2
BMC: 109.77 μM, WTT p−adj: 0.65847, DEG p−adj: 2.15e−03
HepaRG_BHT_48hr − PHH_134 − Linear
BMC: 112.92 μM, BMDL: 77.47 μM, BMDU: 206.21 μM
Module PHH:144
PHH_BHT_8hr − PHH_144 − Linear model
BMC: 153.9 μM / 307.79μM, p−adjust: 1
PHH_BHT_8hr − SRXN1 (PHH_144) − Power
BMC: 206.22 μM, WTT p−adj: 0.6156, DEG p−adj: 2.40e−01
PHH_BHT_8hr − TXNRD1 (PHH_144) − Exp 2
BMC: 128.5 μM, WTT p−adj: 0.80865, DEG p−adj: 5.97e−01
PHH_BHT_8hr − PHH_144 − NA
BMC: NA μM, BMDL: NA μM, BMDU: NA μM
HepaRG_BHT_8hr − PHH_144 − Hill model
BMC: 9.4 μM / 477.25μM, p−adjust: 0.73877
HepaRG_BHT_8hr − SRXN1 (PHH_144) − Exp 4
BMC: 1.96 μM, WTT p−adj: 0.50938, DEG p−adj: 3.35e−01
HepaRG_BHT_8hr − TXNRD1 (PHH_144) − Hill
BMC: 8.34 μM, WTT p−adj: 0.56579, DEG p−adj: 4.44e−01
HepaRG_BHT_8hr − PHH_144 − Hill
BMC: 9.4 μM, BMDL: 0.56 μM, BMDU: 9e+08 μM
Module EGS
Gene log2 expression
Concentration (μM)
